# PI(3,5)P_2_ Controls the Signaling Activity of Class I PI3K

**DOI:** 10.1101/2023.01.25.525550

**Authors:** Jiachen Sun, Julian Zalejski, Seohyeon Song, Ashutosh Sharma, Wei Wang, Yusi Hu, Wen-Ting Lo, Philipp Alexander Koch, Jagriti Singh, Indira Singaram, Baoshu An, Jean J. Zhao, Liang-Wei Gong, Volker Haucke, Ruixuan Gao, Wonhwa Cho

## Abstract

3-Phosphoinositides are ubiquitous cellular lipids that play pivotal regulatory roles in health and disease. Among 3-phosphoinositides, phosphatidylinositol-3,5-bisphosphate (PI(3,5)P_2_) remains the least understood species in terms of its spatiotemporal dynamics and physiological function due to the lack of a specific sensor that allows spatiotemporally resolved quantitative imaging of PI(3,5)P_2_. Using a newly developed ratiometric PI(3,5)P_2_ sensor engineered from the C-terminal SH2 domain of Class I phosphoinositide 3-kinases (PI3K)-p85α subunit we demonstrate that a unique pool of PI(3,5)P_2_ is generated on lysosomes and late endosomes in response to growth factor stimulation. This PI(3,5)P_2_, the formation of which is mediated sequentially by Class II PI3KC2β and PIKfyve, plays a crucial role in terminating the activity of growth factor-stimulated Class I PI3K, one of the most frequently mutated proteins in cancer, via specific interaction with its regulatory p85 subunit. A small molecule inhibitor of p85α-PI(3,5)P_2_ binding specifically blocks the feedback inhibition of Class I PI3K by PI(3,5)P_2_ and thus serves as a PI3K activator that promotes neurite growth. Furthermore, cancer-causing mutations of the Class I PI3K-p85 subunit inhibit p85-PI(3,5)P_2_ interaction and thereby induce sustained activation of Class I PI3K. Our results unravel a hitherto unknown spatiotemporally specific regulatory function of PI(3,5)P_2_ that links Class I and II PI3Ks and modulates the magnitude of PI3K-mediated growth factor signaling. These results also suggest new therapeutic possibilities for treating cancer patients with p85 mutations and promoting wound healing and tissue regeneration.

## INTRODUCTION

Phosphoinositides (PtdInsPs) are phosphorylated derivatives of phosphatidylinositol that play critical roles in cellular signaling and membrane trafficking ^1, 2, 3^. 3’-PtdInsPs consist of PI(3)P, PI(3,5)P_2_, phosphatidylinositol-3,4-bisphosphate (PI(3,4)P_2_), and phosphatidylinositol-3,4,5-triphosphate (PIP_3_) that are metabolically linked through lipid kinases and phosphatases ^4^. In response to cell activation by growth factors and other stimuli, Class I PI3K converts phosphatidylinositol-4,5-bisphosphate (PI(4,5)P_2_) in the plasma membrane (PM) to PIP_3_, from which PI(3,4)P_2_ is generated by 5’-PtdInsP phosphatases ^5, 6, 7, 8 9^, including SHIP2. PI(3,4)P_2_ can be also produced from phosphatidylinositol-4-phosphate (PI4P) by Class II PI3K ^9, 10, 11, 12^. PIP_3_ and PI(3,4)P_2_ activate diverse downstream signaling pathways, most notably Akt signaling ^5, 6^. PI(3)P, which is generally known to be involved in membrane trafficking, is generated primarily in endosomes from phosphatidylinositol by Class II or Class III PI3K ^13^. It can be also produced from PI(3,4)P_2_ by the 4’-PtdInsP phosphatase INPP4. It has been reported that PI(3,5)P_2_ is synthesized exclusively by PIKfyve from PI(3)P on endolysomal membranes and is involved in membrane trafficking and cell signaling ^9, 14, 15^. PIKfyve and PI(3,5)P_2_ have gained much attention lately as PIKfyve-specific inhibitors, apilimod in particular, have emerged as promising drug candidates for cancer ^16, 17, 18^, neurodegenerative diseases ^9, 14^, and COVID-19 ^19^. Compared to other 3’-PtdInsPs, however, the spatiotemporal cellular dynamics and function of PI(3,5)P_2_ have remained elusive due to the lack of a PI(3,5)P_2_ probe that allows robust PI(3,5)P_2_ quantification^15^. In general, PtdInsPs operate by recruiting effector proteins to defined membrane sites where they regulate the cellular activities of these proteins ^1, 2, 3^. Only a limited number of proteins have been reported to bind PI(3,5)P_2_ with significant specificity and affinity ^14, 20^, which has hampered development of a PI(3,5)P_2_-specific biosensor. To date, two PI(3,5)P_2_ sensors have been reported. First, a tandem dimer of the soluble N-terminus of ion channel, TRPML1 (ML1Nx2) was reported as a PI(3,5)P_2_ biosensor ^21^. However, veracity of this probe was disputed due to its poor PI(3,5)P_2_ selectivity in cells ^22^, underscoring the difficulty in developing a genuine PI(3,5)P_2_-specific biosensor. Another PI(3,5)P_2_ sensor was recently developed from *Dictyostelium* sorting nexin-like protein SnxA ^20^. Importantly, none of these fluorescent protein-based probes allows true spatiotemporally resolved quantification of cellular PI(3,5)P_2_, which is crucial for determining the exact cellular functions of PI(3,5)P_2_ in health and disease.

It has been generally thought that three classes of PI3Ks, which are central players in complex 3’-PtdInsP signaling pathways, have independent functions and regulatory mechanisms ^5, 6^. Although the presence of cross-talks and inter-dependence of their signaling pathways has been speculated, the exact nature of the interplay among three classes of PI3Ks in health and disease is not fully understood. In particular, little is known about the potential involvement of Class II and III PI3Ks and their products, most notably PI(3)P and PI(3,5)P_2_, in regulation of Class I PI3K, the most mutated protein in human breast cancer ^5, 6^. To address these critical mechanistic questions, we developed new PI(3)P and PI(3,5)P_2_-specific ratiometric sensors for *in situ* quantification of PI(3)P and PI(3,5)P_2_, respectively, in living cells. In conjunction with our previously reported ratiometric sensors for PIP_3_ and PI(3,4)P_2_ ^8^, this complete set of 3’-PtdInsP sensors allowed us to quantitatively monitor and analyze the action and cross-talk of three classes of PI3Ks during diverse cellular processes. Notably, spatiotemporally resolved lipid quantification coupled with specific enzyme modulation revealed a hitherto unknown feedback inhibition mechanism in which a specific pool of PI(3,5)P_2_, generated by the sequential action of Class II PI3KC2β and PIKfyve, terminates the growth factor-stimulated Class I PI3K activity by interacting with the Src-homology 2 (SH2) domains of its regulatory p85 subunit. Based on this discovery, we developed a specific small molecule inhibitor of the p85-PI(3,5)P_2_ interaction as a PI3K activator that can potentially promote wound healing and tissue regeneration. We also found that many hot-spot oncogenic mutations in the p85 subunit of Class I PI3K induce sustained activation of Class I PI3K by weakening the inhibitory p85-PI(3,5)P_2_ interaction.

## RESULTS

### The C-terminal SH2 domain of p85 selectively binds PI(3,5)P_2_

To identity an ideal PI(3,5)P_2_-binding protein that can serve as a template for specific PI(3,5)P_2_ sensor development, we first surveyed our database on the lipid binding activity of >500 modular human protein domains, including lipid binding domains ^23^ and protein interaction domains^24, 25^. We then selected all PtdInsP-binding domains and performed rigorous quantitative analysis of their PI(3,5)P_2_ affinity and selectivity by the surface plasmon resonance (SPR) analysis. This comprehensive screening identified the C-terminal Src homology 2 (SH2) domains (cSH2) of the regulatory p85α and p85β subunits of Class I PI3K as PI(3,5)P_2_-binding proteins with the highest affinity and selectivity. The SH2 domain is a prototypal protein interaction domain that specifically recognizes the phosphotyrosine (pY) motif of diverse target molecules ^26, 27, 28^. Through genome-wide SPR-based screening we recently showed that most human SH2 domains tightly bind PtdInsPs ^25^, which is crucial for the enzymatic activity and scaffolding function of their host proteins ^29, 30, 31^. p85α-cSH2 prefers PI(3,5)P_2_ to other PtdInsPs in large unilamellar vesicles (LUVs) (**Fig. 1A**). The N-terminal SH2 domain of p85α (p85α-nSH2) also bound PI(3,5)P_2_ but unlike p85-cSH2 it did not display PI(3,5)P_2_ selectivity (**Supplementary Fig. S1B**). The SPR responses in **Fig. 1A** seem to indicate that p85α-cSH2 also binds PI(4,5)P_2_, the most abundant phosphoinositide, with significant affinity. It should be noted, however, that comparison of SPR responses under one specific condition may not reveal accurate relative lipid affinity^32^. It is because different lipids may induce different membrane binding modes of a protein and hence different response values for the same degree of membrane binding^32^. More rigorous binding analysis with varying protein concentrations (**Supplementary Fig. S1C** and **Table S1**) show that p85α-cSH2 has five-fold higher affinity for PI(3,5)P_2_-containing LUVs than for PI(4,5)P_2_ vesicles. Notably, the PI(3,5)P_2_ affinity of p85α-cSH2 (i.e., *K*_d_ ≂ 200 nM) is comparable to that reported for binding of canonical lipid binding domains to their cognate lipids ^23^. p85β-cSH2 also showed PI(3,5)P_2_ selectivity (**Supplementary Fig. S1A**) but since p85α-cSH2 had ca. 3-times higher membrane affinity than p85β-cSH2 (**Supplementary Fig. S1C** and **Table S1**), we focused our ensuing work on p85α-cSH2.

**Fig. 1.**
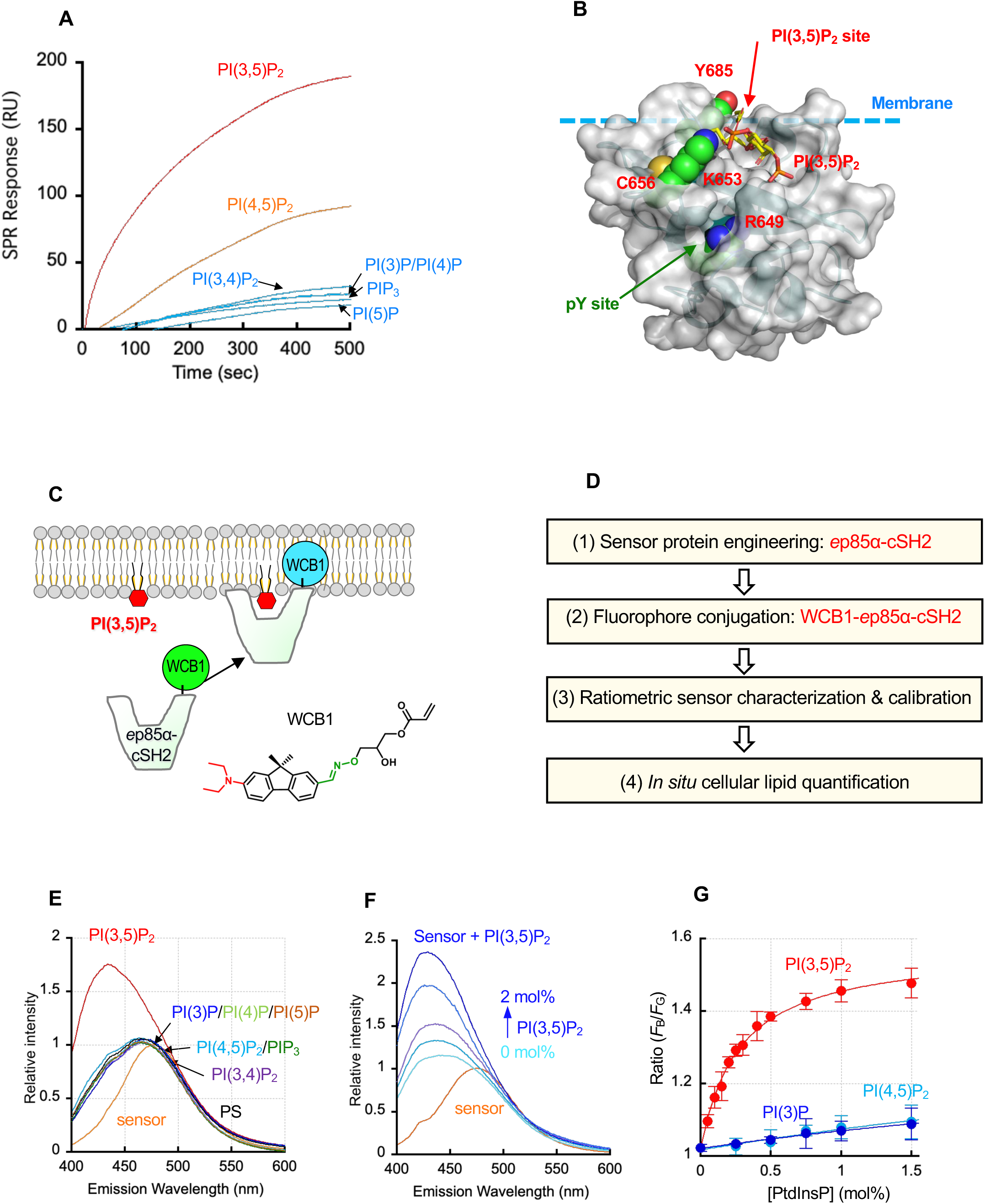
Membrane binding properties of the p85α-cSH2 domain and p85α-cSH2-derived PI(3,5)P_2_ sensor. **A.** PtdInsP selectivity of p85α-cSH2 determined by SPR analysis. POPC/POPS/PtdInsP (77:20:3 in mole ratio) LUVs were coated on the L1 sensor chip and 200 nM p85α-cSH2 was injected into the flow cell for binding measurements. PI(3)P and PI(4)P yielded essentially the same responses. **B.** The structure of p85α-cSH2 (protein data bank ID = 1H9O) shown in a surface diagram with a docked PI(3,5)P_2_ molecule (stick representation). Mutated residues in this study are shown in space-filling representation (carbon in green, oxygen in red, nitrogen in blue and sulfur in orange) and labeled. The molecule is oriented with its potential membrane binding surface facing upward. The dotted cyan line indicates a putative location of the membrane surface. A putative PI(3,5)P_2_ binding site and the reported pY binding pocket are indicated by red and green arrows, respectively. **C.** A basic principle of ratiometric PI(3,5)P_2_ imaging. The ratiometric PI(3,5)P_2_ sensor, WCB1-*e*p85α-cSH2, contains a bright and photostable solvatochromic fluorophore WCB1 ((*E*)-3-((((7-(diethylamino)-9,9-dimethyl-9*H*-fluoren-2-yl)methylene)amino)oxy)-2-hydroxypropyl acrylate) that undergoes a hypsochromic shift (or blue shift) in fluorescence emission upon binding to membranes (see Fig. 1E). **D.** Steps of a ratiometric PI(3,5)P_2_ sensor development and cellular ratiometric imaging. See **Fig. S2A** for detailed procedure for sensor development and characterization. **E.** PtdInsP selectivity of WCB1-*e*p85α-cSH2. Fluorescence emission spectra of WCB1-*e*p85α-cSH2 (500 nM) in response to binding to 10 μM POPC/POPS (80:20) (labeled PS in black) and POPC/POPS/PtdInsP (77:20:3) (each PtdInsP separately labeled and colored) LUVs were obtained spectrofluorometrically with the excitation wavelength set at 380 nm. The spectrum of sensor without lipid is shown in orange at the bottom. **F.** Fluorescence emission spectra of WCB1-*e*p85α-cSH2 in response to binding to POPC/POPS/PI(3,5)P_2_ (80–*x*/20/*x*: *x* = 0–2 mol%) LUVs. The spectra were obtained spectrofluorometrically with the excitation wavelength set at 380 nm. The intensity values were normalized using the maximal intensity of the free sensor as the reference. The emission spectrum of the sensor (orange) was blue-shifted upon vesicle binding and its maximal emission intensity at 440 nm was increased as the PI(3,5)P_2_ concentration varied from 0, 0.5, 1, 1.5, and 2 mol% from bottom to top. **G.** The giant unilamellar vesicle (GUV) binding curve of WCB1-*e*p85α-cSH2 determined by fluorescence microscopy. Lipid compositions of GUVs were POPC/POPS/PtdInsP (80-x/20/x: *x* = 0-2 mol%). PI(3,5)P_2_ (red), PI(4,5)P_2_ (cyan) or PI(3)P (blue) were used. The binding isotherms were analyzed by non-linear least squares analysis using a modified Langmuir equation: *y* = *y*_min_ + (*y*_max_ - *y*_min_)/(1 + *K*_d_ / [PtdInsP]) where *K*_d_, *y*_max_ and *y*_min_ are PtdInsP concentrations ([PtdInsP]) yielding half maximal binding, the maximal and minimal *y* values. Half maximal vesicle binding was achieved with [PI(3,5)P_2_] = 0.27 ± 0.02 mol% and [PI(4,5)P_2_ or PI(3)P] > 10 mol%. The plot for PI(3,5)P_2_ was used as a ratiometric calibration curve for converting experimentally observed *F*_B_/*F*_G_ values to cellular [PI(3,5)P_2_].

Electrostatic and surface cavity analysis ^25^ of p85α-cSH2 predicted that a potential PI(3,5)P_2_ binding site of p85α-cSH2 is formed by a cationic pocket that is topologically distinct from the pY-binding pocket (**Fig. 1B**). Molecular docking analysis of p85α-cSH2-PI(3,5)P_2_ also suggested that the pocket could accommodate a PI(3,5)P_2_ headgroup with the side chain of K653 making energetically favorable interactions with the 3’-phosphate groups of the PI(3,5)P_2_ molecule (**Fig. 1B**). Consistent with this prediction, mutation of K653 to Ala greatly reduced the affinity of p85α-cSH2 for PI(3,5)P_2_ (**Fig. S1D** and **Supplementary Table S1**) and abrogated its selectivity for PI(3,5)P_2_ over PI(4,5)P_2_ (**Fig. S1E**), verifying its crucial role in PI(3,5)P_2_ binding. In contrast, mutation of a conserved cationic residue (R649) in the pY-binding pocket to Ala had no effect on PI(3,5)P_2_ association (**Fig. S1D** and **Supplementary Table S1**). When assayed by fluorescence anisotropy K653A had wild type (WT)-like affinity for the cognate pY-containing peptide of p85α-cSH2 whereas R649A had lower affinity than WT by three orders of magnitude (**Fig. Supplementary S1F**), showing that p85α-cSH2 has non-overlapping binding sites for PI(3,5)P_2_ and pY. Collectively, these studies establish p85α-cSH2 as a *bona fide* PI(3,5)P_2_-binding domain that can be used as a template for PI(3,5)P_2_-specific sensor development. These data also suggest that PI(3,5)P_2_ might play a regulatory role for the PI3K-p85 under physiological conditions by directly interacting with its SH2 domains.

### Development of a PI(3,5)P_2_-specific ratiometric sensor from p85α-cSH2

We recently developed a ratiometric fluorescence sensor-based *in situ* quantitative lipid imaging technology (**Fig. 1C**), which has been successfully applied to the spatiotemporally resolved quantification of diverse cellular lipids, including PI(4,5)P_2_, PIP_3_, PI(3,4)P_2_, phosphatidylserine, and cholesterol, in live cells ^8, 33, 34, 35^. Since it has been estimated that the cellular concentration of PI(3,5)P_2_ is low ^14, 15^, an effective PI(3,5)P_2_ sensor must have high PI(3,5)P_2_ selectivity over more abundant PtdInsP species, PI(4,5)P_2_ in particular, and a linear response range at low PI(3,5)P_2_ levels. To meet these stringent requirements, we generated a PI(3,5)P_2_-specific ratiometric sensor by extensive protein engineering of the p85α-cSH2 (see **Supplementary Fig. S2A** for detailed procedures). Briefly, we generated a large number of multi-site mutants of p85α-cSH2 by saturation and combination mutations of those residues near the putative PI(3,5)P_2_-binding site, including R649, C656 and Y685, to eliminate its binding affinity for pY and other PtdInsPs and to introduce a single Cys on the membrane binding surface for fluorophore conjugation. We then chemically conjugated them with our collection of solvatochromic fluorophores ^36^ and screened them to identify an ideal sensor molecule (**Supplementary Fig. S2B, S2C**). A special emphasis was given to completely suppress PI(4,5)P_2_ binding as p85α-cSH2 has some affinity for PI(4,5)P_2_ (see **Fig. 1A**). Among a large number of candidate sensor molecules, WCB1-*e*p85α-cSH2 (i.e., WCB1-p85α-cSH2-R649A/C656R/Y685C), demonstrated much higher PI(3,5)P_2_ affinity and selectivity than EGFP-p85α-cSH2-WT. When measured spectrofluorometrically, WCB1-*e*p85α-cSH2 showed a large hypsochromic shift (i.e., emission maximum shifted from 490 to 430 nm) upon binding to only PI(3,5)P_2_-containing vesicles (**Fig. 1E**) in a PI(3,5)P_2_ concentration-dependent manner (**Fig. 1F**). It showed essentially no binding to PI(4,5)P_2_-or any other PtdInsP-containing vesicles above the background (i.e., phosphatidylserine-containing vesicles) (**Fig. 1E**). Presumably, this improved specificity and affinity is attributed to the chemically conjugated amphiphilic fluorophore, WCB1, which was reported to affect membrane binding properties of host proteins^33, 36^. Confocal imaging and ratiometric image analysis of binding of WCB1-*e*p85α-cSH2 to PI(3,5)P_2_-containing giant unilamellar vesicles showed that WCB1-*e*p85α-cSH2 has a robust response range of 0-1 mol% of PI(3,5)P_2_ with the concentration for half-maximal binding at 0.27 ± 0.02 mol% of PI(3,5)P_2_ (**Fig. 1G**), indicating that it can successfully cover the low physiological concentration range of PI(3,5)P_2_. Notably, it showed much lower ratiometric responses to giant unilamellar vesicles containing other PtdInsPs, including PI(4,5)P_2_ and PI(3)P, with the concentrations for half-maximal binding > 10 mol% (**Fig. 1G**). Since the average PM concentration of PI(4,5)P_2_, which is the most abundant PtdInsP species, has been reported to be ∼1 mol% ^8, 33^, WCB1-*e*p85α-cSH2 would not respond to other PtdInsPs under physiological conditions. Lastly. WCB1-*e*p85α-cSH2 exhibited drastically suppressed affinity for the cognate pY motif of p85α-cSH2 (**Supplementary Fig. S1F**). Collectively, these results demonstrate that our sensor engineering effectively eliminated modest PI(4,5)P_2_ affinity and high pY motif affinity of p85α-cSH2 WT and that WCB1-*e*p85α-cSH2 meets all requirements for a cellular ratiometric sensor of PI(3,5)P_2_ and is hence well suited for robust, sensitive, and specific ratiometric quantification of cellular PI(3,5)P_2_.

### Spatiotemporally resolved quantification of cellular PI(3,5)P_2_

To test the cellular specificity and efficiency of WCB1-*e*p85α-cSH2, we first microinjected it into unstimulated HeLa and HEK293 cells, respectively. Microinjected WCB1-*e*p85α-cSH2 only detects PI(3,5)P_2_ in the cytoplasmic leaflets of organelle membranes because our ratiometric sensors cannot cross the lipid bilayer ^8, 33, 34, 35^. As shown in **Fig. 2A**, microinjected WCB1-*e*p85α-cSH2 displayed a punctate intracellular distribution. Colocalization analysis with various organelle markers showed that WCB1-*e*p85α-cSH2 was primarily localized at lysosomes and also significantly at late endosomes (**Supplementary Fig. S3A**). No statistically significant localization was detected at early endosomes and, more importantly, PM (**Supplementary Fig. S3A**). It has been reported that the PM contains the highest concentration of PI(4,5)P_2_ (i.e., 1 mol%) ^33, 34^ and also significant levels of PI(4)P ^37^ and PI(5)P ^15^. Thus, the complete lack of sensor signal at the PM demonstrates that WCB1-*e*p85α-cSH2 does not respond to other PtdInsPs, PI(4,5)P_2_ in particular. To preclude any possibility that WCB1-*e*p85α-cSH2 responds to PI(4,5)P_2_, we performed an additional control experiment. Specifically, we increased the PI(4,5)P_2_ level in the PM by overexpressing PI(4)P 5-kinase type 1 β (PIP5K1B), which converts PI(4)P into PI(4,5)P_2_ in the PM ^37^, and quantitatively monitored the responses by WCB1-*e*p85α-cSH2 and a ratiometric PI(4,5)P_2_ sensor, DAN-*e*ENTH ^33^, respectively. Consistent with the previous report ^38^, overexpression of PIP5K1B produced about a 30% increased level of PI(4,5)P_2_ in the PM, as detected by microinjected DAN-*e*ENTH (**Supplementary Fig. S4A**). Under the same conditions, however, microinjected WCB1-*e*p85α-cSH2 did not show any PM localization with and without PIP5K1B overexpression (**Supplementary Fig. S4B**), confirming that WCB1-*e*p85α-cSH2 does not respond to cellular PI(4,5)P_2_.

**Fig. 2.**
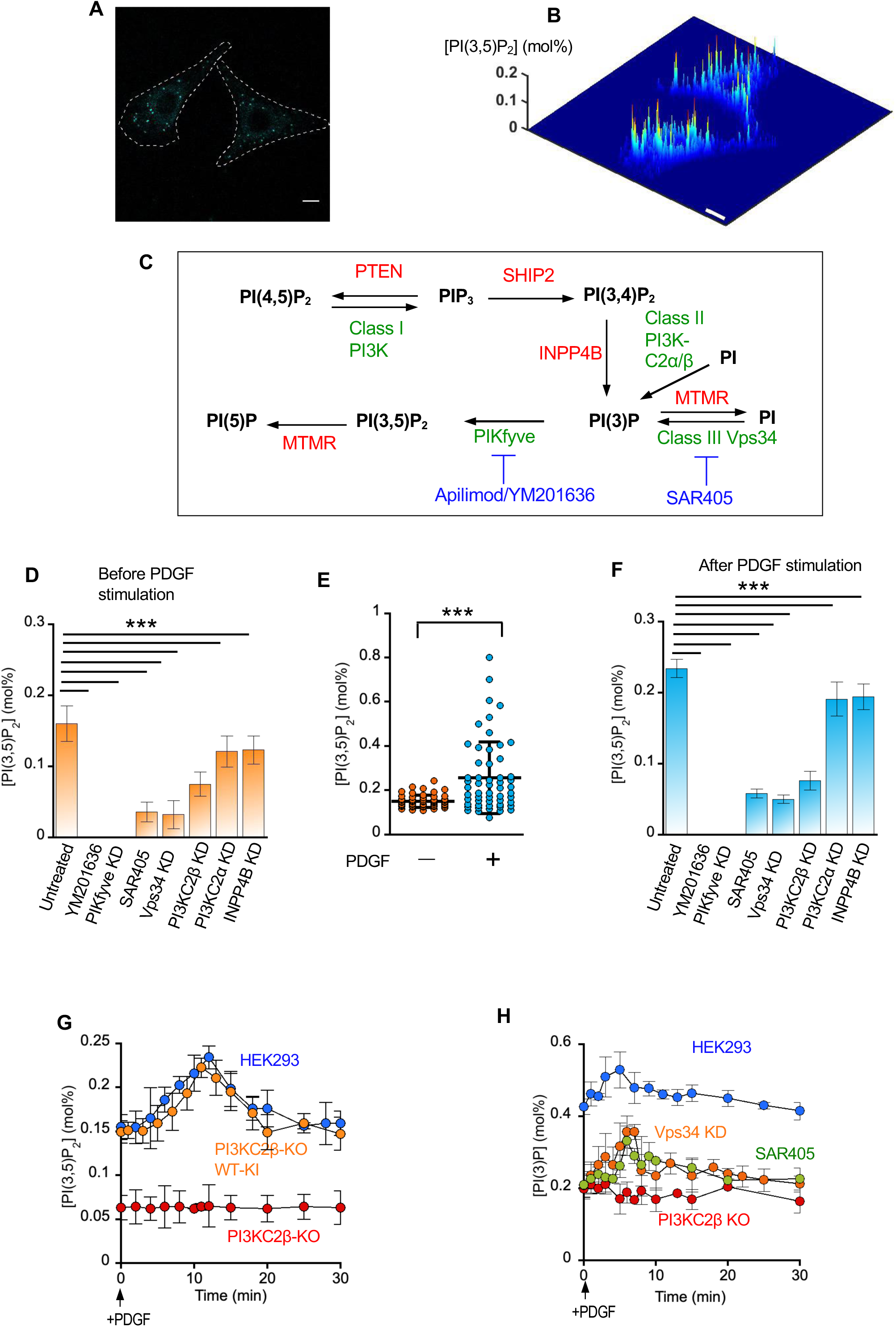
Quantitative cellular imaging of PI(3)P and PI(3,5)P_2_. **A.** Subcellular distribution of microinjected WCB1-*e*p85α-cSH2 in HeLa cells. The blue channel image shows intracellular punctate distribution of the sensor. The scale bar and dotted lines show 10 μm and cell outlines, respectively. The same pattern was observed in HEK293 cells. **B.** A spatially resolved PI(3,5)P_2_ profile calculated from the two-channel cross-sectional ratiometric image of a representative HeLa cell at steady-state. The *z*-axis scale indicates the PI(3,5)P_2_ concentration ([PI(3,5)P_2_]) (mol%). A pseudo-coloring scheme with red and blue representing the highest (0.2 mol%) and the lowest (0 mol%) concentration, respectively, is used to illustrate the spatial PI(3,5)P_2_ heterogeneity. Scale bars in **A** and **B** indicate 10 μm. Essentially the same results were obtained in HEK293 cells. **C.** Cellular metabolism of PI(3,5)P_2_. Most of (not exhaustive) lipid kinases and phosphatase involved in the metabolism are shown in green and red, respectively. Available small-molecule inhibitors are indicated in blue. **D.** The effects of inhibiting lipid kinases and phosphatases on spatially averaged [PI(3,5)P_2_] in HEK293 cells before PDGF stimulation. Cells were pretreated with a PIKFyve inhibitor, apilimod (1 μM, 1 h) or YM201636 (0.8 μM, 1 h), siRNAs (KD) for PIKfyve, a Vps34 inhibitor SAR405 (10 μM, 24 h), and siRNAs for Vps34, PI3KC2α, PI3KC2β, and INPP4B, respectively. Apilimod and YM201636 yielded the same results. All siRNA treatments were for 48 h. *p* < 0.0001 for all data set (e.g., untreated versus SAR405). **E.** An increase in spatially averaged [PI(3,5)P_2_] in HEK293 cells at 12 min post PDGF stimulation (50 ng/ml). *p* < 0.0001. **F.** The effects of inhibiting lipid kinases and phosphatases on spatially averaged [PI(3,5)P_2_] in HEK293 cells 12 min after PDGF stimulation (50 ng/ml). See Fig. 2D for conditions. Apilimod and YM201636 yielded the same results. *p* < 0.0001 for all data set (e.g., untreated versus SAR405). **G.** Time-dependent changes of the spatially averaged [PI(3,5)P_2_] in HEK293 (blue) and PI3KC2β-null HEK293 (KO) (red), and PI3KC2β-null HEK293 cells after reintroduction of PI3KC2β WT (KO + WT-KI) (orange) after PDGF stimulation (50 ng/ml). [PI(3,5)P_2_] values were determined at different time intervals. **H.** Time-dependent changes of the spatially averaged PI(3)P concentration ([PI(3)P]) (*n* = 30-35) in HEK293 cells after PDGF stimulation (50 ng/ml). 10 μM SAR405 (24 h) was used for Vps34 inhibition. For all steady-state PI(3,5)P_2_ quantification measurements, *n* = 5 with >10 cells per experiment. Averages and S.D.’s are shown for each dataset. For all time-dependent PI(3,5)P_2_ quantification, *n* = 3 with 5 cells per experiment. Each data represents an average and S.D. at a given time point.

Ratiometric quantification of cross-sectional images illustrated the spatial distribution of PI(3,5)P_2_ at the cytoplasmic leaflets of lysosomes and late endosomes at a given time (**Fig. 2B**). The spatially averaged PI(3,5)P_2_ concentration was 0.15 ± 0.03 mol% in HeLa cells. Essentially the same results were obtained in HEK293 cells. This steady-state PI(3,5)P_2_ concentration in unstimulated mammalian cells is significantly lower than that reported for PI(4,5)P_2_ ^33, 34^ and comparable to those reported for PIP_3_ and PI(3,4)P_2_ ^8^. To validate that WCB1-*e*p85α-cSH2 quantitatively monitor cellular PI(3,5)P_2_, we measured the effects of suppressing the activity of various lipid kinases and phosphatases on the PI(3,5)P_2_ level in HEK293 cells (**Fig. 2C**). First, we inhibited PIKfyve, which converts PI(3)P into PI(3,5)P_2_, by either apilimod or YM201636. The treatment reduced the concentration of PI(3,5)P_2_ to undetectable levels (**Fig. 2D** and **Supplementary Fig. S3B and S3C**), verifying that WCB1-*e*p85α-cSH2 specifically recognizes cellular PI(3,5)P_2_. siRNA-mediated suppression of PIKfyve (**Fig S5A**) yielded the same results (**Fig. 2D** and **Fig. S3D**). We also measured the effects of inhibiting PI(3)P-producing enzymes, PI3KC2α, PI3KC2β, Vps34, and INPP4B, on PI(3,5)P_2_ levels. When Vps34 was pharmacologically inhibited by SAR405 (**Fig. S3E**) or genetically suppressed by siRNA (**Supplementary Fig. S5B** and **Fig. S3F**), the spatially averaged PI(3,5)P_2_ concentration at late endosomes and lysosomes was decreased to 0.05 mol% (**Fig. 2D**), suggesting that Vps34 is a major supplier of PI(3)P molecules that are converted to PI(3,5)P_2_ at steady-state. Interestingly, siRNA-mediated suppression of PI3KC2β expression (**Fig. S5C**) also significantly reduced PI(3,5)P_2_ levels (**Fig. 2D** and **Supplementary Fig. S3G**), suggesting its involvement in the production of PI(3,5)P_2_ at steady-state. siRNA-mediated suppression of INPP4B and PI3KC2α (**Supplementary Fig. S5D, S5E**) displayed much smaller effects on PI(3,5)P_2_ levels in HEK293 cells than that of Vps34 and PI3KC2β (**Fig. 2D, Supplementary Fig. S3H** and **S3I**).

### Growth factor stimulation produces a special pool of PI(3,5)P_2_ via PI3KC2β

It has been reported that PI(3)P is produced by Class II PI3K in response to growth factor stimulation ^39, 40^. Since growth factor-generated PI(3)P can be converted into PI(3,5)P_2_, we quantified PI(3,5)P_2_ in response to stimulation of HEK293 cells by various growth factors. When HEK293 cells were treated with platelet-derived growth factor (PDGF), the spatially averaged PI(3,5)P_2_ concentration was increased from 0.15 to 0.23 mol% after 15 min, i.e., by >50% (**Fig. 2E**), indicating that additional PI(3,5)P_2_ molecules were generated upon PDGF stimulation. Although the average value was 0.23 mol%, the local PI(3,5)P_2_ concentration on individual late endosome/lysosomes varied widely, with some punctae containing up to 0.8 mol% PI(3,5)P_2_ (**Fig. 2E)**. Again, pharmacological inhibition or siRNA-mediated suppression of PIKfyve abrogated PI(3,5)P_2_ formation (**Fig. 2F**). Also, pharmacological inhibition or knockdown of Vps34 or PI3KC2β knockdown greatly reduced the level of PDGF-induced PI(3,5)P_2_, whereas PI3KC2α and INPP4B knockdown had much smaller effects (**Fig. 2F**). Notably, although suppression of both Vps34 and PI3KC2β greatly reduced overall PI(3,5)P_2_ levels before and after PDGF stimulation, only PI3KC2β-depleted HEK293 cells showed no increase in PI(3,5)P_2_ upon PDGF stimulation (compare the sixth columns in **Fig. 2D** and **2F**). These results suggest that there may be two distinct pools of cellular PI(3,5)P_2_, a steady-state form and a PDGF-induced form and that Class II PI3KC2β may play an important role in producing the latter.

To test the notion of two separate PI(3,5)P_2_ pools, we quantified PI(3,5)P_2_ in HEK293 cells after PDGF stimulation in a spatiotemporally resolved manner. From its basal level of 0.15 mol%, PI(3,5)P_2_ reached 0.23 mol% at 12 min post-PDGF stimulation, then declined to basal levels after 20 min (**Fig. 2G**). Importantly, gene ablation (**Fig. 2G** and **Supplementary Fig S5C**) or knockdown (**Supplementary Fig. S5A** and **Fig S5C**) of the PI3KC2β not only lowered the basal PI(3,5)P_2_ level but also completely eliminated the PDGF-stimulated PI(3,5)P_2_ increase. Furthermore, PDGF-induced PI(3,5)P_2_ production was fully restored when PI3KC2β WT was re-introduced into PI3KC2β-null HEK293 cells (**Fig. 2G** and **Supplementary Fig S5C**), confirming the essential role of PI3KC2β in this process. In contrast to PI3KC2β suppression, pharmacological inhibition (i.e., SAR405) or knockdown of Vps34 lowered the basal level of PI(3,5)P_2_ to 0.04 mol% without affecting the PDGF-induced PI(3,5)P_2_ increase (**Supplementary Fig. S6A**). Moreover, knockdown of PI3KC2α or INPP4B had only small effects on the basal PI(3,5)P_2_ level and little to no effect on PDGF-induced PI(3,5)P_2_ synthesis (**Supplementary Fig. S6A**). Lastly, pharmacological inhibition (YM201636) of PIKfyve, the enzyme that converts PI(3)P to PI(3,5)P_2_, completely abolished PI(3,5)P_2_ throughout the time course of PDGF stimulation, confirming that PIKfyve is the only enzyme that produces PI(3,5)P_2_ (**Supplementary Fig. S6A**).

These observations were not limited to PDGF activation of HEK293 cells as similar results were obtained after stimulation of different cells with diverse growth factors (**Supplementary Fig. S6B**). For example, stimulation of HEK293 cells with epidermal growth factor (EGF) and insulin-like growth factor1 (IGF1), respectively, resulted in similar spatiotemporal changes in PI(3,5)P_2_. Activation of NIH 3T3 cells by PDGF and HeLa cells by IGF1, respectively, also yielded similar results. Furthermore, PI3KC2β knockdown abrogated EGF-induced PI(3,5)P_2_ formation in HEK293 cells, IGF1-induced PI(3,5)P_2_ formation in HeLa cells, and PDGF-induced PI(3,5)P_2_ production in NIH 3T3 cells (**Supplementary Fig. S6B**). Taken together, these results support the notion that growth factor-induced, PI3KC2β-mediated production of PI(3,5)P_2_ is a universal phenomenon in both non-cancerous and cancer-derived cell lines.

### Growth factor stimulation produces a special pool of PI(3)P via PI3KC2β

To determine how PI3KC2β contributes to the PDGF-induced PI(3,5)P_2_ increase, we quantified the cellular PI(3)P level. For this purpose, we generated a ratiometric PI(3)P sensor, DAN-*e*EEA1, from the highly PI(3)P-specific FYVE domain of EEA1 (see **Supplementary Fig. S7A** for details). Tandem FYVE domains have been commonly used for fluorescence protein-based genetically encoded PI(3)P probes because a single FYVE domain typically does not have high enough membrane affinity ^41^. However, they are not ideal for ratiometric sensor development due to technical difficulties involved in single-site fluorophore conjugation conducive to a favorable hypsochromic shift. Thus, we engineered a single EEA1-FYVE domain and incorporated 6-acrylo-2-dimethylaminonaphthalene (DAN) group^33^ into an ideal location to both enhance membrane affinity and induce a large hypsochromic shift upon membrane binding (**Supplementary Fig. S7A**). The amphiphilic DAN group is known to enhance the membrane affinity of a lipid binding domain when covalently attached to a Cys in its membrane binding surface ^33^. The resulting PI(3)P sensor, DAN-*e*EEA1 showed a large hypsochromic shift in response only to PI(3)P binding (**Supplementary Fig. S7B**) and had high enough PI(3)P affinity to give a robust linear response to the physiological level of PI(3)P (i.e., 0-1.5 mol%) (**Supplementary Fig. S7C**). We then microinjected DAN-*e*EEA1 into HEK293 cells, stimulated them with PDGF, and performed *in situ* quantifications of PI(3)P at early endosomes, where a large majority of PI(3)P is known to reside ^13^. The spatially averaged PI(3)P concentration was 0.41 ± 0.04 mol% in unstimulated HEK293 cells (**Supplementary Fig. S7D**), which rose to 0.52 ± 0.04 mol% at 6 min post-PDGF stimulation and gradually declined to the basal level (**Fig. 2H**). Pharmacological inhibition or knockdown of Vps34 greatly reduced the basal level of PI(3)P to 0.2 mol% but did not eliminate the PDGF-induced PI(3)P spike (**Fig. 2H**). These data indicate that Vps34 is responsible primarily for producing the steady-state pool of PI(3)P but it is dispensable for the generation of PDGF-induced PI(3)P. Importantly, gene ablation (**Fig. 2H**) or suppression (**Supplementary Fig. S7E**) of PI3KC2β not only reduced the basal PI(3)P level but also eliminated the PDGF-induced PI(3)P peak, indicating that the latter derives primarily from phosphorylation of phosphatidylinositol by PI3KC2β. A combination of Vps34 inhibition and PI3KC2β knockdown essentially abolished all PI(3)P signals (**Supplementary Fig. S7E**). siRNA knockdown of PI3KC2α and INPP4B had little to no effect on the PDGF-induced PI(3)P spike while modestly reducing the basal PI(3)P level (**Supplementary Fig. S7E**). Lastly, siRNA knockdown of PIKfyve enhanced the total PI(3)P concentration, as expected from its role in converting PI(3)P to PI(3,5)P_2_, but did not affect the kinetics of induced PI(3)P formation (**Supplementary Fig. S7E**).

### PI(3,5)P_2_ contributes to termination of growth factor-induced Class I PI3K activity

The mechanism of activation of Class I PI3K has been studied in detail ^42^. In the resting state, the catalytic p110 subunit is autoinhibited by the regulatory p85 subunit. Upon cell stimulation two SH2 domains in p85 bind to pY’s in an activating protein, such as a receptor tyrosine kinase or an adaptor protein, which relieves p110 from inhibitory tethering by p85 ^42^ (see also **Fig. 7B**) Although it is generally thought that Class I PI3K signaling is terminated through the action of lipid phosphatases, including PTEN and INPP4B ^5, 6^, which remove PIP_3_ and PI(3,4)P_2_, respectively, little is known about the mechanism that inactivates Class I PI3K itself post-stimulation. We recently reported ^8^ that the growth factor-induced formation of PIP_3_, which directly reflects the cellular Class I PI3K activity, was limited to the PM and that the PIP_3_ concentration at the PM increases and decreases rapidly (i.e., within 10 min post stimulation) (see also **Fig. 3A**). Genetic ablation of PTEN, which converts PIP_3_ to PI(4,5)P_2_ and is thus generally thought to counteract the Class I PI3K, had little effect on the kinetics of PIP_3_ decrease after PDGF stimulation although it enhanced the amplitude of the PIP_3_ peak (**Supplementary Fig. S8A**). Furthermore, pharmacological inhibition by AS1949490 of SHIP2, which is mainly responsible for conversion of PIP_3_ to PI(3,4)P_2_ in PDGF-stimulated cells ^8^, only slightly slowed down the PIP_3_ decrease (**Supplementary Fig. S8A**). These results suggest the presence of a hitherto unknown mechanism for termination of growth factor-stimulated Class I PI3K activity.

**Fig. 3.**
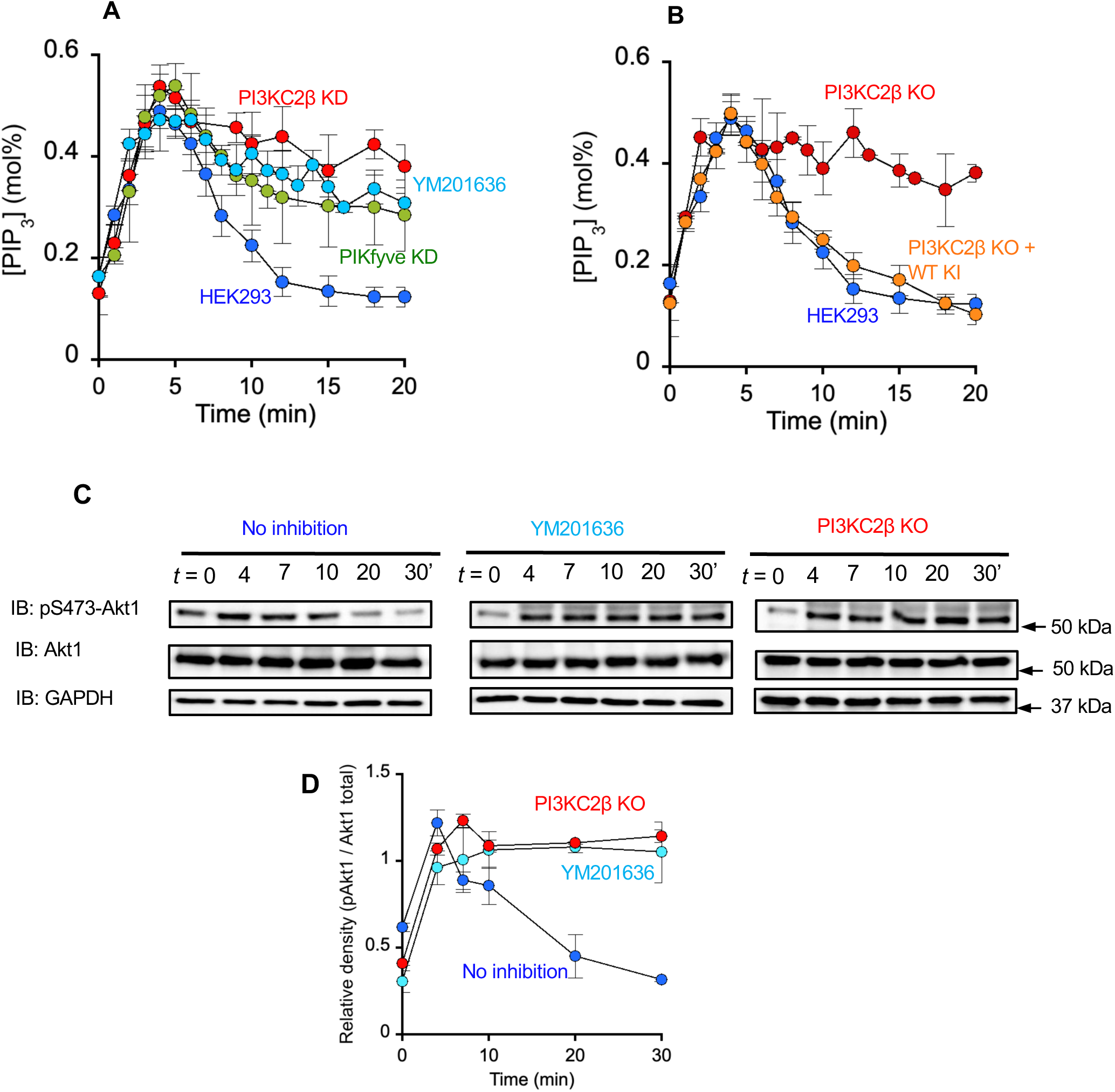
Time courses of PIP_3_ formation and disappearance at the plasma membrane. **A.** PDGF (50 ng/ml)-stimulated changes in the spatially averaged PIP_3_ concentration ([PIP_3_]) at the PM of HEK293 cells. HEK293 cells were either untreated (blue) or pre-treated with 0.8 μM (1 h) YM201636 (cyan), PIKfyve siRNA (KD) and PI3KC2β siRNA (red), respectively. **B.** PDGF-stimulated changes in [PIP_3_] at the PM of HEK293 cells (blue) and PI3KC2β-null (KO) HEK293 cells before (red) and after adding back (KI) PI3KC2β WT (orange). **C**. The time course of PDGF-stimulated phosphorylation of S473 of endogenous Akt1 in HEK293 cells before (left) and after (middle) YM201636 treatment (0.8 μM, 1 h) and in PI3KC2β-null HEK293 cells (right). Representative western blots (*n* = 3) show pS473-Akt1and total Akt1 detected by immunoblot (IB) with the pS473-Akt1-specific and Akt1-specific antibodies, respectively. Glyceraldehyde 3-phosphate dehydrogenase (GAPDH) was used as a loading control. **D.** Quantification of Fig. 3C. The ratio of the gel density of the pS473-Akt1 band to that of the total Akt band is plotted as a function of time. All siRNA treatments were for 48 h. For all time-dependent PIP_3_ quantification, *n* = 3 with 5 cells per experiment. Each data represents an average and S.D. at a given time point.

Our finding that PI(3,5)P_2_ directly binds to both p85-cSH2 (see **Fig. 1A, Supplementary Fig S1C** and **Supplementary Table S1**) and p85-nSH2 (see **Supplementary Fig S1B** and **Supplementary Table S1**) implies that PI(3,5)P_2_ might be involved in regulating PI3K activity. To test this notion, we performed *in situ* quantification of PDGF-induced PIP_3_ in the PM of HEK293 cells using our recently developed ratiometric PIP_3_ sensor ^8^ while modulating the activity of lipid kinases and phosphatases involved in PI(3,5)P_2_ metabolism. Consistent with our previous results ^8^, PDGF stimulation raised the PIP_3_ concentration from 0.15 to 0.50 mol% within 4 min, which was followed by a rapid decrease of PIP_3_ to basal levels within 10 min (**Fig. 3A**). Interestingly, the kinetics of PIP_3_ decrease was synchronized with the kinetics of PI(3,5)P_2_ formation (see **Fig. 2G**): i.e., decrease of PIP_3_ coincided with an increase of PI(3,5)P_2_. Furthermore, pharmacological inhibition or knockdown of PIKFyve, which essentially removed all cellular PI(3,5)P_2_ (see **Supplementary Fig. S6A**), greatly slowed the PIP_3_ decline phase (**Fig. 3A**). Even at 20 min post PDGF stimulation, the PIP_3_ level remained at about 0.3 mol%, which was about 60% of its peak concentration, indicating that Class I PI3K activation was sustained to a large extent. This effect was PI(3,5)P_2_-specific because other treatments that exerted minimal effects on PI(3,5)P_2_ levels (i.e., PI3KC2α and INPP4B knockdown; see **Fig. 2E** and **Supplementary Fig. S6A**) did not alter the kinetics of the PIP_3_ in either the rise or the decline phase (**Supplementary Fig. S8B**). Furthermore, pharmacological inhibition or knockdown of Vps34, which reduced steady-state PI(3)P and PI(3,5)P_2_, but not the PDGF-induced rise in PI(3)P and PI(3,5)P_2_ (see **Supplementary Fig. S6A**), had no effect on PIP_3_ kinetics (**Fig. Supplementary S8B**). These results indicate that the PDGF-induced pool of PI(3,5)P_2_ inhibits Class I PI3K activity. Importantly, this idea was corroborated by the finding that PIP_3_ levels reached a peak and did not appreciably decline for >30 min in PI3KC2β-suppressed (**Fig. 3A**) or PI3KC2β-null HEK293 cells (**Fig. 3B**). Furthermore, re-introduction of PI3KC2β WT to the PI3KC2β-null HEK293 cells fully restored the original PIP_3_ kinetics of WT HEK293 cells (**Fig. 3B**). Similarly, PI3KC2β knockout slowed the PIP_3_ decrease in IGF1-stimulated HEK293 cells (**Supplementary Fig. S8C**) and PIKfyve knockdown slowed the PIP_3_ decrease in EGF-stimulated HeLa cells (**Supplementary Fig. S8D**), supporting the universal nature of the effect of PI(3,5)P_2_ on Class I PI3K activity. Collectively, these results suggest that PI3KC2β/PIKfyve-generated PI(3,5)P_2_ may play a crucial role of in feedback inhibition and termination of growth factor-stimulated Class I PI3K signaling activity.

To test the functional significance of PI(3,5)P-mediated feedback inhibition of Class I PI3K, we measured the cellular activity of endogenous Akt1 that is activated by PIP_3_. S474 phosphorylation (pS474) of Akt1 by mTORC2 in HEK293 cells was reversed within 20 min after PDGF stimulation but remained largely unchanged after 20 min when cells were pre-treated with a PIKFyve inhibitor, YM201636 (**Fig. 3C** and **3D**). Akt1 also remained in its active phosphorylated state for an extended period of time in PI3KC2β-null HEK293 cells (**Fig. 3C** and **3D**).

### PI(3,5)P_2_ inhibits Class I PI3K by displacing it from PI(4,5)P_2_-containing membranes

To test if the observed inhibition of Class I PI3K by PI(3,5)P_2_ is caused by direct PI(3,5)P_2_-p85 interaction and to understand how PI(3,5)P_2_ that is mainly found in endolysosomes can inhibit the action of Class I PI3K at PM, we performed cell-free enzyme kinetic and membrane binding studies of purified Class I PI3Kα (equimolecular p110α + p85α) using model membranes with defined lipid compositions. For enzyme kinetics of PI3Kα, we employed a recently developed *in vitro* activity assay that allows direct *real-time* monitoring of the reaction and accurate quantitative analysis of kinetic parameters ^43^. The assay system consists of POPC/POPS/PI(4,5)P_2_ (77:20:3) LUVs, a ratiometric PI(4,5)P_2_ sensor (DAN-*e*ENTH), PI3Kα, a pY-containing peptide derived from PDGFβ (pY2), and ATP. The reaction was triggered by adding ATP to the reaction mixture containing all other elements and the reaction progress was directly and continuously monitored spectrofluorometrically by following the time course of the ratio of fluorescence emission intensity at two maximal wavelengths before and after the hypsochromic shift, respectively^43^. We reasoned that for PI(3,5)P_2_, which is primarily synthesized within the endolysosomal system, to inhibit PI3Kα at the PM, it should be able to inhibit the action of PI3Kα in a *trans*-vesicular mode (**Fig. 4A**). **Fig. 4B** shows the typical kinetic curves of PI(4,5)P_2_ phosphorylation by PI3Kα, which could be analyzed using a simple Michaelis-Menten kinetic equation ^43^. We previously reported that PI3Kβ was competitively inhibited by its reaction product, PIP_3_ in a *cis* mode ^43^. Consistent with this finding, PI3Kα was also inhibited by pre-added PIP_3_ in the PI(4,5)P_2_-containing vesicles (i.e., POPC/POPS/PI(4,5)P_2_/PIP_3_ (77-*x*/20/3/*x*; *x* = 0-3 mole%)) in a concentration-dependent manner (**Supplementary Fig. S9A**). Under the same conditions, PI(3,5)P_2_ had no effect on PI3Kα activity: i.e., the initial rate was unaltered when the PI(3,5)P_2_ composition was varied in POPC/POPS/PI(4,5)P_2_/ PI(3,5)P_2_ (77-*x*/20/3/*x*; *x* = 0-3 mole%) LUVs (**Supplementary Fig. S9B**). This indicates that PI(3,5)P_2_ neither competes with PI(4,5)P_2_ for the catalytic site in PI3Kα nor causes allosteric inhibition of PI3Kα in a *cis* mode. Importantly, however, when separate POPC/POPS/PI(3,5)P_2_ (77:20:3) LUVs were added to the reaction mixture containing POPC/POPS/PI(4,5)P_2_ (77:20:3) LUVs, the rate of reaction was significantly reduced (**Fig. 4B**). Systematic inhibition analysis using equimolar mixtures of POPC/POPS/PI(4,5)P_2_ (77:20:3) and POPC/POPS/PI(3,5)P_2_ (80-*x*/20/*x*; *x* = 0-3 mole%) LUVs showed that PI(3,5)P_2_ could potently inhibit the catalytic action of PI3Kα in a dose-dependent manner (**Fig. 4B** and **4D**) in a *trans* mode; i.e. by displacing it from PI(4,5)P_2_-containing vesicles. Equimolar PI(3,5)P_2_ in the trans LUVs caused ∼70% inhibition of PI3Kα. When LUVs containing other PtdInsPs (e.g., POPC/POPS/PIP_3_ (77:20:3) or POPC/POPS/PI(3)P (77:20:3)) were added to the reaction mixture, none caused appreciable inhibition of PI3Kα in a *trans* mode (**Supplementary Fig. S9C** and **S9D**). This demonstrates the specificity of *trans*-vesicular PI(3,5)P_2_ inhibition. To verify that the observed inhibition is specifically mediated by p85-cSH2-PI(3,5)P_2_ interactions, we employed a PI3Kα mutant whose p85α subunit harbors the K653A mutation (i.e., p110α + p85α-K653A) that specifically reduces PI(3,5)P_2_ binding of the p85-cSH2 (see **Supplementary Fig. S1D** and **Table S1**). This variant was as active as PI3Kα WT in the absence of PI(3,5)P_2_ but was not inhibited by externally added POPC/POPS/PI(3,5)P_2_ (77-*x*/20/*x*; *x* = 0-3 mole%) LUVs **(Fig. 4C** and **4D)**. These cell-free kinetic studies show that PI(3,5)P_2_ in one membrane can potently and specifically inhibit Class I PI3Kα acting on the other spatially separated membrane presumably by displacing the p110α-p85α heterodimer from the membrane.

**Fig. 4.**
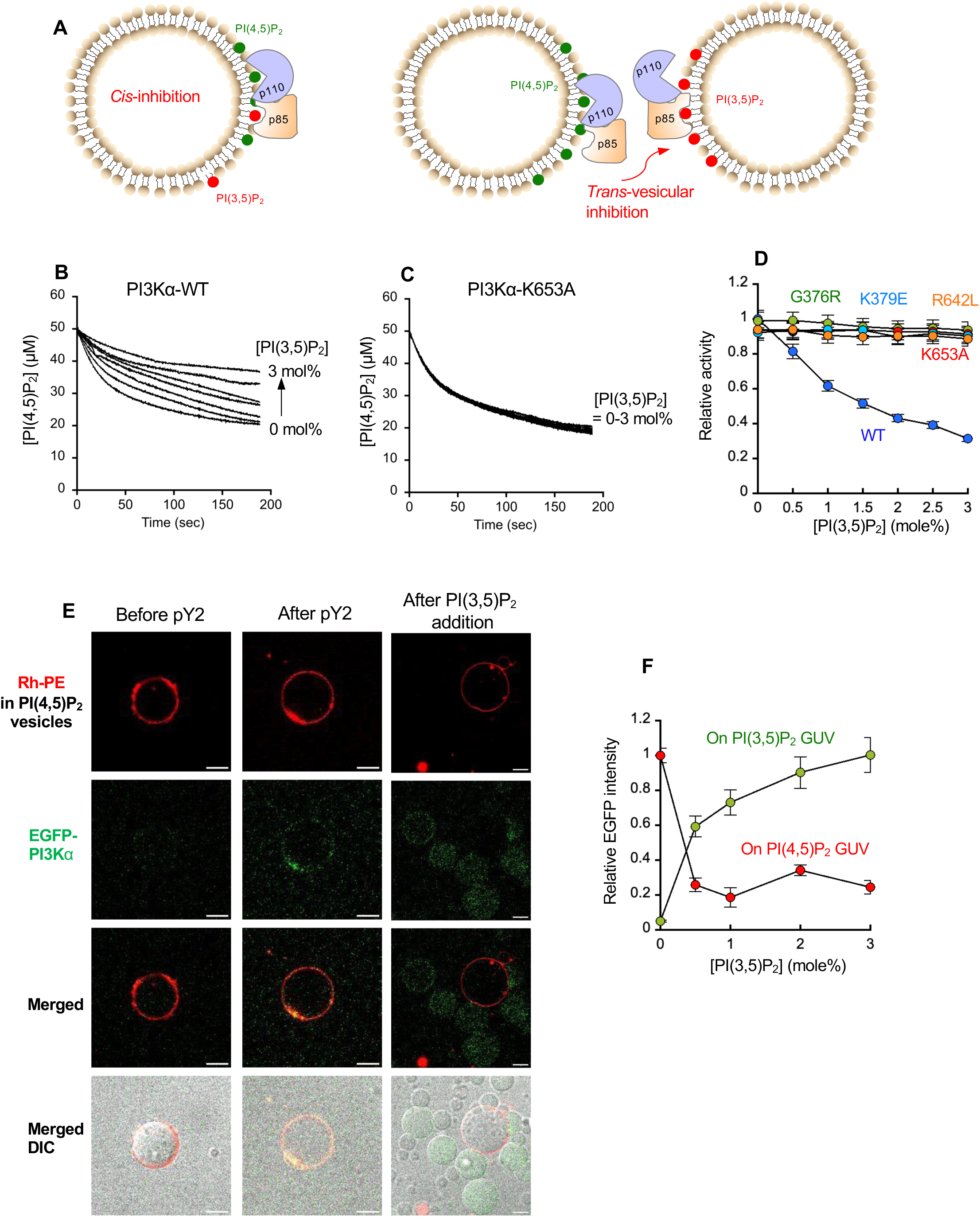
Inhibition of the enzyme activity and plasma membrane binding of PI3Kα by PI(3,5)P_2_. **A.** *cis*-versus *trans*-vesicular inhibition of Class I PI3K by PI(3,5)P_2_. For *cis*-vesicular inhibition, PI(3,5)P_2_ is added to the same vesicles where PI(4,5)P_2_ is present whereas for *trans*-vesicular inhibition, PI(3,5)P_2_-containing vesicles were mixed with PI(4,5)P_2_-containing vesicles. *Cis* inhibition would indicate that PI(3,5)P either competes with PI(4,5)P_2_ for the catalytic sites of PI3K or causes allosteric inhibition. Conversely, *trans* inhibition would support the PI3K-displacement model. **B.** The kinetics of PI3Kα-catalyzed phosphorylation of PI(4,5)P_2_ in POPC/POPS/PI(4,5)P_2_ (77:20:3) LUVs in the presence of POPC/POPS/PI(3,5)P_2_ (80-*x*/20/*x*; *x* = 0, 0.5, 1, 1.5, 2, 2.5 and 3 mol%) LUVs. DAN-*e*ENTH (500 nM) was incubated with 50 μM each of the two LUVs for 1 min, and then 50 nM PI3Kα (p110α-p85α-WT), 10 μM pY2 peptide, and 100 μM ATP were added to initiate the reaction. **C.** The same measurement was performed with the p110α-p85α-K653A mutant. **D.** Concentration-dependent *trans*-vesicular inhibition of PI3Kα WT and mutants by PI(3,5)P_2_. Relative enzyme activity was determined by calculating initial rates of PI(4,5)P_2_ phosphorylation by PI3Kα WT (blue), K653A (red), G376R (green), K379E (cyan), and R642L (orange) from Fig. 4B**-C**, and **Fig S12A-C** and normalized them against that of PI3Kα WT in the absence of PI(3,5)P_2_ (= 0.75 μM/s). Each data point represents average ± S.D. from three independent measurements. **E.** Direct visualization of *trans*-vesicular movement of activated EGFP-PI3Kα (green) from POPC/POPS/PI(4,5)P_2_/1,2-dioleoyl-sn-glycero-3-phosphoethanolamine-N-(lissamine rhodamine B sulfonyl) (Rh-PE) (67:20:3:10) GUVs (red) to POPC/POPS/PI(3,5)P_2_ (80-*x*:20*x*, *x* = 0-3) GUVs (no color) by fluorescence microscopic imaging. The orange color in merged images indicates colocalization of EGFP-PI3Kα and PI(4,5)P_2_ GUVs. Differential interference contrast (DIC) images of GUVs are shown in the bottom row to better illustrate relative locations of GUVs and EGFP-PI3Kα. These images are representative of similar images collected in three independent measurements (>30 GUVs per measurement). Scale bars indicate 10 μm. **F.** Quantification of Fig. 4E. Relative EGFP intensity on PI(4,5)P_2_ (or PI(3,5)P_2_) GUVs was calculated by normalizing the total EGFP fluorescence intensity on these GUVs in the presence of GUVs with varying PI(3,5)P_2_ contents against that in the absence of PI(3,5)P_2_ GUVs. EGFP fluorescence intensity on GUVs were background corrected against the signal in the absences of EGFP-PI3Kα. Each data point represents average ± S.D.

To test the possibility that PI(3,5)P_2_ can physically displace the p110α-p85α heterodimer from PI(4,5)P_2_-containing membranes during the enzyme catalysis, we performed confocal microscopic imaging experiments under the same conditions. Specifically, we monitored the binding and unbinding of enhanced green fluorescence protein (EGFP)-tagged PI3Kα (i.e., EGFP-p110α + p85α) to and from giant unilamellar vesicles (GUVs) containing POPC/POPS/PI(4,5)P_2_/1,2-dioleoyl-sn-glycero-3-phosphoethanolamine-N-(lissamine rhodamine B sulfonyl) (Rh-PE) (67:20:3:10) in the presence of externally added PI(3,5)P_2_-containing GUVs. Rh-PE was used as a marker for PI(4,5)P_2_-containing GUVs. It was reported that a short phosphotyrosine peptide (pY2) induces conformational changes and membrane binding of Class I PI3K by simulating the effect of binding of p85 to an activated growth factor receptor or its adaptor proteins ^42^. Upon adding pY2 to the mixture containing EGFP-PI3Kα and POPC/POPS/PI(4,5)P_2_/Rh-PE GUVs, a significant increase in EGFP fluorescence intensity on these GUV surfaces was detected (**Fig. 4E**), indicating that pY2 triggered binding of EGFP-PI3Kα to PI(4,5)P_2_-containing GUVs. We then added POPC/POPS/PI(3,5)P_2_ (80-*x*/20/*x*; *x* = 0-3 mole%) GUVs to the mixture, which caused immediate and full unbinding of EGFP-PI3Kα from PI(4,5)P_2_-containing GUVs as essentially all EGFP fluorescence intensity was shifted from PI(4,5)P_2_-containing GUVs to PI(3,5)P_2_-containing GUVs **(Fig. 4E, 4F)**. The PI(3,5)P_2_ concentration dependency of EGFP-PI3Kα dissociation from PI(4,5)P_2_-containing GUVs shows that even 0.5 mole% of PI(3,5)P_2_ can remove >80% of EGFP-PI3Kα from GUVs containing 1 mole% PI(4,5)P_2_, again demonstrating the efficiency of this PI(3,5)P_2_-mediated *trans*-vesicular displacement (**Fig. 4F**). Also the PI(3,5)P_2_ concentration dependency of EGFP-PI3Kα binding to PI(3,5)P_2_ GUVs was essentially the mirror image of that of EGFP-PI3Kα dissociation from PI(4,5)P_2_ GUVs (**Fig. 4F**). Collectively, these enzyme kinetic and membrane binding data show that PI(3,5)P_2_ effectively inhibits the Class I PI3K activity in a *trans*-membrane mode by physically displacing it from the substrate (i.e., PI(4,5)P_2_)-containing membranes. The high efficiency of this inhibition also supports the notion that PI(3,5)P_2_ can inhibit cellular activity of Class I PI3K by the same mechanism.

### PI(3,5)P_2_ inhibits Class I PI3K by displacing it from the plasma membrane of mammalian cells via direct interaction

To check if this inhibition-by-displacement mechanism also works in living cells, we performed a wide range of quantitative cellular imaging measurements while modulating the cellular PI(3,5)P_2_ level. For PIKfyve inhibition, either siRNA knockdown or pharmacological inhibition was employed depending on feasibility under specific experimental conditions. First, we performed total internal reflection fluorescence microscopy (TIRFM)-based single molecule tracking of PI3Kα at the PM of p110-null mouse embryonic fibroblast (MEF) cells, whose endogenous p85α and p85β regulatory subunits were also depleted by siRNAs (**Supplementary Fig. S5F**). Specifically, we exogenously expressed EGFP-p110α-WT-p85α-WT (PI3Kα WT) and EGFP-p110α-WT-p85α-K653A (PI3Kα-K653A) at the lowest possible levels, respectively, in these MEFs for single molecule tracking analysis and compared their PM residence upon PDGF stimulation. In general, EGFP-PI3Kα WT rapidly appeared on and disappeared from the PM after PDGF stimulation (**Supplementary Fig. S10A**). Single molecule tracking analysis showed that the average PM dwell time of EGFP-PI3Kα WT, expressed in terms of the half-life of PM dissociation, peaked (≂150 msec) at 5 min post-stimulation and declined afterwards (**Fig. 5A**). This analysis indicates that the population of PI3Kα WT at PM reached a maximum at 5 min after stimulation, which is well synchronized with the kinetics of PIP_3_ formation at the PM (see **Fig. 3A**). When compared with PI3Kα WT, PI(3,5)P_2_ binding-defective mutant PI3Kα-K653A exhibited much more sustained PM residence after reaching its peak (**Fig. 5A** and **Supplementary Fig S10A**). Furthermore, treatment of cells with a PIKfyve inhibitor, YM201636, greatly slowed the dissociation phase and elongated the PM dwell time of PI3Kα WT (**Fig. 5A** and **Supplementary Fig S10A**). Overall, these results are consistent with the notion that PI3Kα, which is recruited to the PM upon PDGF stimulation, is rapidly displaced from the PM via p85-PI(3,5)P_2_ interaction after PDGF stimulation. To monitor the spatiotemporal dynamics of PI3Kα under physiological conditions, we also determined the subcellular localization of endogenous Class I PI3K in HEK293 cells after PDGF stimulation by immunostaining using a p85-specific antibody. It has been reported that the amplitude and duration of growth factor signaling is regulated by receptor trafficking ^44, 45, 46^. These reports suggested that our observed inhibition of the PI3Kα activity on PM by lysosomal PI(3,5)P_2_ could occur by at least two distinct mechanisms; i.e., via direct trans-organelle movement or via endocytosis of PI3Kα. If PI3Kα reaches lysosomes by direct trans-organelle hopping, one would expect that endocytosis inhibition would not interfere with the kinetics of PIP_3_ decline and the lysosomal localization of PI3Kα. As is the case with the exogenously expressed PI3K, the relative population of endogenous Class I PI3K at PM peaked at 5 min after PDGF stimulation, then reached the basal level after 20 min (**Fig. 5B** and **Supplementary Fig. S10B**), which is again consistent with the kinetics of PIP_3_ formation at the PM (see **Fig. 3A**). Also, PM dissociation of Class I PI3K was greatly inhibited by PI(3,5)P_2_ depletion by PIKFyve knockdown (**Fig. 5B** and **Supplementary Fig. S10B**). However, inhibition of endocytosis in the presence of the clathrin inhibitor Pitstop2 ^47^, which has been also reported to inhibit clathrin-independent endocytosis ^48, 49^, did not affect PM dissociation of Class I PI3K (**Fig. 5B** and **Supplementary Fig. S10B**). Furthermore, Pitstop2 treatment did not change the kinetics of PIP_3_ decline (see **Supplementary Fig. S8A**). These results indicate that PDGF-stimulated Class I PI3K is rapidly displaced from the PM via direct p85-PI(3,5)P_2_ interaction, but not via endocytosis of PM-bound PI3K under our experimental conditions. Consistently, Class I PI3K, when stained with the p85-specific antibody, showed statistically significant colocalization with LAMP1-containing lysosomes at 10 min post-stimulation (**Fig. 5C** and **Supplementary Fig. S10C**). Colocalization of PI3K with lysosomal LAMP1 was synchronized with the decline in PIP_3_ formation at the PM (see **Fig. 3A**) and was abrogated by siRNA knockdown of PIKfyve, but not by Pitstop2 treatment (**Fig. 5C** and **Supplementary Fig. S10C**). Similar results were observed when a p110α-specific antibody was used in lieu of the anti-p85 antibody (**Supplementary Fig. S11A**).

**Fig. 5.**
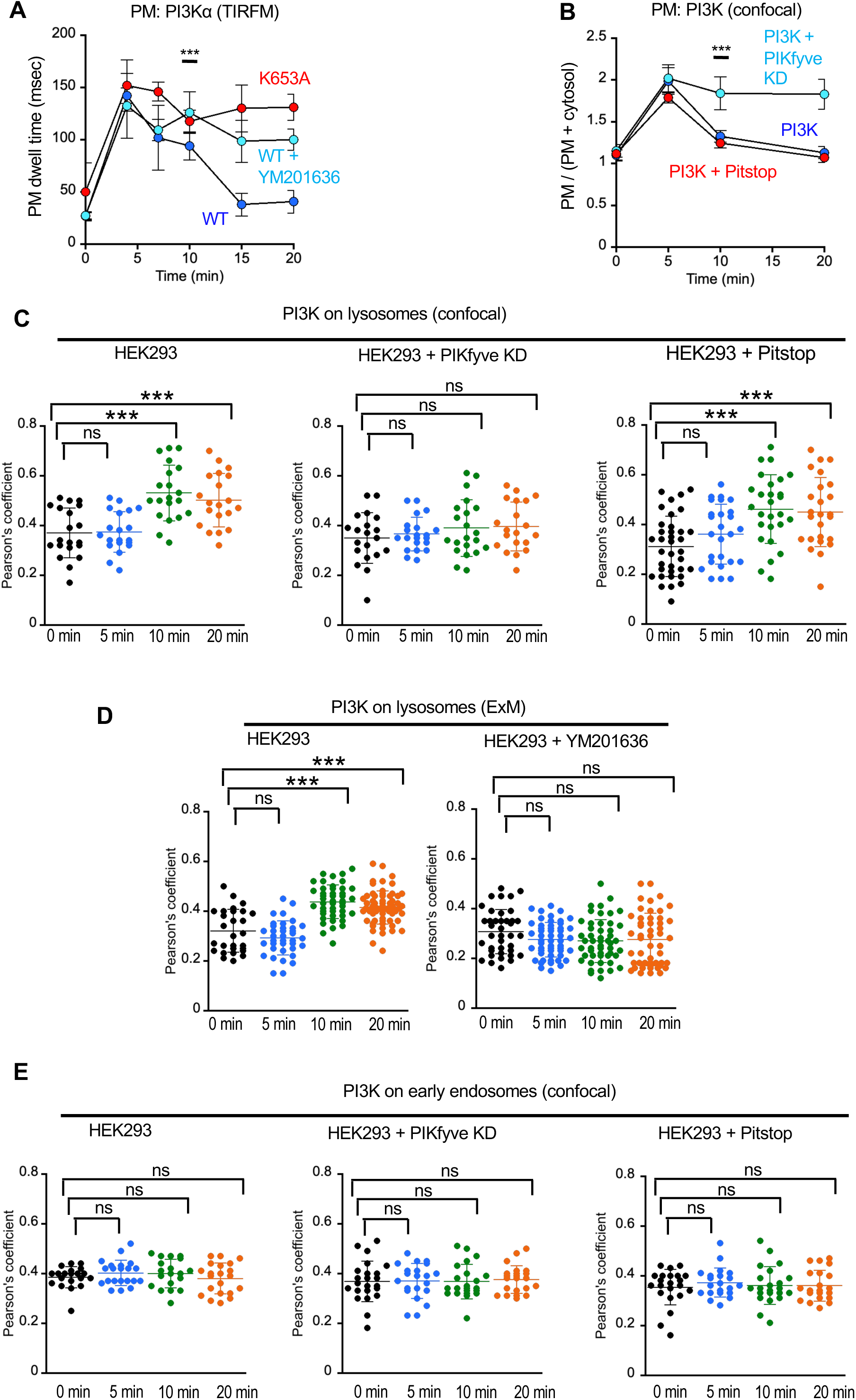
Subcellular localization of PI3Kα in response to PDGF stimulation. **A.** Kinetics of PDGF-stimulated (50 ng/ml) PM localization of PI3Kα WT (blue) and K653A (red) in p110α-null MEF cells. The endogenous p85α and p85β of MEF cells were suppressed by siRNAs and human EGFP-tagged PI3Kα WT (p110α-p85α-WT) (blue) and PI3Kα-K653A (p110α-p85α-K653A) (red) were exogenously expressed. PI3Kα WT was also inhibited by 0.8 μM YM201636 (1 h) (cyan). PM dwell time was calculated as *ln*2/*k* where *k* is the dissociation rate constant from PM. Data points indicate averages ± S.D.’s from five independent measurements (>30 cells per measurement). At 10 min, *p* <0.0001 for both between PI3Kα WT and PI3Kα-K653A and between PI3Kα WT and PI3Kα WT + YM201636. **B.** Degree of PM localization of endogenous PI3Kα before (blue) and after siRNA-mediated PIKfyve knockdown (KD) (cyan) and inhibition by 30 μM Pitstop2 for 15 min (red). Relative population of PI3Kα on PM was calculated as *I*_PM_ /(*I*_PM_ *+ I*_cytosol_) where indicated the fluorescence intensity at the PM and cytosol, respectively. Endogenous PI3Kα was stained with p110α-specific antibody. Data points indicate averages ± S.D.’s from three independent determinations (>10 cells per measurement). At 10 min, *p* <0.0001 for both between PI3K and PI3K + PIKfyve KD and between PI3K and PI3K + Pitstop2. **C.** Time-dependent lysosome localization of endogenous PI3K with and without PIKfyve knockdown (KD) or Pitstop2 treatment (30 μM, 15 min) was determined by monitoring the Pearson’s correlation coefficient between p85 and LAMP1 at different times. For HEK293 cells, *p* = 0.904 (5 min), <0.0001 (10 min), 0.00029 (20 min); For HEK293 + PIKfyve KD, *p* = 0.562 (5 min), 0.249 (10 min), 0.150 (20 min); For HEK293 + Pitstop2, *p* = 0.114 (5 min), <0.0001 (10 min), 0.00021 (20 min). **D.** The same as Fig. 4C except that expansion microscopy data (see **Fig. S10D**) were analyzed. For HEK293 cells, *p* = 0.148 (5 min), <0.0001 (10 min), <0.0001 (20 min); For HEK293 + YM201636, *p* = 0.053 (5 min), 0.05 (10 min), 0.138 (20 min). **E.** Time-dependent early endosome localization of endogenous PI3K (no treatment, with 30 μM Pitstop2, and with PIKfyve KD) was determined by monitoring the Pearson’s correlation coefficient between p85 and EEA1 at different times. For HEK293 cells, *p* = 0.255 (5 min), 0.397 (10 min), 0.743 (20 min); For HEK293 + PIKfyve KD, *p* = 0.368 (5 min), 0.746 (10 min), 0.758 (20 min); For HEK293 + Pitstop2, *p* = 0.957 (5 min), 0.965 (10 min), 0.716 (20 min). Endogenous PI3K was stained with p85 antibody. For **4D-4E**, all data points from three independent determinations are shown with averages ± S.D.’s.

To obtain higher resolution images of PI(3,5)P_2_-dependent lysosomal localization of PI3K during its catalysis, we also performed the expansion microscopy of PI3K in HEK293 cells, using the p85 antibody, after PDGF stimulation. Imaging data after 4-fold expansion of the specimen more clearly demonstrate the lysosomal localization of PI3K after PDGF stimulation, which was again abrogated by PIKfyve inhibition (**Fig. 5D** and **Supplementary Fig, S10D**). Also, the kinetics of lysosomal localization of PI3K was well synchronized with the kinetics of PIP_3_ decline in the PM (see **Fig. 3A**). Collectively, our results show that both p85 and p110 subunits of PI3K are displaced from the PM and directly recruited to lysosomes in the presence of PI(3,5)P_2_.

Our results showing that Pitstop2 exerted no effect on the PM localization of PI3K (see **Fig. 5B**), lysosomal localization of PI3K (**Supplementary Fig. S10C**), and the kinetics of PDGF-induced PIP_3_ formation at the PM (**Supplementary Fig. S8A**) argue against the possibility that PI3K on PM is inhibited by lysosomal PI(3,5)P_2_ via endocytosis of PI3K. To preclude the possibility that PI(3,5)P_2_ facilitates endocytosis and lysosomal translocation of PI3K, we also monitored endocytic movement of endogenous PI3K to early endosomes in HEK293 cells after PDGF stimulation by immunostaining. Specifically, we monitored the time-lapse movement of PI3K from PM to early endosomes while inhibiting endocytosis by Pitstop2 and suppressing PI(3,5)P_2_ production by siRNA-mediated PIKfyve knockdown. After PDGF stimulation, PI3K did not show any detectable early endosome localization within 20 min whether or not cells were pre-treated with Pitstop2 and PIKfyve siRNA, respectively (**Fig. 5E** and **Supplementary Fig. S10E**). Similar results were observed when the p110α-specific antibody was used in lieu of the anti-p85 antibody (**Supplementary Fig. S11B**).

Collectively, these cell imaging results obtained by different imaging methods under various conditions strongly support the notion that under our experimental conditions PI(3,5)P_2_ present in lysosomes can inhibit the action of Class I PI3K at the PM by specifically binding to p85 and thereby directly removing Class I PI3K from its primary site of action.

### A small molecule inhibitor of p85α-cSH2-PI(3,5)P_2_ interaction prolongs PI3K activation

If PI(3,5)P_2_ inhibits Class I PI3K by our proposed displacement mechanism, one would expect that a small molecule inhibitor of p85-cSH2-PI(3,5)P_2_ interaction will prevent PM detachment of Class I PI3K and thus prolong its activation. We recently established a new strategy to develop specific and potent inhibitors of the lipid-SH2 domain interaction, which led to the discovery of a first-in-class lipid-targeting kinase inhibitor for acute myeloid leukemia ^50^. Based on this strategy, we searched for small molecules that can specifically block p85α-cSH2-PI(3,5)P_2_ binding from a custom-built library of ∼1000 non-lipidic molecules ^50^ by a high-throughput fluorescence quenching-based lipid-protein binding assay ^51^. Through multiple rounds of screening and characterization, we identified a compound (VG220) that selectively inhibited p85α-cSH2-PI(3,5)P_2_ binding (**Supplementary Fig. S12A**). We recently reported that the efficacy of lipid-protein interaction inhibitors can be best expressed in terms of *IC*_50_ and the maximal inhibition (*I*_max_) due to their unique mode of inhibitory actions ^50^. Its *IC*_50_ (= 4.2 ± 1.3 μM) and *I*_max_ (= 80 ± 10%) values indicate that VG220 can effectively block p85α-cSH2-PI(3,5)P_2_ binding at micromolar concentrations (**Supplementary Fig. S12B**). To demonstrate its specificity, we performed three sets of measurements. First, we measured the inhibitory activity of VG220 for p85α-cSH2-K653A, a PI(3,5)P_2_ binding-compromised mutant of p85α-cSH2. We found that VG220 did not significantly inhibit (i.e., *IC*_50_> 100 μM and *I*_max_ < 20%) binding of this mutant to PI(3,5)P_2_ vesicles (**Supplementary Fig. S12B**). We also found that VG220 did not effectively inhibit PI(3,5)P_2_ binding of p85α-nSH2 (**Supplementary Fig. S12B;** *IC*_50_> 50 μM and *I*_max_ < 40%), although p85α-nSH2 could also bind PI(3,5)P_2_ tightly (see **Supplementary Table S1** and **Supplementary Fig S1B**). Lastly, we observed that VG220 did not interfere with membrane binding of other membrane binding proteins, including two representative PIP_3_-binding PH domains from Akt1 and PDK1 (**Supplementary Fig. S12C**). Collectively, these results show that VG220 can effectively and selectively inhibit p85α-cSH2-PI(3,5)P_2_ binding at micromolar concentrations.

We then tested if VG220 could prevent PM detachment of PI3K and thus prolong Class I PI3K activation in cellular assays, not only to validate our proposed mechanism but also to develop a novel Class I PI3K activator. We first tested if VG220 could prevent PM displacement of Class I PI3K by growth factor-induced PI(3,5)P_2_. The single molecule tracking analysis of EGFP-PI3Kα-WT (i.e., p110α-p85α-WT) and EGFP-PI3Kα-K653A (i.e., p110α-WT-p85α-K653A) at the PM of p85α/β-double knockdown-p110-null MEF cells (see **Fig. 5A** for the experimental approach) at 10 min post PDGF stimulation showed that 5 μM VG220 enhanced the PM dwell time of PI3Kα WT by 33% (**Fig. 6A**). The specific nature of this effect was supported by the finding that VG220 did not further increase the intrinsically higher PM dwell time of PI3Kα-K653A due to its compromised PI(3,5)P_2_ binding (**Fig. 6A**). We then measured the effect of VG220 on the kinetics of PDGF-induced PIP_3_ formation in the same cells: Preincubation of PI3Kα-WT-expressing cells with 5 μM VG220 greatly slowed the PIP_3_ decline phase (**Fig. 6B**), indicating that blocking the p85α-cSH2-PI(3,5)P_2_ interaction substantially prolonged Class I PI3Kα activation. Again, the specific nature of this effect was confirmed by the finding that VG220 did not significantly slow the decline of PIP_3_ in PI3Kα-K653A (i.e., p110α-WT-p85α-K653A)-expressing p85α/β-double knockdown-p110-null MEF cells, which already had slower PIP_3_ decline than PI3Kα-WT-expressing cells (**Fig. 6C**). Furthermore, VG220 significantly slowed the decline phase of PDGF-induced PIP_3_ kinetics in WT HEK293 cells (**Supplementary Fig. S13A**) but had little to no effect on that in PI3KC2β-null-HEK293 cells (**Supplementary Fig. S13B**) that could not produce the induced pool of PI(3,5)P_2_ in response to PDGF stimulation and thus had a slower decline phase (see **Fig. 3B**). Collectively, these results show that VG220 prolongs PDGF-induced PI3Kα activation by specifically blocking p85α-PI(3,5)P_2_ binding and thus preventing PM displacement of PI3Kα. They in turn validate our proposed mechanism in which growth factor-induced PI(3,5)P_2_ terminates Class I PI3K activation by displacing it from the PM via specific interaction with the p85 subunit.

**Fig. 6.**
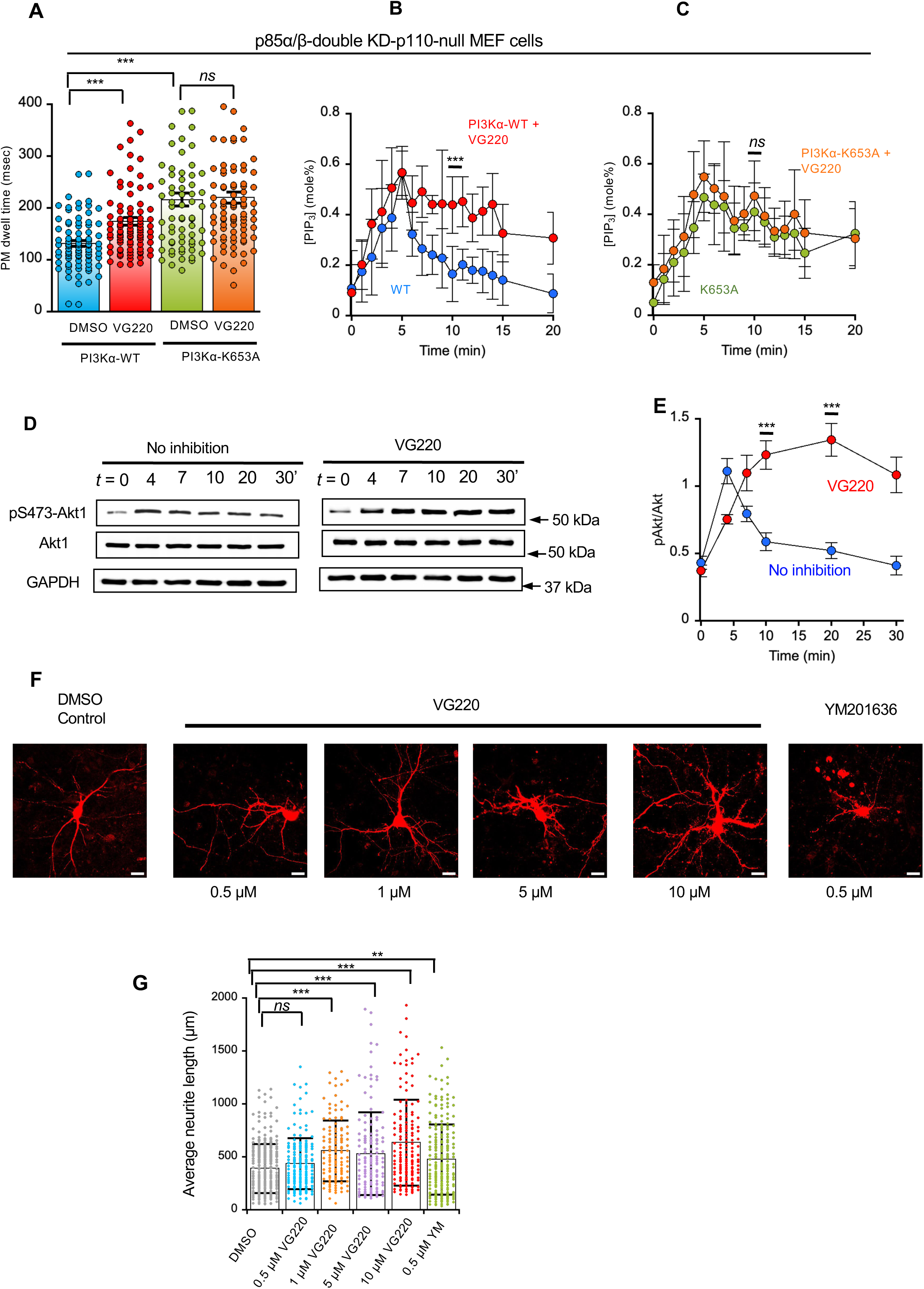
Effects of VG220 on the cellular activities of PI3K. **A.** Effects of VG220 (5 μM, 12h) on the PM dwell times of PI3Kα WT and K653A expressed in p110α-null MEF cells at 10 min after PDGF-stimulation (50 ng/ml). The PM dwell time was calculated as *ln*2/*k* where *k* is the dissociation rate constant from PM. *p* = <0.0001 (before and after VG220 treatment for PI3Kα-WT and between untreated PI3Kα WT and K653A) and 0.562 (before and after VG220 treatment for PI3Kα-K653A). **B.-C.** Effects of VG220 (5 μM, 12h) on PDGF (50 ng/ml)-stimulated formation of [PIP_3_] at the PM by PI3Kα WT (**B**) and K653A (**C**) expressed in p110α-null MEF cells. For **A-C**, the endogenous p85α and p85β of MEF cells were suppressed by siRNAs and human EGFP-tagged PI3Kα WT (p110α-p85α-WT) and PI3Kα-K653A (p110α-p85α-K653A) were exogenously expressed. *p* values between untreated and VG220-treated cells were <0.0001 (PI3Kα WT) and 0.9 (PI3Kα-K653A) at 10 min after PDGF stimulation. Data points indicate averages ± S.D.’s from five independent measurements (>30 cells per measurement). **D.** The effect of VG220 (5 μM, 12h) on the time course of PDGF-stimulated phosphorylation of S473 of endogenous Akt1 in HEK293 cells. Representative western blots (*n* = 3) show pS473-Akt1 and total Akt1 detected by immunoblot with the pS473-Akt1-specific and Akt1-specific antibodies, respectively. Glyceraldehyde 3-phosphate dehydrogenase (GAPDH) was used as a loading control. **E.** Quantification of Fig. 3D. *p* <0.0001 at 10 min after PDGF stimulation. **F.** The effects of VG220 (0-10 μM, 12h) and YM201636 (0.5 μM, 12h) on the neurite length of mouse cortical neurons. A representative image from >90 similar neurons is shown for each condition. Scale bars indicate 100 μm. **G.** Quantification of Fig. 3F. All data points from triplicate independent measurements (20-30 neurons per measurement) are shown with averages ± S.D.’s. *p* = <0.0001 (before and after 1-10 μM VG220 treatment), 0.552 (before and after 0.5 μM VG220 treatment) and 0.00147 (before and after 0.5 μM YM201636 treatment).

Our results also suggest that an inhibitor of p85-PI(3,5)P_2_ binding may serve as a Class I PI3K activator by prolonging its growth factor-mediated activation. Class I PI3K activators have been under development as a therapy to promote tissue regeneration and wound healing ^52^. To test if a p85-PI(3,5)P_2_ binding inhibitor such as VG220 can serve as a Class I PI3K activator under physiological conditions, we performed two functional assays. We first monitored the effect of VG220 on the phosphorylation of endogenous Akt1 on S473 (pS473) in HEK293 cells as a cellular readout of the PI3K signaling axis (see **Fig. 3C and D** for the approach). When compared to untreated HEK293 cells in which pS473 of Akt1 was reversed within 20 min after PDGF stimulation, pS473 remained largely unchanged after 20 min when cells were pre-treated with 5 μM VG220 (**Fig. 6D, 6E**), indicating that VG220 can indeed promote sustained activation of the PI3K-Akt signaling axis. We then measured the effect of VG220 on the neurite outgrowth, which has been used as a phenotypical readout of the Class I PI3K activity in neuronal cells ^52^. When compared to DMSO controls, VG220 induced substantial neurite growth of mouse cortical neurons in a dose-dependent manner (**Fig. 6F, 6G**). At 10 μM (i.e., 2.5 x *IC*_50_) VG220 promoted the neurite outgrowth by 63% over the control and was even more effective than 0.5 μM (i.e., >10 x *IC*_50_) of the PIKFyve inhibitor, YM201636. These results thus suggest that a p85-PI(3,5)P_2_ binding inhibitor may serve as a physiological activator of Class I PI3K that may promote tissue regeneration and wound healing.

### Hot-spot mutations in p85 cause sustained Class I PI3K activation by inhibiting p85-PI(3,5)P_2_ binding

Somatic mutations of p110α and p85α subunits have been found in various cancer cells and some ‘hot-spot’ mutations are known to drive cancer cell transformation ^5, 6, 53^. Although much is known about the effects of hot-spot mutations in p110α, less is known about those of hot-spot mutations in p85α. The most common hot-spot p85α mutations are located in the nSH2 (e.g., G376R and K379E) and iSH2 domains (e.g., R461A and N564D) but some mutations (e.g., R642L) are also found in the PI(3,5)P_2_-binding site of the cSH2 domain ^5, 6, 54^. These mutations are generally known to enhance PI3K activity by relieving autoinhibition of p110 by p85 ^5, 6, 54^. As described earlier, although p85-cSH2 has higher PI(3,5)P_2_ specificity than p85-nSH2, the latter can also tightly bind PI(3,5)P_2_ (see **Supplementary Table S1**). We thus hypothesized that hot-spot mutations in p85-nSH2 could also affect the p85-PI(3,5)P_2_ interaction directly or indirectly. To test this notion, we measured the effects of the common oncogenic p85α mutations on the kinetics of PIP_3_ formation by reintroducing p85α mutants to HEK293 cells, whose endogenous p85α and p85β were depleted by siRNA (**Fig. S5G**). When compared to reintroduction of p85α WT, reintroduction of all common oncogenic p85α mutants into p85-deficient cells (**Supplementary Fig. S5H**) consistently led to more sustained PIP_3_-production (**Fig. 7A**), although the magnitude of their effects varied to some degree. Between the two iSH2 domain mutants, N564D behaved like other nSH2 and cSH2 mutants whereas R461A had only a minor effect on the PIP_3_ kinetics (**Fig. 7A**). This is consistent with the fact that the iSH2 region is not directly involved in PI(3,5)P_2_ binding but can indirectly affect PI(3,5)P_2_ binding of nSH2 and cSH2 domains. We also measured the cell-free activity of the selected p85α mutants (i.e., p110α WT + p85α-mutant (G376R, K379E, and R642L)) using POPC/POPS/PI(4,5)P_2_ (77:20:3) LUVs in the presence of separate POPC/POPS/ PI(3,5)P_2_ (80-*x*/20/*x*; *x* = 0-3 mole%) LUVs. Akin to the specific PI(3,5)P_2_-binding site mutant (K653A), these oncogenic mutants were not inhibited by PI(3,5)P_2_-containing vesicles (**Fig. 4D** and **Supplementary Fig. S14, A-C**). Collectively, these results indicate that many hot-spot mutations in the nSH2 and cSH2 domains of the p85α subunit of Class I PI3K promote sustained PI3K signaling activity by suppressing p85α-PI(3,5)P_2_ binding and consequent feedback inhibition of Class I PI3K. They also underscore the pathophysiological significance of p85α-PI(3,5)P_2_ binding in controlling the signaling activity of Class I PI3K.

**Fig. 7.**
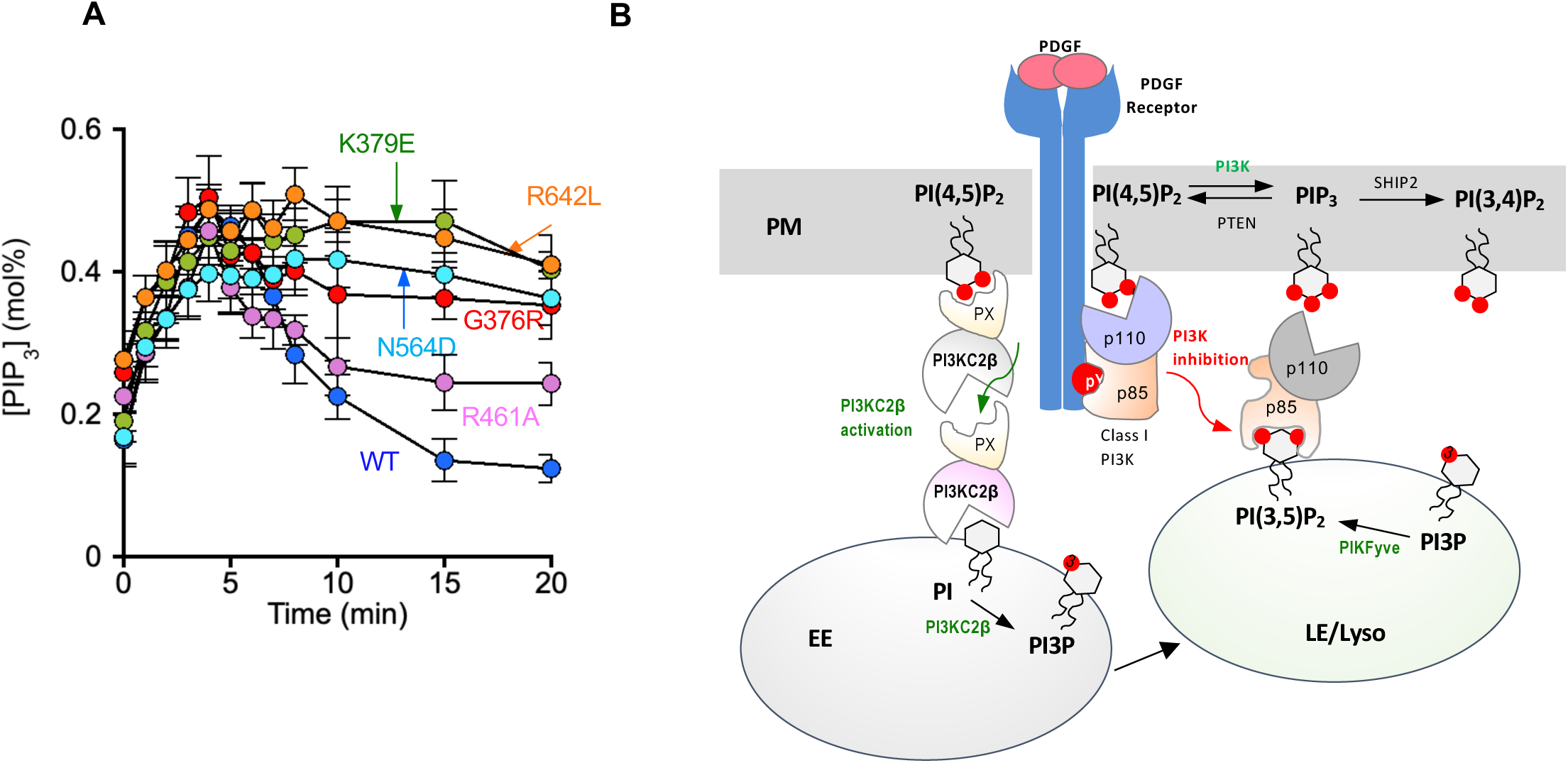
Effects of p85α mutations on kinetics of PIP_3_ formation at the plasma membrane and a hypothetical model of PI3K regulation by PI(3,5)P_2_. **A.** PDGF (50 ng/ml)-stimulated changes in [PIP_3_] at the PM of p85α/β-double knockdown HEK293 cells expressing a mouse p85α WT (blue) or a mutant, including K379E (green), R642L (orange), N564D (cyan), G376R (red), or R461A (light purple). [PIP_3_] values at different time points represent averages ± S.D.’s from three independent determinations with 5 cells per measurement. **B.** A hypothetical model of PI35P_2_-mediated feedback inhibition of Class I PI3K signaling. In response to growth factor stimulation (e.g., PDGF), Class I PI3K is recruited to the activated growth factor receptor (directly or via an adaptor protein) and activated. The activated PI3K phosphorylates PI(4,5)P_2_to PIP_3_ in the plasma membrane (PM), which is then dephosphorylated by SHIP2 to PI(3,4)P_2_. Growth factor stimulation also activates Class II PI3KC2β in the PM by an unknown mechanism. The activated PI3KC2β phosphorylates PI to PI(3)P primarily in early endosomes (EE). Constitutively active PIKfyve converts PI3KC2β-produced PI(3)P to PI(3,5)P_2_ mainly in late endosomes (LE) and lysosomes (Lyso), which inhibits Class I PI3K by specifically binding to p85 SH2 domains and thereby displacing the PI3K molecules from the PM. This feedback inhibition by PI(3,5)P_2_ terminates growth factor-induced PI3K activation.

## DISCUSSION

Due to their involvement in various human diseases, most notably cancer, Class I PI3Ks have been a focus of extensive studies ^5, 6^. Although much is known about the structure, function, and cellular activation mechanisms of Class I PI3Ks ^5, 6^, less is known about how the activated Class I PI3Ks are turned off after stimulation. Class I PI3Ks can be activated by diverse upstream proteins, including growth factor receptors (or receptor tyrosine kinases), G protein-coupled receptors, and RAS, often in combination ^5, 6^. Among Class I PI3K isoforms, PI3Kα is known to be activated primarily by growth factor stimulation^5, 6^ and we thus focused on growth factor-stimulation of PI3Kα in this study. Our spatiotemporally resolved quantification of PIP_3_ has shown that growth factor-induced formation of PIP_3_ at the PM of both non-cancerous and cancer-derived cell lines, including HEK293, NIH3T3, and HeLa cells, is rapidly (i.e., within 15 min) declined after reaching a peak at about 5 min post stimulation, resulting in only a short burst of PIP_3_ ^8^. It is generally thought that PI3K-mediated PIP_3_ formation is counterbalanced by lipid phosphatases, most notably PTEN and SHIP2/INPP4, which convert PIP_3_ to PI(4,5)P_2_ and PI(3,4)_2_/PI(3)P, respectively. However, neither genetic ablation of PTEN nor SHIP2 inhibition nor INPP4B knockdown ^8^ substantially alters the kinetics of PIP_3_ disappearance, indicating that the rapid removal of PIP_3_ at PM is not primarily driven by enzymatic transformation of PIP_3_ by lipid phosphatases. These results also suggest that lipid phosphatases alone cannot attenuate the PIP_3_ signal at PM as long as Class I PI3K remains active. Since the activation of Class I PI3K is not generally known to be driven by rapid post-translational modification ^5, 6^, it would not be readily reversed by counterbalancing modification under normal physiological conditions. One can speculate alternative mechanisms, such as degradation or endocytic sequestration of Class I PI3K. However, conclusive evidence for these mechanisms has not been reported. The present study shows that hitherto unknown PI(3,5)P_2_-mediated feedback inhibition plays a pivotal role in reversing activation of Class I PI3K, most notably PI3Kα, induced by PDGF and other growth factors. Key evidence for the new mechanism is provided by our spatiotemporally resolved *in situ* quantification of two other 3’-phosphoinositides, PI(3)P and PI(3,5)P_2_, which was made possible by our newly developed ratiometric sensors for these lipids. In particular, development of the PI(3,5)P_2_-specific ratiometric sensor represents a major technical breakthrough that has enabled us to gain new mechanistic insight into PI(3,5)P_2_-mediated cell regulation. Our imaging data under various conditions clearly show that growth factor stimulation leads to Class II PI3KC2β-mediated formation of PI(3)P on endosomes, which is converted into a special pool of PI(3,5)P_2_ by PIKfyve on late endosomes/lysosomes (**Fig. 7B**). This induced pool of PI(3,5)P_2_, but not the steady-state pool of PI(3,5)P_2_, turns off the PIP_3_-producing activity of Class I PI3K by displacing it from the PI(4,5)P_2_-abundant PM through direct interaction with the two SH2 domains in the p85 subunit (**Fig. 6C**). This inhibition-by-displacement mechanism is supported by multiple lines of biophysical, enzyme kinetics, cell imaging, and pharmacological data.

Results from the cell-free enzyme activity assay and imaging measurements show that PI(3,5)P_2_ can efficiently and potently inhibit PI3Kα acting on PI(4,5)P_2_-containing vesicles in a *trans*-vesicular mode. The enzyme activity of PI3Kα on PI(4,5)P_2_-containing vesicles is reduced by 70% in the presence of separate vesicles containing equimolar PI(3,5)P_2_. Confocal imaging data show that PI(3,5)P_2_ in separate vesicles can efficiently displace activated PI3Kα bound to PI(4,5)P_2_-containing vesicles under the same conditions. Our previous studies have shown that the concentration of PI(4,5)P_2_ at the PM fluctuates between 0.6 and 1.0 mol% during PDGF-induced PI3K activation in various mammalian cells ^8, 34^. The present study shows that although spatially averaged PI(3,5)P_2_ concentration is raised from 0.15 to 0.25 mol% after PDGF stimulation, the local PI(3,5)P_2_ concentration varies widely, with some endolysosomal spots exhibiting up to 0.8 mol% of PI(3,5)P_2_ (see **Fig. 2E**). Our immunostaining data (**Fig. S10B**) suggest that only a small fraction of endogenous Class I PI3K is recruited to the PM upon PDGF stimulation, which is consistent with previous reports ^55, 56^. Thus, quantitative inhibition of PM-bound Class I PI3K by endosomal/lysosomal PI(3,5)P_2_ should be locally feasible and efficient under physiological conditions.

This notion is fully supported by our quantitative cell imaging. Both single molecule tracking of exogenously expressed PI3Kα and immunostaining of endogenous PI3K show that Class I PI3K is recruited to the PM upon PDGF stimulation but rapidly displaced from the PM via PI(3,5)P_2_-p85 interaction. The kinetics of PI(3,5)P_2_-mediated PM displacement of PI3K is well synchronized with the kinetics of PIP_3_ decline at the PM, indicating that the PI(3,5)P_2_-mediated PM displacement of PI3K is primarily responsible for the rapidly decreasing PI3K activity at the PM. Importantly, both conventional and super-resolution expansion microscopy-based imaging demonstrate that the PI(3,5)P_2_-p85 interaction moves a majority of PM-associated PI3K to lysosomes, the kinetics of which is again well synchronized with the kinetics of PIP_3_ clearance at the PM. The lysosomal movement of PI3K may be also facilitated by the growth factor-induced dispersion of lysosomes to the cell periphery ^57^.

It is well documented that the amplitude and duration of growth factor signaling is regulated by receptor trafficking ^44, 45, 46^. It was recently reported that PI3Kα is endocytosed upon growth factor stimulation and actively produces PIP_3_ in endomembranes ^58^. We ^8^ and others ^59^ have reported endocytosis-mediated delivery of PI(3,4)P_2_ to endomembranes for downstream signaling pathways; however, our spatiotemporally resolved quantitative imaging detected neither the substrate (i.e., PI(4,5)P_2_) nor the product (i.e., PIP_3_) of Class I PI3K on endomembranes, early endosomes in particular, after PDGF stimulation ^8^. Further studies are needed to fully understand the origin of this discrepancy but our results clearly show that the unique, induced pool of PI(3,5)P_2_ on endomembranes moves PI3K from the PM to lysosomes through direct interaction with the SH2 domains of p85, but not through modulating endocytosis of PI3K. Also, the fact that PIKfyve inhibition, which removes all lysosomal PI(3,5)P_2_ and causes sustained PIP_3_ formation at the PM, has no effect on early endosomal localization of PI3K precludes the possibility PI(3,5)P_2_ exerts its inhibitory effect on Class I PI3K by modulating its endocytosis. In fact, although PI(3,5)P_2_ has been reported to be involved in endocytic recycling to the PM, its involvement in receptor endocytosis to early endosomes has not been reported to date ^14^.

In this study, we demonstrated the pathophysiological relevance and significance of the feedback inhibition of Class I PI3Ks by PI(3,5)P_2_-p85 interaction by both pharmacological and genetic approaches. We first show that a small molecule inhibitor of p85α-cSH2-PI(3,5)P_2_ interaction (VG220) specifically blocks PM detachment of PI3Kα by intracellular PI(3,5)P_2_ and thereby prolongs its PDGF-stimulated activation at the PM, leading to sustained Akt activation and enhanced neurite outgrowth. Although Class I PI3K-targeting drug development has mainly focused on inhibitors for cancers and other diseases, there has been a significant interest in development of Class I PI3K activators for promoting tissue regeneration and wound healing^52^. The fact that we were able to achieve efficient PI3Kα activation by inhibiting p85α-cSH2-PI(3,5)P_2_ interaction not only validates the physiological relevance of our proposed mechanism but also suggests that p85-PI(3,5)P_2_ binding inhibitors may serve as a therapeutic agent for promoting tissue regeneration and wound healing. Further optimization of VG220 to improve its potency and specificity and evaluation of these new compounds for various biological activities are currently in progress, which is beyond the scope of this report.

We also show that hot-spot somatic mutations in the p85 subunit lead to sustained Class I PI3K activation by suppressing PI(3,5)P_2_-p85 interaction. Extensive somatic mutations of Class I PI3Ks have been identified in various human cancer ^5, 6^. In particular, *PIK3CA*, which encodes the p110α catalytic subunit of PI3Kα, is the most frequently mutated gene in breast cancer ^5, 6^. Common hot-spot mutations in p110α (e.g., E542K, E545K and H1047R) activate PI3K by perturbing inhibitory intramolecular interaction between p85 and p110α and/or enhancing membrane affinity of p110α ^5, 6, 53^. Somatic mutations of *PIK3R1*, which encodes p85α, is less common than *PIK3CA* mutations, but they are found at high frequency in several cancer types, including endometrial, colorectal, and prostate cancer and glioblastoma ^54^. The most common p85α point mutations are located within the nSH2 domain and iSH2 domain ^54^. It has been speculated that hot-spot mutations in the nSH2 domain of p85α, such as G376R and K379E, may cause gain-of-function effects by weakening the inhibitory p110-p85α interaction ^54^. Our results show that these mutations as well as mutations in the cSH2 neither constitutively activate PI3Kα nor increase the amplitude of the PIP_3_ signal. Instead, they suppress p85-PI(3,5)P_2_ interaction and thereby block the feedback inhibition of Class I PI3K by PI(3,5)P_2_, causing sustained activation of Class I PI3K. These results not only provide further credence to the pathophysiological significance of newly discovered PI(3,5)P_2_-mediated feedback inhibition of Class I PI3K activity but also suggest a new therapeutic possibility of treating cancer patients with p85α mutations by modulating the cellular level of a specific pool of PI(3,5)P_2_.

Lastly, our results demonstrating the role of PI3KC2β-generated PI(3,5)P_2_ in feedback inhibition of Class I PI3K call for caution with the use of PIKfyve inhibitors for cancer therapy. It has been reported that PIKfyve inhibitors suppress various cancer cells by disrupting their endo-lysosomal pathway, lysosomal homeostasis, and autophagy ^17, 18^ Our study shows the presence of two separate (i.e., steady-state and growth factor-induced) pools of PI(3,5)P_2_ with different functions. Since PIKfyve is the only source of both pools of PI(3,5)P_2_, complete inhibition of PIKfyve would not only cause autophagy inhibition but also amplify the PI3K-Akt signaling axis in tumor cells, which would exert opposing effects on tumor cells. Thus, a combination of a PIKfyve inhibitor and a PI3K-Akt pathway inhibitor may be necessary to achieve full anti-tumor activity. This notion is supported by a recent report showing that Akt inhibition was required for the cytotoxicity of PIKfyve inhibitors against cancer cells ^60^.

## ACKNOWLEDGEMENTS

This work was supported by grants from the National Institutes of Health (R35GM122530 to WC, R56MH107387 to LWG and R35CA210057 to JJZ), the European Union (ERC AdG 884 281 to VH) and the Deutsche Forschungsgemeinschaft (TRR186/ A08 to VH). RG acknowledges funding support from Scialog grant #28707, sponsored jointly by Research Corporation for Science Advancement and the Walder Foundation. WC thanks Kyli Berkley, Elena Ristevska and Madalyn Puckett for their assistance in the SPR and the neurite outgrowth assays. WC also thanks Dr. Vladimir Gevorgyan for sharing his small molecule library.

## AUTHOR CONTRIBUTIONS

JS and JZ carried out lipid quantification and single molecule imaging. SS and IS performed SPR studies. JZ contributed to inhibitor development. AS and JS synthesized solvatochromic fluorophores and inhibitors. YH contributed to initial lipid sensor development. BA and LWG contributed to the neurite outgrowth assay. WTL, PAG, and VH assisted in PI3KC2β work. WW and RG contributed to expansion microscopy imaging. JJZ assisted in p110 gene ablation work. WC conceived and supervised the work and wrote the paper.

## DECLARATION OF INTERESTS

The authors declare no competing interests.

## METHODS

### Materials

1-Palmitoyl-2-oleoyl-sn-glycero-3-phosphocholine (POPC), 1-palmitoyl-2-oleoyl-sn-glycero-3-phosphoethanolamine (POPE), and 1-palmitoyl-2-oleoyl-sn-glycero-3-phosphoserine (POPS), cholesterol, and liver phosphatidylinositol (PI) were from Avanti Polar Lipids. 1,2-dipalmitoyl derivatives of phosphatidylinositol 3-phosphate (PI(3)P), phosphatidylinositol 4-phosphate (PI(4)P), phosphatidylinositol 5-phosphate (PI(5)P), phosphatidylinositol 4,5-bisphosphate (PI(4,5)P_2_), phosphatidylinositol 3,4-bisphosphate (PI(3,4)P_2_), phosphatidylinositol 3,5-bisphosphate (PI(3,5)P_2_), and phosphatidylinositol 3,4,5-trisphosphate (PIP_3_) were from Cayman Chemical Co. All cell lines were purchased from ATCC. PTEN-null MEF cells were a generous gift of Dr. Nissim Hay of University of Illinois at Chicago. Human platelet-derived growth factor (PDGF)-BB were from PEPROTECH. Human epidermal growth factor (EGF) was from Sigma-Aldrich. The pY2 peptide (ESDGGpYMDMSKDESIDpYVPMLDMKGDIKYA) was synthesized as described ^43^. The pY peptide (F-Ahx-ADNDpYIIPLPD) was custom-purchased from AlanScientific. Acrylodan and WCB1 were synthesized as described ^36^. YM201636, apilimod, SAR405, and AS1938909 were from AdooQ BioSciences, MedKoo Biosciences, Cayman Chemical, and Echelon Biosciences, respectively. Antibodies for pS473-Akt1, Akt1, and glyceraldehyde 3-phosphate dehydrogenase were from Cell Signaling Technologies. Antibodies for pS473-Akt1, Akt1, p85, p110α, Vps34, PI3KC2 α, INPP4B, Lamp1, EEA1, and GAPDH were from Cell Signaling Technologies. The antibodies for PI3KC2β and PIKfyve were from Abcam and Santa Cruz Biotechnologies, respectively. Goat anti-Rabbit IgG (H+L) secondary antibodies were from Thermo-Fisher.

### Cell lines

HEK293, NIH 3T3, MEF, and HeLa cells were purchased from ATCC and cultured in DMEM containing 4.5 g/l D-glucose, L-glutamine, 110 mg/l sodium pyruvate, 10% heat-inactivated FBS, 100 U/ml penicillin, and 100 μg/ml streptomycin (Gibco). Cells were routinely tested for mycoplasma contamination.

### Class I PI3K expression and purification

p85α-cSH2 was subcloned into the pESETB-eGFP vector for bacterial expression as an EGFP-tagged protein for SPR studies. Two subunits of PI3Kα, p110α and p85α, were co-expressed in insect cells as described previously for PI3Kβ ^43^. Recombinant baculoviruses for p110α and p85α were amplified in *Spodoptera frugiperda* (Sf9) cells. BTI-Tn-5B1-4 (High Five) suspension insect cells (2 × 10^6^ cells/ml) were then co-infected with p110α and p85α baculoviruses (MOI ratio = 1:3) for protein expression. Cells were harvested 72 h after infection. Cell pellets were suspended in a buffer containing 50 mM Tris, 300 mM NaCl, 10 mM imidazole, 1 mM tris(2-carboxyethyl) phosphine (TCEP), 1 mM phenylmethylsulphonyl fluoride (PMSF) (PH 7.9) and lysed using a hand-held homogenizer (Tissue Grinder Size C; Thomas Scientific). After centrifugation of the homogenate, the supernatant was incubated with the Ni-NTA resin for 2 h. The resin was then poured into a small column and washed with Buffer A (20 mM Tris-HCl, 300 mM NaCl, 10 mM imidazole (pH 7.9)) and then Buffer B (20 mM Tris-HCl, 300 mM NaCl, 40 mM imidazole (pH 7.9)). The protein was eluted with the elution buffer (20 mM Tris-HCl, 160 mM NaCl, 300 mM imidazole, 1 mM PMSF, 0.5 mM TCEP, 50 mM arginine, 50 mM glutamic acid (pH 7.9)).

### Preparation of the PI(3,5)P_2_ sensor

For sensor preparation, p85α-cSH2 was subcloned into the pET-30a(+) vector and sequentially mutated. The engineered PI(3,5)P_2_ sensor (WCB1-*e*p85α-cSH2) was expressed as a His_6_-tagged proteins in *E. coli* BL21 RIL codon plus cells. Protein expression was induced at 19°C with 0.5 mM (final concentration) isopropyl β-d-1-thiogalactopyranoside when the optical density of the media reached 0.6-0.8. Cells were harvested and cell pellets were suspended in lysis buffer containing 50 mM Tris, 300 mM NaCl, 10 mM imidazole, TCEP, 1 mM PMSF (PH 7.9) and lysed by sonication. The supernatant was incubated with the Ni-NTA resin after centrifugation of the homogenate. The resin mixture was then poured into a small column and washed Buffer A (20 mM Tris-HCl, 300 mM NaCl, 10 mM imidazole (pH 7.9)). After the resin became clear, the buffer solution was replaced by 5 ml of labeling buffer (50 mM Tris-HCl, 300 mM NaCl, 1 mM TCEP, 50 mM arginine, 50 mM glutamic acid (pH 8.0)). After adding 100 μl of WCB1 (10 mg/ml in dimethylsulfoxide (DMSO)), the mixture was gently shaken at room temperature for 2 h. The resin was first washed with Buffer A with 5% DMSO until the free dye was completely removed and then Buffer B (20 mM Tris-HCl, 300 mM NaCl, 40 mM imidazole (pH 7.9)). The sensor was eluted with the elution buffer (20 mM Tris-HCl, 160 mM NaCl, 300 mM imidazole, 0.5 mM TCEP, 50 mM arginine, 50 mM glutamic acid (pH 7.9)). The purity of the sensor was confirmed by the sodium dodecylsulfate polyacrylamide gel electrophoresis and the protein concentration was determined by the Bradford assay. The activity of the purified sensor was routinely checked by a quick three-point fluorometric measurement at 470 nm (*F*_470_) and at 530 nm (*F*_530_) using e.g., 10, 50, and 100 μM of POPC/POPS/PI(3,5)P_2_ (77:20:3) vesicles. The ratio (*F*_470_/*F*_530_) values from these measurements should lie within the 10% range of the standard calibration curves.

### Preparation of the PI(3)P and PI(4,5)P_2_ sensors

EEA1-FYVE-V21C was subcloned into the pGEX4T-1 vector and expressed as a GST-tagged protein in BL21 (DE3) pLysS cells. The expressed protein was extracted from the cells as described above. The cell supernatant was mixed with the glutathione resin and incubated with 100 μl of acrylodan solution (1 mg/ml in DMSO) in 5 ml of 20 mM Tris buffer, pH 7.4, containing 160 mM NaCl and 0.5 mM TCEP for 2h at room temperature with gentle mixing. Resins were washed with 100 ml of 20 mM Tris buffer with 160 mM NaCl at pH 7.4 with 5% DMSO. After washing, DAN-*e*EEA1 was incubated with 1 μl of thrombin (Invitrogen) in 1 ml of thrombin cleavage buffer (20 mM Tris, 160 mM NaCl, 25 mM CaCl_2_, 0.5 mM TCEP, 50 mM arginine, 50 mM glutamic acid at pH 8) overnight at 4°C. The eluted protein was used immediately used for imaging. The PI(4,5)P_2_ sensor, DAN-*e*ENTH, and the PIP_3_ sensor, DAN-eMyoX-tPH, were prepared as described previously ^33, 43^.

### Surface plasmon resonance (SPR) analysis

All SPR measurements were performed at 23°C in 20 mM Tris, pH 7.4, containing 0.16 M NaCl using a lipid-coated L1 chip in the BIACORE X100 system (Cytiva) as described previously ^25, 32^. LUVs of POPC/POPS/PI(3,5)P_2_ (77:20:3) and POPC/POPS (80:20) were used as the active surface and the control surface, respectively. Equilibrium SPR measurements were carried out at a flow rate of 5 μl/min to allow sufficient time for the R values of the association phase to reach near-equilibrium values (*R*_eq_). Sensorgrams were analyzed assuming a Langmuir-type binding between the protein (P) and protein binding sites (M) on vesicles (that is, P + M↔PM). The *R*_eq_ values were plotted against the protein concentrations (*P_o_*), and the *K_d_* was determined by nonlinear least squares analysis of the binding isotherm using the equation, *R*_eq_ = *R*_max_/(1 + *K_d_*/*P_o_*). For kinetic SPR measurements, the flow rate was maintained at 30 μl/min for both association and dissociation phases.

### Molecular docking analysis

The binding modes of the PI(3,5)P_2_ and p85-cSH2 were calculated using Autodock4 software (The Scripps Research Institute, La Jolla, USA) ^61^. 1,2-dioleoyl derivative of PI(3,5)P_2_ was drawn and energy-minimized with Chem Draw ultra and Chem 3D ultra, respectively. The structure was saved as sdf file and protein data bank (PDB) file was generated using OpenBabel2.3.1 software ^62^. The crystal structure of p85-cSH2 (PDB ID: IH9O) was used for docking analysis. The macromolecule was then prepared by removing the water molecules and adding hydrogens and Kollman charges and saved in PDBQT format for the Autodock4 program. A cube shape grid coordinates (dimension: *x* = 70, *y* = 70, *z* = 70 and center *x* = −0.861, *y* = 12.582, *z* = 5.833) were set to cover the binding site of p85-cSH2. To run the docking, the parameters were kept as default. Finally, the generated dlg file was analyzed and the lowest energy binding conformation was considered as the best docking pose. The docking pose was exported and illustrated using PyMOL (The PyMOL Molecular Graphics System, Version 2.0 Schrödinger, LLC).

### *In vitro* PI3K activity assay

All cuvette-based continuous PI3K activity assays were performed with the FluoroLog3 spectrofluorometer at 37°C in a 1 ml quartz cuvette (Hellma Analytics) as described previously ^43^. For the PI3K activity assay, 874 μl of 20 mM Tris buffer, pH 7.4, containing 0.16 M NaCl was mixed with 100 μl of POPC/POPS/PI(4,5)P_2_ (77:20:3 in mole%; 400 μg/ml) LUVs at the indicated concentration, 5 μl of 50 μM DAN-eENTH ^33^ (final concentration: 500 nM), 10 μl of 0.1 mM pY2 peptide (final concentration: 500 nM), and 10 μl of 0.1 M ATP (final concentration: 0.1 mM). The reaction was initiated by adding 10 nM of enzyme solution to the mixture continuously monitored by measuring the blue-shifted emission of the sensor at 470 nm with the excitation wavelength set at 380 nm.

### Cell culture and stimulation

HEK293 (or other) cells were seeded into 50 mm round glass–bottom plates and grown at 37°C in a humidified atmosphere of 95% air and 5% CO_2_ in Dulbecco’s modified Eagle’s medium (DMEM) with 4.5 g/l D-glucose, L-glutamine, 110 mg/l sodium pyruvate containing 10% heat-inactivated FBS, 100 U/ml penicillin, and 100 μg/ml streptomycin (Gibco) and cultured in the Corning cover glasses for 24 h before lipid quantification. All cell lines were cultured bi-weekly and stocks of cell lines were passaged no more than ten times for use in experiments. Growth factor stimulation was performed as described ^8^. Briefly, HEK293 (or other cells) were stimulated with 50 ng/ml PDGF-BB, EGF or IGF. Serum starvation was not employed because some inhibitors, such as YM201636, caused cell death under serum depletion.

### Gene knockout and knockdown

PI3KC2β-null HEK293 cells were prepared by CRISPR-CAS9 as described previously ^63^. siRNA-based gene knockdown was performed as reported previously ^64^. HEK 293 cells were grown to 30-40% confluency and transfected with various siRNA’s using the JetPRIME system (Polyplus-transfection) according to the manufacturer’s protocol. After cells were incubated with siRNA’s for 48 h and the suppressed expression level of a target protein was confirmed, they were used for further analysis.

### Western blot Analysis

Cells were seeded at a density of 3.0 × 10^5^ cells/well 24 h prior to various treatments. After treatments, cells were lysed in the NP40 lysis buffer (50 mM Tris, pH 7.5, 150 mM NaCl, 1 mM EDTA, 1% NP40, 10% glycerol, 10 mM NaF, 10 mM Na3VO4, and protease inhibitor cocktail, pH = 7.4) at 4°C and the mixture cleared by centrifugation. Samples were analyzed by SDS-PAGE. Proteins were separated and transferred to Immobilon-P PVDF membrane (0.45 μm, Millipore Sigma). The membrane was blocked with 5% bovine serum albumin for 1 h and incubated overnight at 4°C with various antibodies (1:500 dilution for the PIKfyve antibody and 1:1000 dilution for all other antibodies). After the unbound antibodies were removed by washing with 0.1% Tris buffer saline with 0.1% Tween20, the membranes were incubated with the horseradish peroxidase secondary antibody (1:2000 dilution) for 1 h at room temperature. The membranes were washed three more times with 0.1% Tris buffer saline with 0.1% Tween20 to remove the unbound horseradish peroxidase secondary antibody before imaging. The chemiluminescence intensity of protein bands in the gel was analyzed and documented by the Azure 500Q Imaging System.

### Single-molecule imaging analysis

Single-molecule imaging was performed using a custom-built total internal reflection fluorescence (TIRF) microscope. Mouse embryonic fibroblast (MEF) cells were treated with p85α and p85β siRNA’s and then plated on the 8-well chambered cover glass at the density of 1 × 10^5^ for 24 h, and EGFP-tagged-p110α and p85α (WT or K653A, respectively) were co-transfected into cells using the jetPRIME system (Polyplus-transfection) according to the manufacturer’s protocols. 800 nM YM201636 (24 h) was used to inhibit PIKfyve. Before and after stimulation with 50 ng/ml PDGF-BB, the EGFP signal on the PM was tracked for a given time and the data were analyzed as described previously ^65^. All single molecule tracking, data analysis and image processing were carried out with in-house programs written in MATLAB. The PM dissociation rate constant (*k*) was determined by the non-linear least-squares using a single exponential decay equation (i.e., *I* = *I*_0_ x *e*^−kt^ where *I* and *I*_0_ indicate the membrane fluorescence intensity and the maximal *I*, respectively). The PM dwell time was then expressed as the half-life of dissociation (= *ln*2/*k)*. >250 images from 5 independent measurements were analyzed for every data point.

### Colocalization analysis of endogenous PI3K with organelle makers

The same number (1 × 10^4^) of HEK293 cells were seeded onto 24-well plate-compatible coverslips and grown at 37 ℃ in a humidified atmosphere of 5% CO_2_ for about 24 h. Cells were washed twice with phosphate buffer saline (PBS) and then fixed in fresh 4% paraformaldehyde for 10 min, followed by 5 min incubation in 100 mM glycine in PBS to quench the fixation, and further washed twice with PBS, each time for 5 min The cells were then permeabilized with 0.1% (w/v) Triton X-100 in PBS at room temperature for 15 min, followed by treatment with the blocking buffer (5% (v/v) BSA in PBS) at room temperature for 30 min. Cells were incubated with the primary antibodies (1:200 diluted) against PI3K (p110α or p85) and an organelle marker (LAMP1 or EEA1) in PBS (with 1% BSA) overnight at 4 °C. Cells were then washed 4 times with PBS, each time for 5 min, and incubated with the secondary antibodies (Goat anti-Mouse Alexa Fluor™ 488 and Goat anti-Rabbit Alexa Fluor™ 568) (1:200 diluted in PBS with 1% BSA) for 2 h at room temperature. Cells were washed 4 times with PBS, each time for 5 min, and used for imaging. Imaging was performed with the custom-designed six channel Olympus FV3000 confocal microscope. Imaging data were analysed using ImageJ (FIJI, ImageJ 1.53t, with Coloc2 (3.0.5) plugin for colocalization analysis). The degree of colocalization was expressed in terms of Pearson’s correlation coefficient (*R*) ^66^, which was calculated from > 20 cells. The degree of PM localization of PI3K was calculated as *I*_PM_ /(*I*_PM_ *+ I*_cytosol_) where *I*_PM_ and *I*_cytosol_ indicated the fluorescence intensity at the PM and cytosol, respectively.

### Expansion microscopy imaging

The same number (1 × 10^4^) of HEK 293 cells were seeded onto 24-well plate-compatible coverslips and grown at 37°C in a humidified atmosphere of 5% CO_2_ for about 24 h. Cells were washed twice with PBS and then fixed in fresh 4% paraformaldehyde for 10 min. The cells were then washed with 100 mM glycine in PBS for 5 min to quench the fixation, and twice with PBS, each time for 5 min. Expanded cells were prepared using a protein-retention expansion microscopy (ExM) protocol with minor changes ^67, 68^. Briefly, the fixed cells were permeabilized with 0.1% (w/v) Triton X-100 in PBS at room temperature for 15 min and washed with the blocking buffer [5% (v/v) normal goat serum (NGS, Jackson ImmunoResearch) and 0.1% (w/v) Triton X-100 in PBS] at room temperature for 15 min. The cells were then incubated with the primary antibody solution (rabbit anti-p85 antibody with 1:50 dilution and mouse anti-LAMP1 antibody with 1:100 dilution in the blocking buffer) overnight at 4 °C. The next day, the cells were washed 4 times with the blocking buffer for 5 min each time, and incubated with the secondary antibody solution (goat anti-mouse Alexa Fluor 488 and goat anti-rabbit Alexa Fluor 568 with1:100 dilution in the blocking buffer) for 2 h at room temperature. Then cells were washed 4 times with PBS, each time for 5 min, and incubated with Acryloyl-X SE solution (0.1 mg/ml in PBS) for 3 h at room temperature. Afterwards, the cells were washed twice with PBS, each time for 15 min, and incubated in a gelling solution prepared by mixing monomer solution [1x PBS, 2 M NaCl, 8.6% (w/v) sodium acrylate (Pfaltz&Bauer), 2.5% (w/v) acrylamide, 0.15% (w/v) N,N’-Methylenebisacrylamide], ammonium persulfate (APS) [10% (w/v)], tetramethylethylenediamine (TEMED) [10% (w/v)], and 4-hydroxy-2,2,6,6-tetramethylpiperidin-1-oxyl (4HT) [0.5% (w/v)] at a 47:1:1:1 ratio. The cells were polymerized in an incubator at 37°C for 1 h, and then the gelled cells were digested overnight at room temperature with digestion buffer [1 mM EDTA, 0.5% Triton X-100, 0.05 M Tris-Cl, 1 M NaCl and 8 units/ml Proteinase K (New England Biolabs) in 1x PBS]. The gelled samples were washed and expanded 3 times with pure water, each time for 20 min. Finally, the expanded samples were trimmed and immobilized on Corning cover glasses (No.1.5) using the poly-L-lysine mounting method for imaging ^67^. Imaging was performed with the custom-designed six channel Olympus FV3000 confocal microscope using DAPI, Alexa Fluor 488 and Alexa Fluor 568 channels. Image data analysis was performed as described above for confocal images.

### Calibration of PI(3)P, PI(3,5)P_2_, and PIP_3_ sensors

*In vitro* calibration of WCB1-*e*p85α-cSH2 was performed with late endosome-mimicking GUVs (POPC/POPE/POPS/cholesterol/PI/PI(3,5)P_2_ (20-x/40/20/10/10/*x*: *x* = 0-2 mol%)) or POPC/POPS/PI(3,5)P_2_ (77/20/3) as described previously ^35^. GUVs were mixed with the lipid sensors (200 nM) and fluorescence imaging was performed with the custom-designed six channel Olympus FV3000 confocal microscope with the environmentally controlled full enclosure incubator (CellVivo). WCB1-*e*p85α-cSH2 was excited with the 405 nm laser source and the emission intensity was collected in two separate channels with the spectral detector setting of 435-465 nm (blue channel) and 495-525 nm (green channel). For each lipid concentration, at least 30 GUVs were selected for image analysis by Image-Pro Plus7. Calibration curve fitting for WCB1-*e*p85α-cSH2 was performed by non-linear least-squares analysis using the equation: *F*_B_/*F*_G_ = (*F*_B_/*F*_G_)_min_ + ((*F*_B_/*F*_G_)_max_ − (*F*_B_*F*_G_)_min_) / (1 + *K*_d_ /[PI(3,5)P_2_]). *F*_B_/*F*_G_, *K*_d_, (*F*_B_/*F*_G_)_max_, and (*F*_B_/*F*_G_)_min_ are the ratio of the fluorescence intensity in the blue channel to that in the green channel, equilibrium dissociation constant (in mol%), and the maximal and minimal *F*_B_/*F*_G_ values, respectively ^35^. Calibration of DAN-*e*EEA1 was performed using early-endosome-mimicking GUVs. Calibration of DAN-*e*MyoX-tPH was performed as described ^8^.

### *In situ* quantitative imaging

The same number (2.5 × 10^4^) of cells were seeded into Corning cover glasses and grown at 37 ℃ in a humidified atmosphere of 5% CO_2_ in phenol red-free DMEM supplemented with 10% (v/v) FBS, 100 U/ml penicillin G, and 100 mg/ml streptomycin sulfate and cultured in the plates for about 24 h before lipid quantification. Imaging was performed with the custom-designed six channel Olympus FV3000 confocal microscope with the environmentally controlled full enclosure incubator (CellVivo). Cells were maintained at 37℃ and with 5% CO_2_ atmosphere throughout the imaging period to maintain the cell viability. Typically, 20–30 femtoliter of the sensor solution was microinjected into the cell to reach the final cellular concentration of 200-400 nM. All image acquisition and imaging data analysis were performed as described for GUV calibration. All lipid determination was performed using the GUV calibration curves as described previously ^35^. The three-dimensional display of local lipid concentration profile was calculated using the Surf function in MATLAB.

### Small molecule inhibitor screening

The membrane binding of the EGFP-tagged p85α-cSH2 domain was measured by the fluorescence quenching assay at 25°C in untreated black polystyrene 96-well plates (Corning) using the Synergy™ Neo microplate reader (Agilent) as described previously ^50^. Briefly, our home-built library of 1000 non-lipidic molecules were screened for their activity to inhibit quenching of the EGFP fluorescence (emission at 516 nm and excitation at 485 nm) by a dark quencher lipid, dabsyl-PE, incorporated into lipid vesicles (e.g., POPC/POPS/PI(3,5)P_2_/dabsyl-PE (67/20/3/10 in mol%)). Once the optimal total lipid concentration for a fixed concentration of the protein (100 nM) in 20 mM Tris buffer (pH 7.4) containing 0.16 M NaCl was determined to ensure maximal membrane protein interaction (i.e., maximal EGFP quenching), 5 μM of each compound was added to the mixture to screen for dequenching activity. Those molecules that caused >50% of EGFP florescence dequenching were selected in two stages. To select PI(3,5)P_2_-specific inhibitors, those molecules that non-specifically inhibited binding of EGFP-p85α-cSH2 to POPC/POPS/dabsyl-PE (70/20/10)) were eliminated. Once lead compounds had been identified, their I_max_ and IC_50_ values of inhibitors were determined using the equation, I = I_max_ / (1 + IC_50_ / [I]) (I: % inhibition, [I]: inhibitor concentration) as described previously^50^ and the inhibitor with the lowest IC_50_ and the highest I_max_ values was selected for functional studies. Average and SD values were obtained from triplicate determinations. VG220 was synthesized and characterized as reported previously ^69^.

### Neurite outgrowth assay

Neuronal culture from cortex of newborn pups with either sex was prepared as described previously ^70, 71^, and neurons in culture were maintained in a 12-well plate at 37°C in 5% CO2. These neurons at days in vitro (DIV) 5 were transfected with the mScarlet ^72^ plasmid (Addgene Catalog# 85042) using Ca^2+^ phosphate transfection, and isolated individual neurons were transfected due to low transfection efficiency of this method. At DIV 6, neurons in each well was prepared to exhibit one of the following conditions: 0.5% DMSO as a negative control, 1 μM YM201636 as a positive control, and 0.5-10 μM VG220. After treatment, the plates were placed back in the 37°C incubator with 5% CO_2_. After incubation for 72 h at 37 °C and 5% CO2, cells were fixed by adding 4% paraformaldehyde for 10 min. Cells were washed 3 times in PBS before attaching coverslips to a glass slide with mounting media (Fisher Scientific). Coverslips and glass slides were left to harden overnight at room temperature. Image acquisition was performed with the Olympus FV3000 confocal microscope using a 60x oil immersion objective. Images were captured using excitation/emission wavelengths of 561/591 nm. A minimum of ten fields per coverslip were taken, and images were analyzed using Simple Neurite Tracer (SNT) ^73^ in Fiji/ImageJ software. The neuronal length was manually measured with the help of SNT, and all neurites measured started from a cell body. Treatments were performed in triplicate and the data is represented as an average of 3 biological repeats.

### Quantification and statistical analysis

Statistical analysis was performed using Synergy Kaleidagraph v.5.0. All data are presented as mean ± SD and were obtained from ≥ 3 independent experiments. Statistical significance was calculated by the unpaired Student’s t-test (ns, *p* > 0.05; *, p < 0.05; **, *p* < 0.01; ***, *p* < 0.001). The number of independent experiments, the number of total cells analyzed, and *p* values are reported in the figure legends. Sample sizes for cellular imaging and assays were chosen as the minimum number of independent observations required for statistically significant results.

## Notes

### Competing Interest Statement

The authors have declared no competing interest.

### Summary of Updates

A small molecule inhibitor of p85α-PI(3,5)P2 binding that specifically blocks the feedback inhibition of Class I PI3K by PI(3,5)P2 and thus serves as a PI3K activator has been included.

