## Supplementary figures and table for "PI(3,5)P_2_ Controls the Signaling Activity of Class I PI3K"

**CONTENTS**

**Supplementary Figures**

Fig.S1. Membrane and peptide binding properties of the p85α/β-SH2 domains and mutants determined by SPR (a-e) and fluorescence anisotropy (f) analyses
Fig.S2. Development and characterization of a ratiometric $PI(3,5)P_2$ sensor (WCB-*e*p85α-cSH2)
Fig.S3. Fig. S3. Effects of lipid kinases and phosphatases on subcellular localization of $PI(3,5)P_2$ detected by WCB-ep85α-cSH2 in HeLa cells.
Fig.S4. Effects of $PI(4,5)P_2$ changes in the PM on subcellular localization of WCB1-ep85α-cSH2 in HEK293 cells.
Fig.S5. Western blot analysis of HEK 293 cells after genetic modulation of lipid kinases and phosphatases.
Fig. S6. Time courses of the PDGF-stimulation changes in [$PI(3,5)P_2$] under different conditions.
Fig. S7. Protein engineering of a ratiometric PI(3)P sensor, DAN-*e*EEA1, and *in situ* quantification of cellular PI(3)P by DAN-*e*EEA1.
Fig. S8. Effects of lipid kinase and phosphatase inhibition on the kinetics of PDGF-induced $PIP_3$ changes
Fig. S9. Effects of PtdInsP on the enzyme activity of PI3Kα
Fig. S10. Time courses of subcellular localization of PI3Kα after PDGF stimulation of HEK293 cells
Fig. S11. Time courses of subcellular localization of p110α after PDGF stimulation of HEK293 cells
Fig. S12. Development and characterization of a small molecule inhibitor of p85-cSH2-$PI(3,5)P_2$ interaction.
Fig. S13. Effects of VG220 on the $PIP_3$ formation in WT and PI3KC2β-null HEK cells
Fig. S14. Inhibition of the enzymatic activity of PI3Kα-p85α mutants by $PI(3,5)P_2$

**Supplementary Tables**

Supplementary Table 1. Lipid and peptide binding properties of p85-SH2 domains and mutants

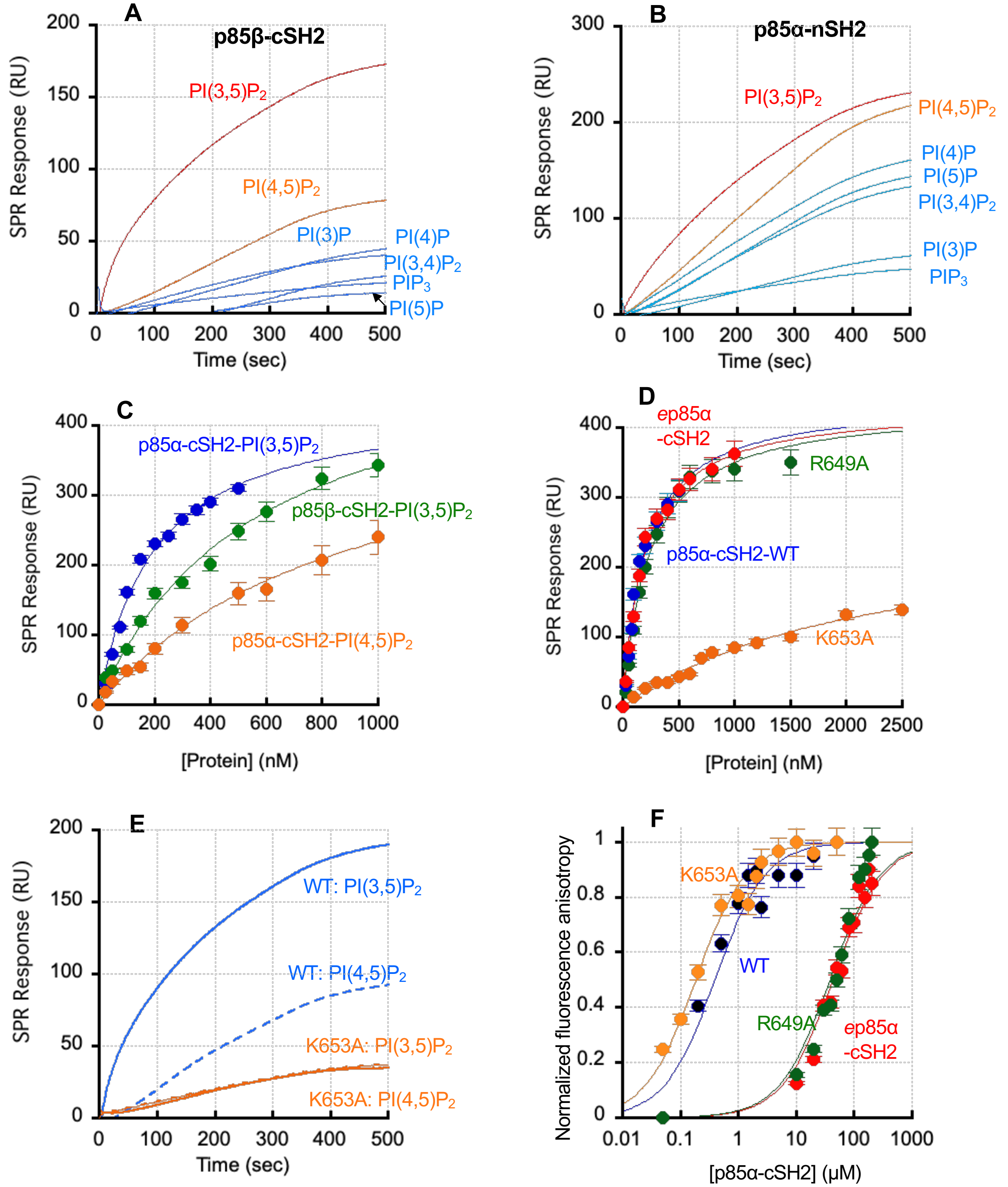


**Fig. S1. Membrane and peptide binding properties of the p85α/β-SH2 domains and mutants determined by SPR (A-E) and fluorescence anisotropy (F) analyses.**
**A.-B**. PtdInsP selectivity of p85β-cSH2 (A) and p85α-nSH2 (B) determined by SPR analysis. See Fig. 1a for experimental conditions. **C-D**. Determination of $K_d$ for binding of p85α/β-cSH2 WT to POPC/POPS/PI(3,5)$P_2$ (or PI(4,5)$P_2$) (77:20:3) LUVs (C) and p85αβ-cSH2 WT, mutants, and *e*p85α-cSH2 to POPC/POPS/PI(3,5)$P_2$ (77:20:3) LUVs (D). $K_d$ was determined by nonlinear least squares analysis of the binding isotherm using the equation, $R_{eq} = R_{max}/(1 + K_d/P_o)$ where $P_o$ , $R_{eq}$, and $R_{max}$, indicate the protein concentration, the maximal SPR response at each $P_o$ , and the saturating $R_{eq}$, respectively. **E.** Selectivity of p85α-cSH2 WT (cyan) and K653A (orange) for POPC/POPS/PI(3,5)$P_2$ (77:20:3) (solid line) over POPC/POPS/PI(4,5)$P_2$ (77:20:3) (dotted line) vesicles determined by SPR analysis. The protein concentration was 300 nM. Notice that K653A shows essentially no PI(3,5)$P_2$ selectivity. **F.** Binding of p85α-cSH2 WT, mutants, and *e*p85α-cSH2 to the peptide (F-Ahx-ADNDpYIIPLPD). The experimentally observed anisotropy (*A*) values were normalized ($A_{norm}$) using the equation: $A_{norm} = (A - A_{min}) / (A_{max} - A_{min})$ where $A_{max}$ and $A_{min}$ are maximal and minimal *A* values, respectively, for each measurement. The plots of $A_{norm}$ versus [p85α-cSH2] were analyzed by the non-linear least-squares analysis using the equation: $A_{norm} = 1/(1 + K_d/[\text{p85α-cSH2}])$.

**A**

**(1) PI(3,5)$P_2$ sensor protein engineering: ep85α-cSH2**

**(a) Introduction of a single Cys for WCB1 conjugation**: To eliminate C656 that cannot be used as a dye-conjugation site because of its involvement in PI(3,5)$P_2$ binding and introduce a new surface-exposed Cys, we performed extensive mutagenesis of residues predicted to be near its lipid binding site but not directly involved in PI(3,5)$P_2$ binding (**Fig. 1B**). Saturation mutagenesis of C656 (i.e. C656X; X= all amino acids) revealed that C656R had WT-like affinity for PI(3,5)$P_2$ (see **S2B, S2C** and **Table S1**). Also, among various Cys-introducing mutations in the lipid binding site, Y685C had essentially the same affinity for PI(3,5)$P_2$ as WT (see **S2C** and **Table S1**). We thus selected a double mutant, C656R/Y685C as an ideal PI(3,5)$P_2$ sensor for Cys-based chemical conjugation by WCB1. p85α-cSH2 has two additional endogenous cysteins (C659 and C670) but they are buried inside and thus inaccessible to chemical modification unless the protein is fully unfolded. This was confirmed by our finding that C656R was not labeled by WCB1.

**(b) Elimination of protein binding activity**: To ensure that our new sensor selectively responds only to PI(3,5)$P_2$, we eliminated the phosphotyrosine (pY) binding activity of p85α-cSH2 by mutating its pY-binding residue, R649, to Ala. R649A retained the PI(3,5)$P_2$-binding activity of WT (see **Fig. S1D** and **Table S1**). When assayed by fluorescence anisotropy R649A showed drastically (i.e., by 3 orders of magnitude) lower affinity for its cognate pY-containing peptide than p85α-cSH2 WT (**Fig. S1F**). Based on these results, we generated a triple-site mutant, R649A/C656R/Y685C (referred to as *e*p85α-cSH2 hereafter), which had desirable properties for a PI(3,5)$P_2$-specific sensor: i.e., high affinity for PI(3,5)$P_2$ (**Fig. 1E, Fig. 1G** and **Table S1**), greatly suppressed protein-binding affinity (**Fig. S1F** and **Table S1**), and the presence of a single conjugation site.

⇩

**(2) Fluorophore conjugation and elimination of PI(4,5)P2 binding**: Among Cys-specific solvatochromic fluorophores, WCB1 exhibited the most desirable spectral properties as a ratiometric fluorophore when conjugated to *e*p85α-cSH2: i.e., a large spectral shift and a large increase in fluorescence intensity upon lipid binding (see **Fig. 1E, 1F**). Importantly, WCB1-*e*p85α-cSH2 also demonstrated much imrpoved PI(3,5)$P_2$ selectivity over PI(4,5)$P_2$ than p85α-cSH2-WT (**Fig. 1E, 1G**). Thus, WCB-*e*p85α-cSH2 was selected as the ratiometric sensor for PI(3,5)$P_2$.

⇩

**(3) Ratiometric sensor characterization and calibration:** Once PI(3,5)$P_2$ specificity of WCB1-*e*p85α-cSH2 is verified (see **Fig. 1E**) using POPC/POPS/PtdInsP (77/20/3 in mol%) LUVs, then ratiometric calibration curve (see **Fig. 1G**) was obtained by measuring ($F_B/F_G$) as a function of the PI(3,5)$P_2$ concentration in the giant unilamellar vesicles (GUV) under the same conditions in which cellular PI(3,5)$P_2$ quantification is performed.

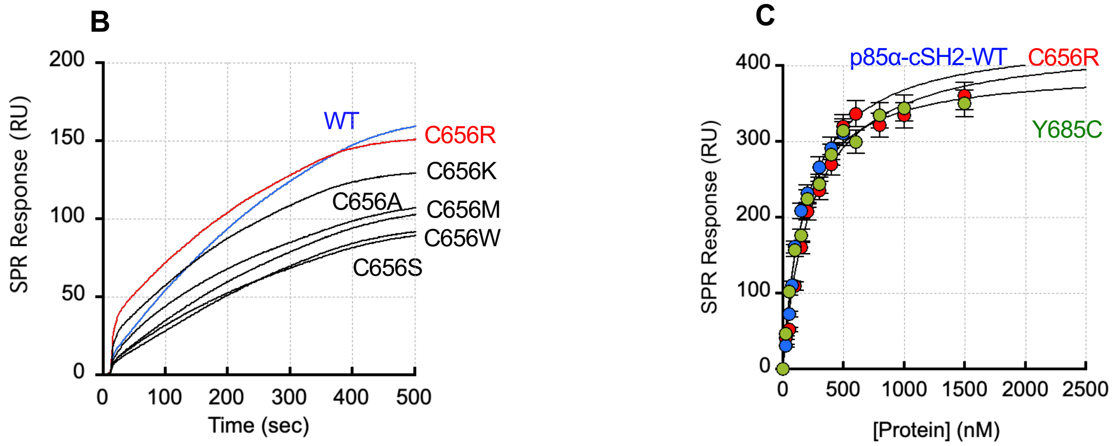


**Fig. S2. Development and characterization of a ratiometric PI(3,5)$P_2$ sensor (WCB1-ep85α-cSH2)**
**A.** Steps of a ratiometric PI(3,5)$P_2$ sensor development and cellular ratiometric imaging.
**B.** Relative affinity of p85α-cSH2 WT and selected C656X mutants for POPC/POPS/PI(3,5)$P_2$ (77:20:3) vesicles determined by SPR analysis. The protein concentration was 300 nM. **C.** Determination of $K_d$ for binding of p85α/β-cSH2 WT and mutants to POPC/POPS/PI(3,5)$P_2$ (77:20:3) LUVs as described in **Fig. S1D**. Total protein concentrations were 300 nM.

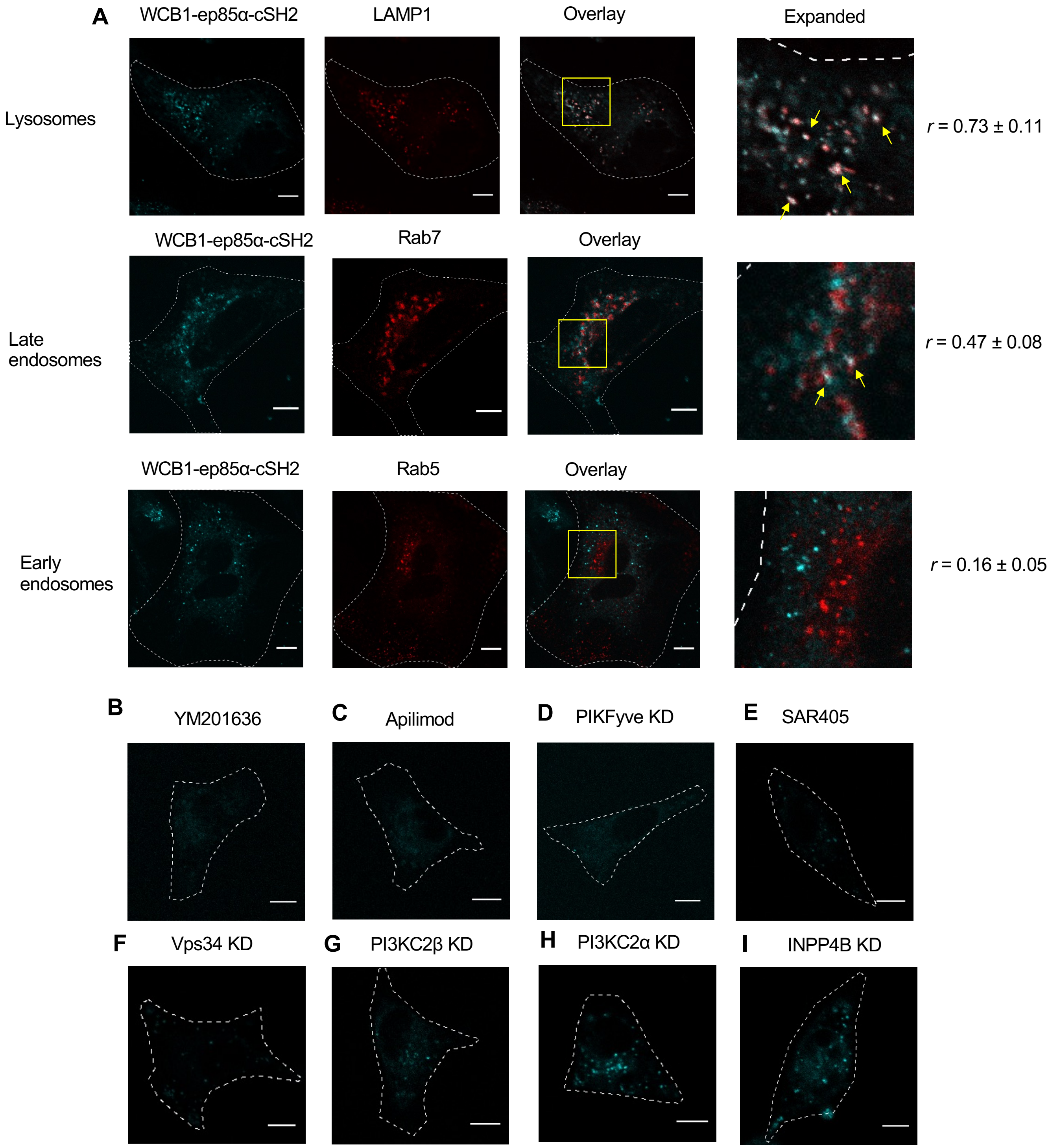


**Fig. S3. Effects of lipid kinases and phosphatases on subcellular localization of PI(3,5)$P_2$ detected by WCB1-ep85α-cSH2 in HeLa cells**. **A.** Colocalization of microinjected WCB-ep85α-cSH2 with organelle markers, including mCherry-LAMP1 (lysosomes), iRFP-Rab7 (late endosomes), and iRFP-Rab5 (early endosomes). The right panels show expanded views of colocalized regions (yellow boxes). Yellow arrows indicate representative colocalized spots (in white color). *r* values indicate Pearson's correlation coefficients. **B.-G.** Effects of PIKfyve inhibition by YM201636 (**B,** 0.8 μM, 1 h), apilimod (**C,** 1 μM, 1 h), PIKfyve knockdown (KD) (**D**), Vps34 inhibition by SAR405 (**E,** 10 μM, 24 h) and VPS34 KD (**F**), PI3KC2β KD (**G**), PI3KC2α KD (**H**), and INPP4B KD (**I**) on subcellular localization of WCB-ep85α-cSH2. The images are representatives of the majority of cells (*n* > 50 cells in 5 independent measurements) treated under the same conditions. Cell images were shown only for larger HeLa cells for better illustration but HeLa and HEK293 cells have very similar localization patterns. Scale bars indicate 10 μm.

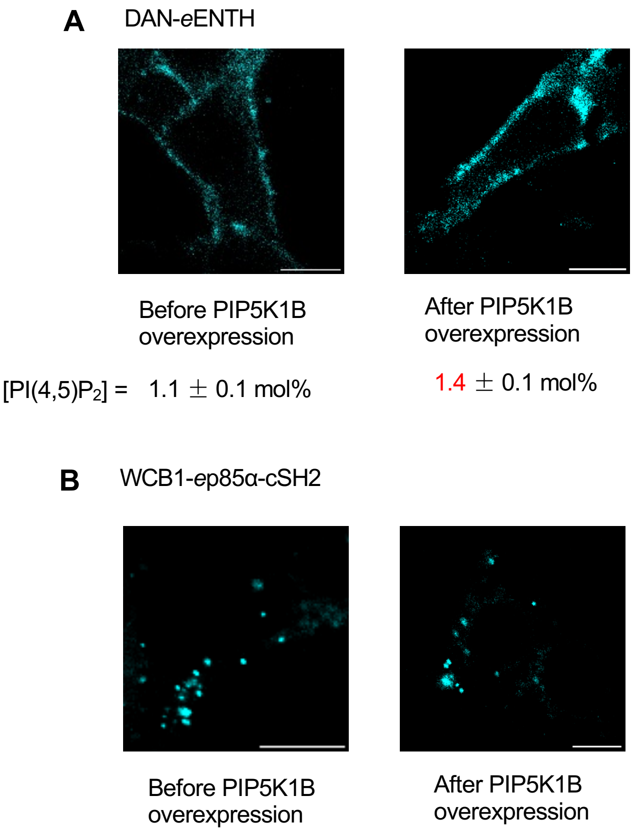


**Fig. S4. Effects of PI(4,5)$P_2$ changes in the PM on subcellular localization of WCB1-ep85α-cSH2 in HEK293 cells**. **A.** Detection and quantification of PI(4,5)$P_2$ in the PM of HEK293 cells before and after PIP5K1B overexpression. A ratiometric PI(4,5)$P_2$ sensor, DAN-*e*ENTH was microinjected into the cells for PI(4,5)$P_2$ quantification. Spatially averaged PI(4,5)$P_2$ concentrations were 1.1 ± 0.1 mol% (before PIP5K1B overexpression) and 1.4 ± 0.1 mol% (after PIP5K1B overexpression). **B.** Lack of responses to PI(4,5)$P_2$ changes in the PM by WCB1-*e*p85α-cSH2. WCB1-*e*p85α-cSH2 showed no detectable PM localization before and after PIP5K1B overexpression. Each image is representative of 10 images with similar patterns. Scale bars indicate 10 μm.

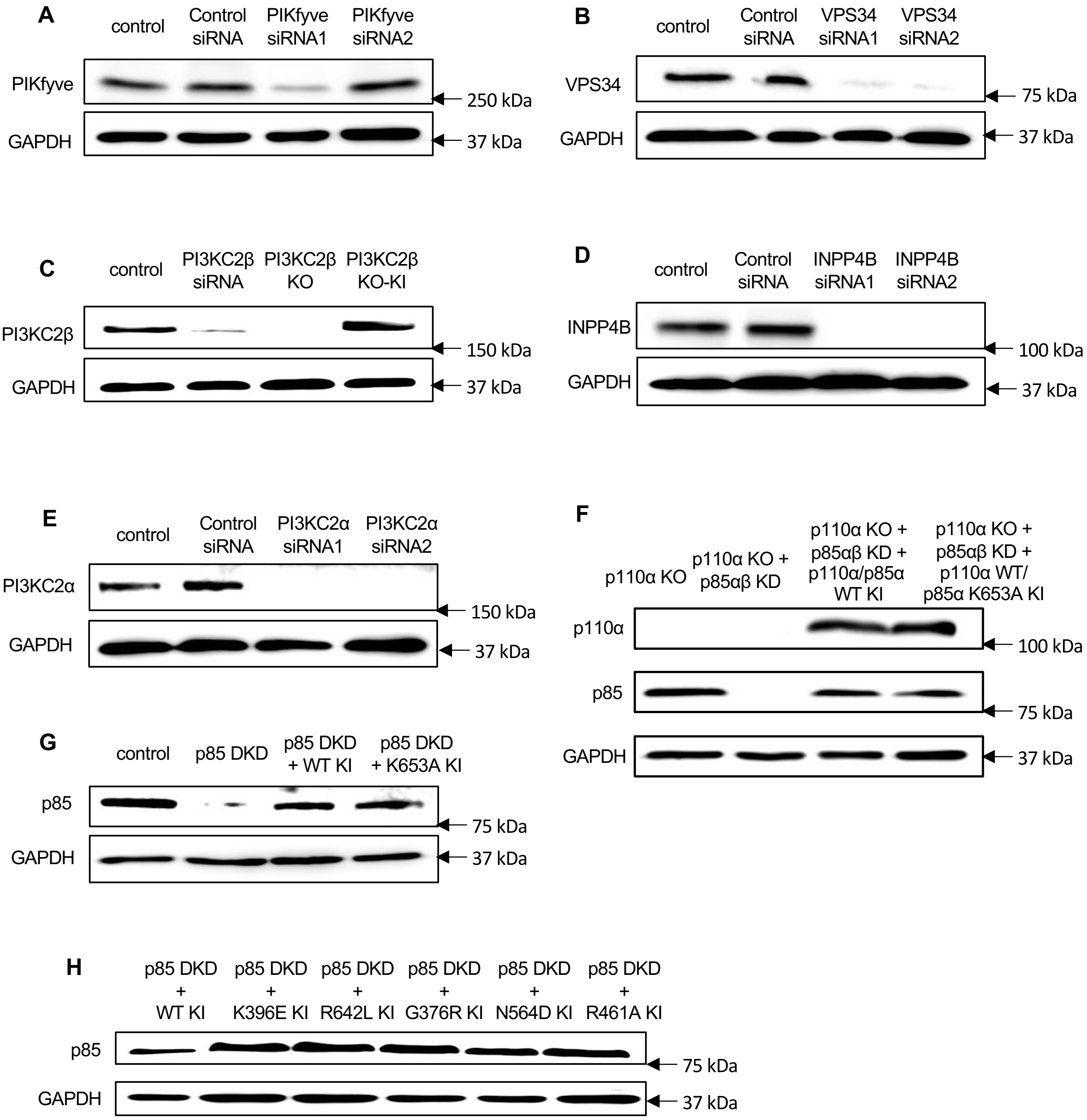


**Fig. S5. Western blot analysis of HEK 293 cells after genetic modulation of lipid kinases and phosphatases.**

Western blots confirmed efficient suppression of PIKfyve (**A**), VPS34 (**B**), PI3KC2β (**C**), INPP4B (**D**), PI3KC2α (**E**), p85 (**F-G**), by siRNA and efficient add-back of indicated genes (**C, F, G, H**). Non-targeting siRNAs were used as negative controls and GAPDH as loading controls. From these experiments, the most effective siRNA for each enzyme was selected for subsequent knockdown experiments. Western blots confirmed successful gene ablation of PI3KC2β (**C**) and p110α (**F**). KO, KD, DKD, and KI indicate knockout, knockdown, double-knockdown, and knock-in, respectively.

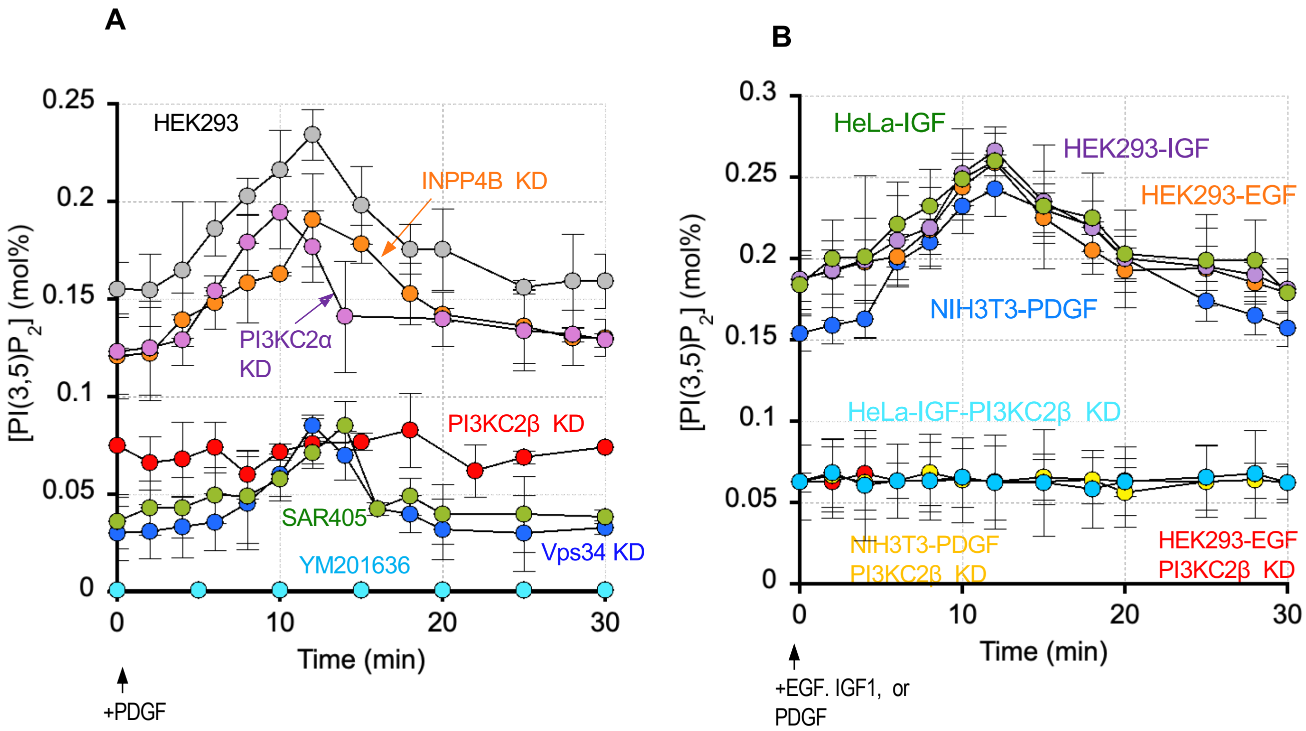


**Fig. S6. Time courses of the PDGF-stimulation changes in [PI(3,5)$P_2$] under different conditions. A.** Kinetics of PDGF-stimulated [PI(3,5)$P_2$] changes in HEK293 cells after cells were pre-treated with INPP4B siRNA (KD) (orange), PI3KC2α siRNA (light purple), PI3KC2β siRNA (red), a Vps34 inhibitor SAR405 (green, 10 μM, 24 h), Vps34 siRNA (blue), and a PIKfyve inhibitor, YM201636 (cyan, 0.8 μM, 1 h). PIKFyve knockdown yielded the same result as YM201636 treatment. The control HEK293 cell curve is shown in gray. **B.** Kinetics of [PI(3,5)$P_2$] changes after HEK293 cells were stimulated with EGF (orange) and IGF1 (light purple), NIH3T3 cells were stimulated with PDGF (blue), and HeLa cells were stimulated with IGF1 (green). Also, the kinetics of [PI(3,5)$P_2$] changes are shown after EGF stimulation of HEK293 cells (red), PDGF stimulation of NIH 3T3 cells (yellow), and IGF1 stimulation of HeLa cells (cyan), all pre-treated with PI3KC2β siRNA. Each [PI(3,5)$P_2$] value represents an average and S.D at a given time point ($n$ = 3 with 5 cells per experiment). 50 ng/ml growth factors were used.

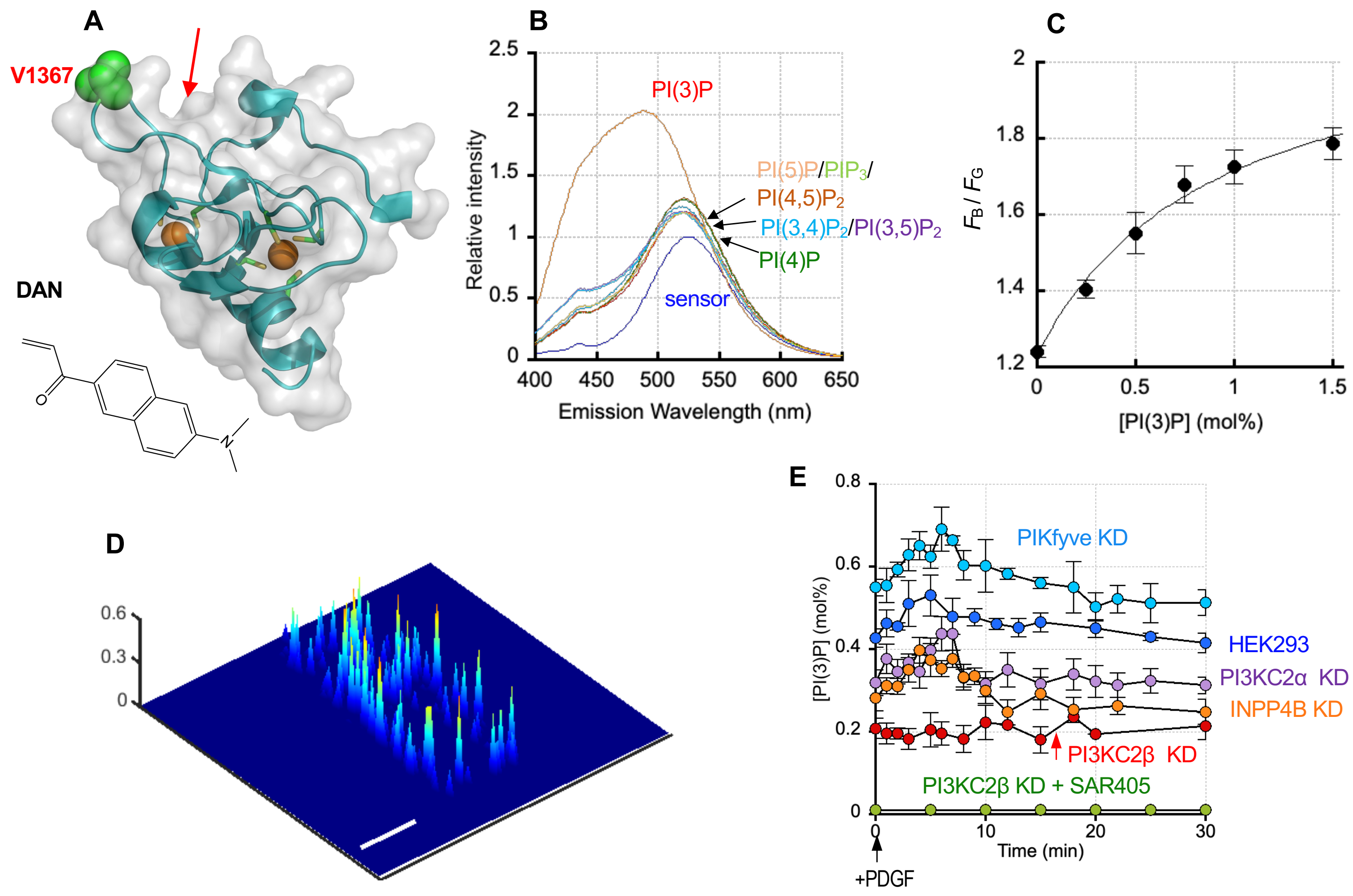


**Fig. S7. Protein engineering of a ratiometric PI(3)P sensor, DAN-*e*EEA1, and *in situ* quantification of cellular PI(3)P by DAN-*e*EEA1.**

**A.** Protein engineering of DAN-eEEA1. The structure of EEA1 (protein data bank ID = 1JOC) shown in a ribbon diagram with its surface shown in light gray. The molecule is oriented with its potential membrane binding surface facing upward. The PI(3)P binding site is indicated by the red arrow. Although the EEA1 FYVE domain contains 8 cysteines (stick representation), they are not accessible for fluorophore labeling because they are involved in $Zn^{2+}$ (gold globes) coordination. We thus introduced an additional cysteine on its membrane binding surface by V1367C mutation (green) and labeled this engineered protein (*e*EEA1) with a solvatochromic fluorophore, acrylodan, to yield DAN-*e*EEA1. **B.** PtdInsP selectivity of DAN-*e*EEA1. Fluorescence emission spectra of DAN-*e*EEA1 (500 nM) in response to binding to 50 μM POPC/POPS/PtdInsP (77/20/3 in mol%) LUVs were obtained spectrofluorometrically with the excitation wavelength set at 380 nm. DAN-*e*EEA1 showed a strong solvatochromic shift upon binding to PI(3)P-containing vesicles, which was not observed with vesicles containing other PtdInsPs, demonstrating its PI(3)P specificity. The spectra are representative of three similar independent data. **C.** The ratometric calibration curve of DAN-*e*EEA1 determined by fluorescence microscopy. Lipid compositions of GUVs were POPC/POPS/PI(3)P (80-x/20/x: $x$ = 0-2 mol%). The binding curve was analyzed by non-linear least squares analysis using a modified Langmuir equation: $y = y_{min} + (y_{max} - y_{min})/(1 + K_d / [PI(3)P])$ where $K_d$, $y_{max}$ and $y_{min}$ are [PI(3)P] yielding half maximal binding, the maximal and minimal $y$ values. Half maximal vesicle binding was achieved with 0.90 ± 0.28 mol% [PI(3)P]. Each data represents an average ± S.D. from three independent determinations. **D.** A spatially resolved PI(3)P profile calculated from the two-channel cross-sectional ratiometric images of a representative unstimulated HEK293 cell ($n$ =3 with 10 cells per experiment) at a given time. The z-axis scale indicates the PI(3)P concentration ([PI(3)P]) (mol%). A pseudo-coloring scheme with red and blue representing the highest (1 mol%) and the lowest (0 mol%) concentration, respectively, is used to illustrate the spatial PI(3)P heterogeneity. Scale bar, 10 μm. **E.** Kinetics of PDGF-stimulated [PI(3)P] changes in HEK293 cells before (blue) and after cells were pre-treated with INPP4B siRNA (KD) (orange), PI3KC2α siRNA (purple), PI3KC2β siRNA (red), PIKfyve siRNA (cyan), and a combination of PI3KC2β siRNA and a Vps34 inhibitor SAR405 (green, 10 μM, 24 h). [PI(3)P] values (average ± S.D. from three independent determinations with 10 cells per experiment) were determined at different time intervals. 50 ng/ml PDGF was used.

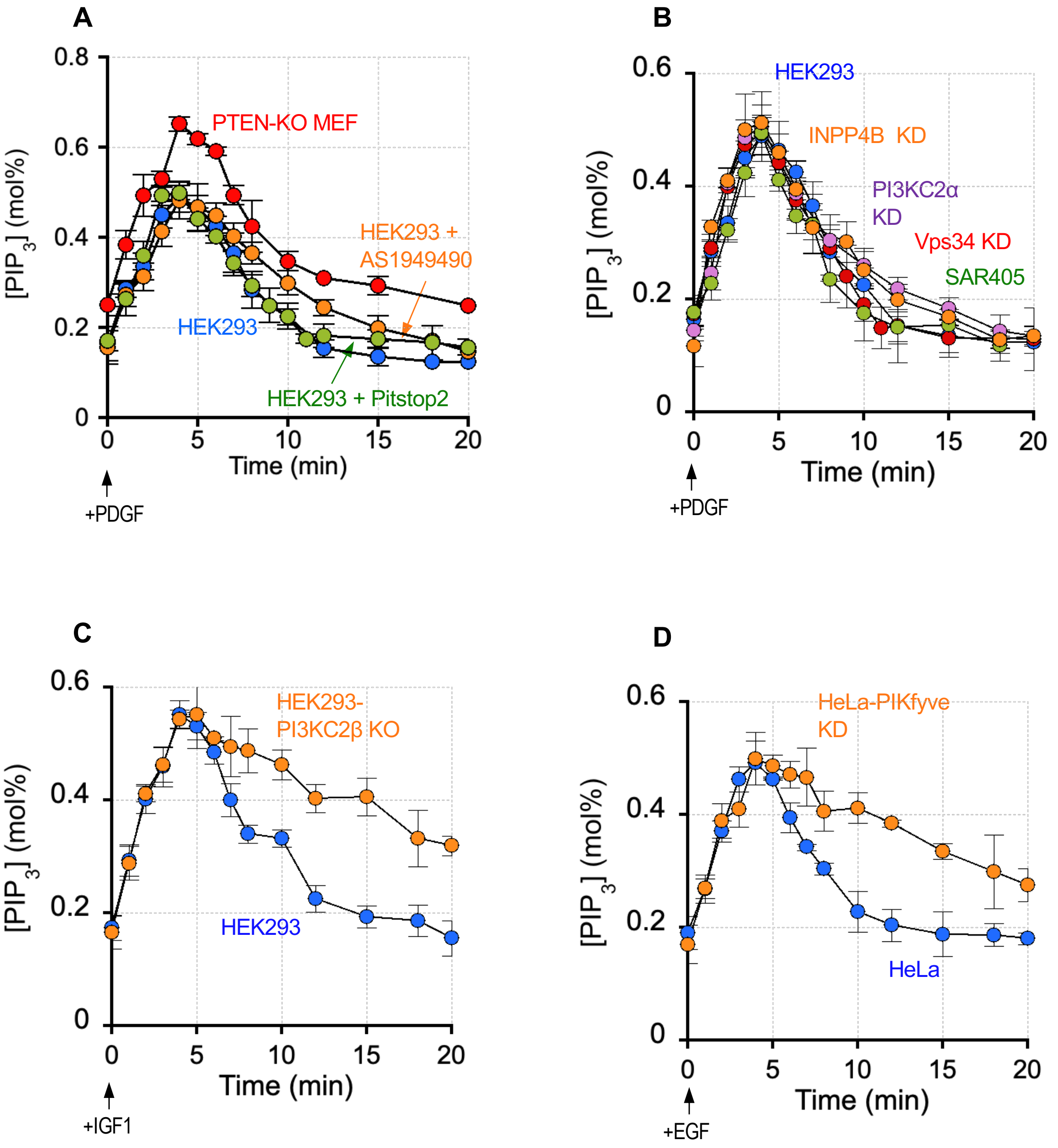


**Fig. S8. Effects of lipid kinase and phosphatase inhibition on the kinetics of PDGF-induced $PIP_3$ changes**

**A.** PDGF-stimulated changes in [$PIP_3$] at the PM of HEK293 cells before (blue) and after (orange) SHIP2 inhibition (20 μM AS1949490 for 2 h). The same measurement was performed after Pitstop2 treatment (green; 30 μM for 15 min) in HEK293 cells. It was also repeated in PTEN-null (KO) MEF cells (red). **B.** PDGF-stimulated changes in [$PIP_3$] at the PM of HEK293 cells before (blue) and after cells were pre-treated with INPP4B siRNA (KD) (orange), PI3KC2α siRNA (purple), a Vps34 inhibitor 10 μM SAR405 (green, 24h), and Vps34 siRNA (red). **C**. IGF1-stimulated changes in [$PIP_3$] at the PM of WT (blue) and PI3KC2β-null (orange) HEK293 cells. **D**. EGF-stimulated changes in [$PIP_3$] at the PM of HeLa cells before (cyan) and after (orange) PIKfyve knockdown. 50 ng/ml growth factors were used. [$PIP_3$] values (average ± S.D. from three independent determinations with 10 cells per experiment) were determined at different time intervals.

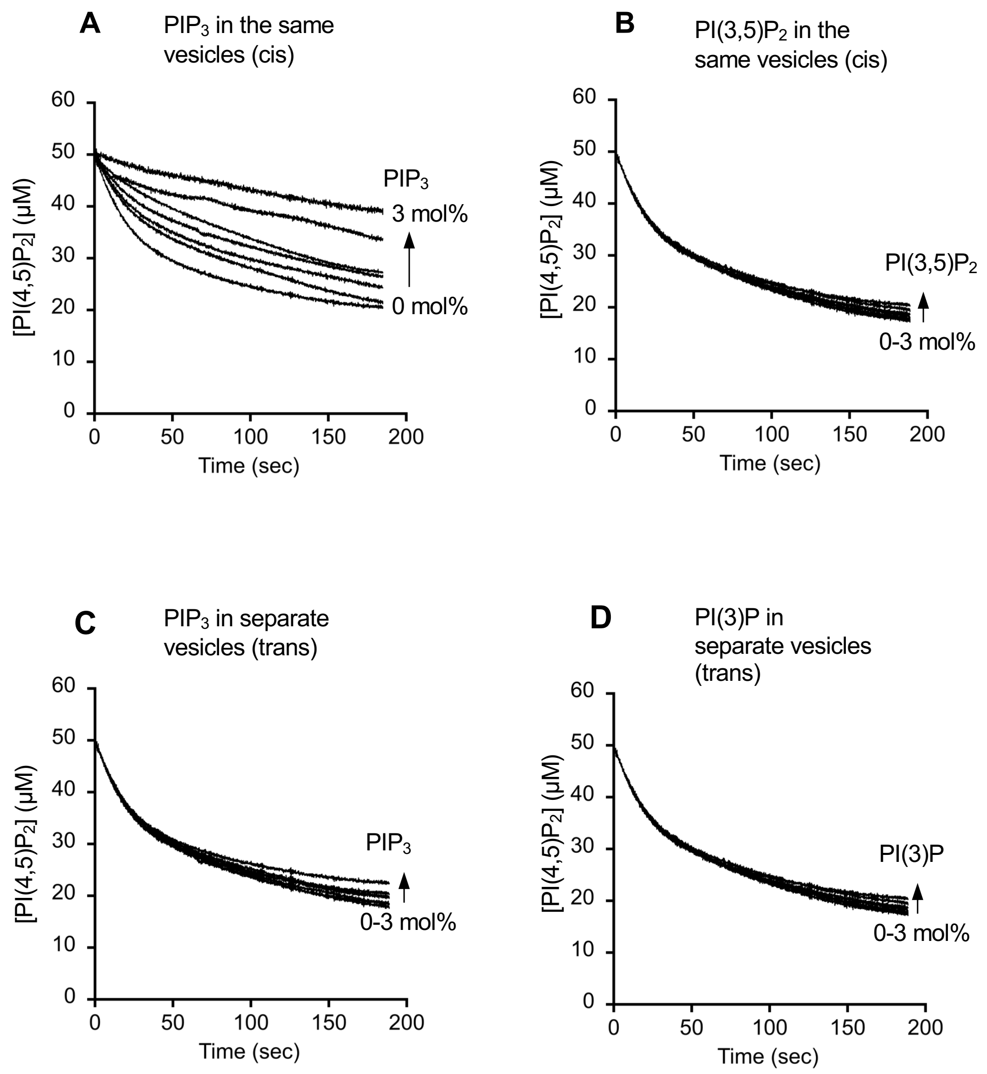


**Fig. S9. Effects of PtdInsP on the enzyme activity of PI3Kα**
**A.** An inhibitory effect of $PIP_3$ on PI3Kα acting on PI(4,5)$P_2$ in the same vesicle was measured using POPC/POPS/PI(4,5)$P_2$/$PIP_3$ (77-*x*/20/3/*x*; *x* = 0-3 mole%)) LUVs. **B.** Under the same conditions, PI(3,5)$P_2$ showed no inhibition (i.e., in POPC/POPS/ PI(4,5)$P_2$/PI(3,4)$P_2$ (77-*x*/20/3/*x*; *x* = 0-3 mole%) LUVs). **C.-D.** An inhibitory effect of $PIP_3$ (**C**) and PI(3)P (**D**) on PI3Kα acting on PI(4,5)$P_2$ in separate vesicles was measured using POPC/POPS/PI(4,5)$P_2$ (77:20:3) and POPC/POPS/$PIP_3$ (or PI(3)P) (77-*x*/20/*x*; *x* = 0-3 mole%)) LUVs. See **Fig. 4B** for experimental conditions.

**A** **PI3Kα on PM (TIFRM)**

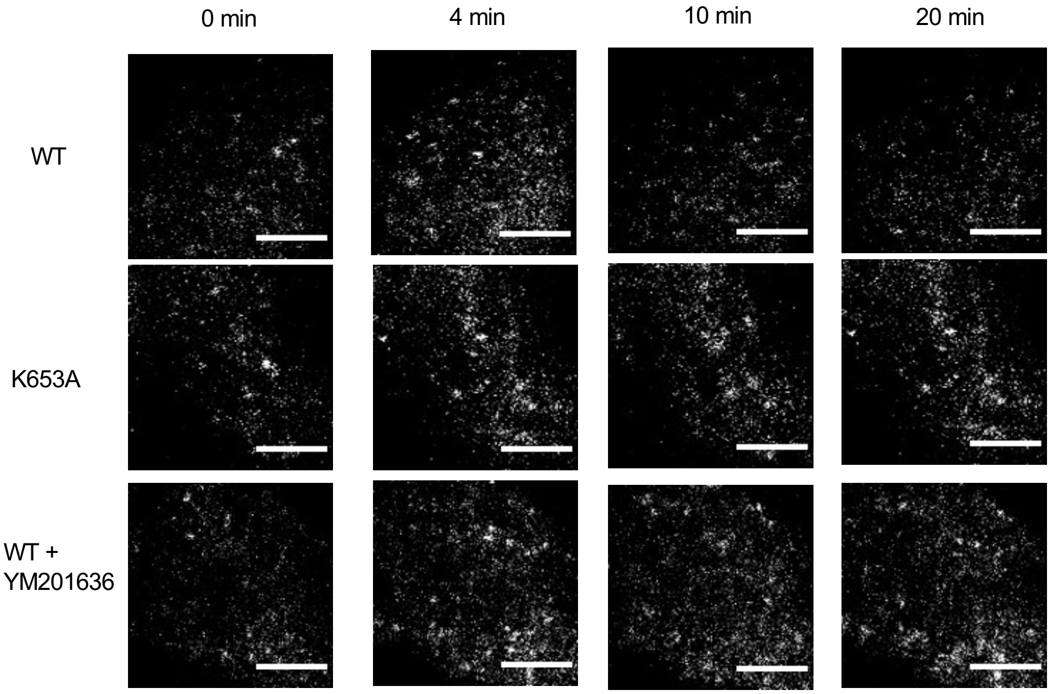


**B** **PI3K on PM (Confocal microscopy)**

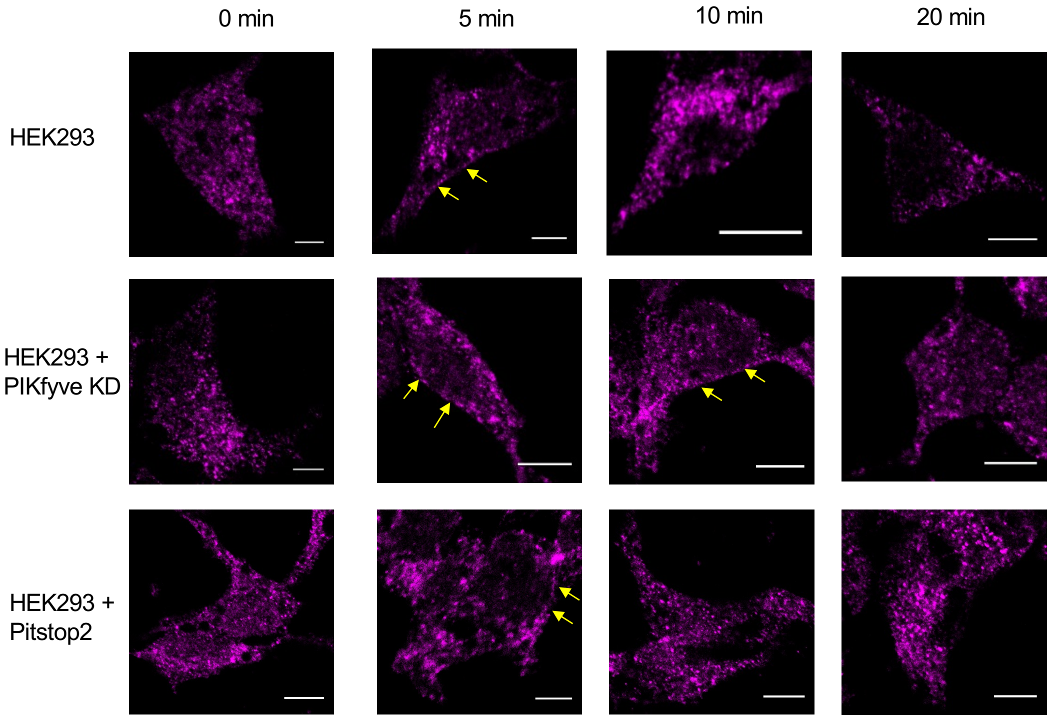


***Fig. S10***

C PI3K on lysosomes (confocal microscopy)

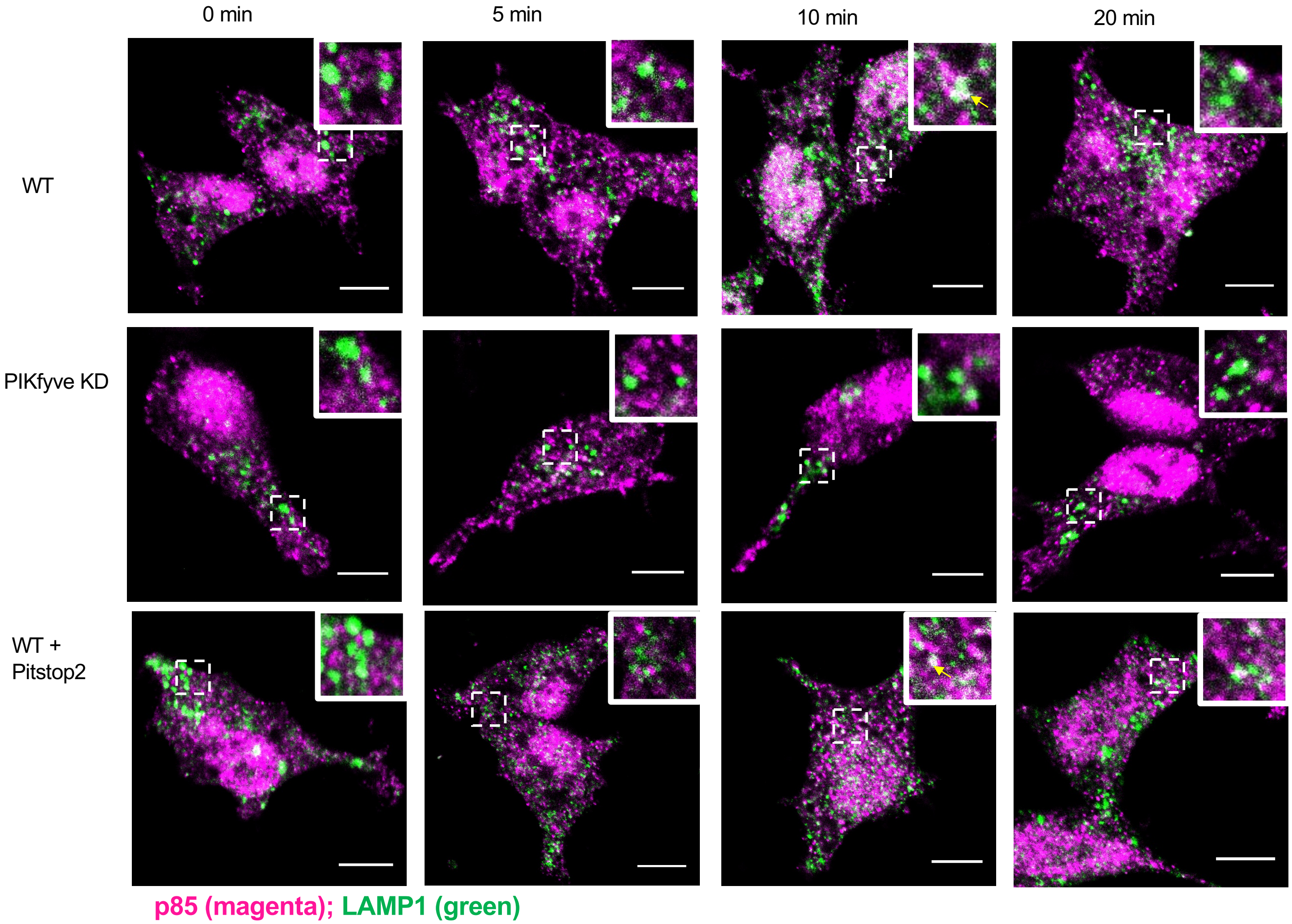


D PI3K on lysosomes (Expansion microscopy)

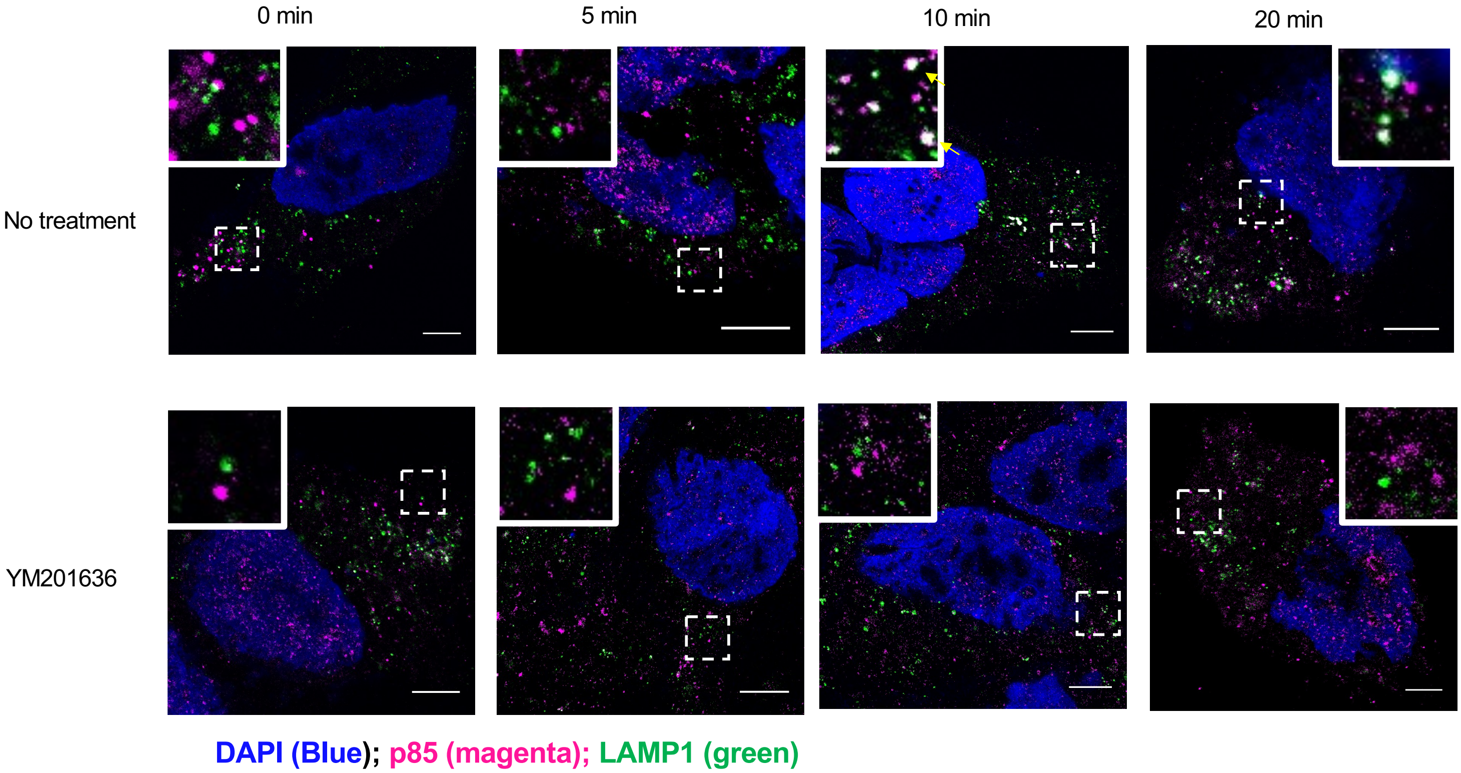


*Fig. S10*

**E PI3K on early endosomes (confocal microscopy)**

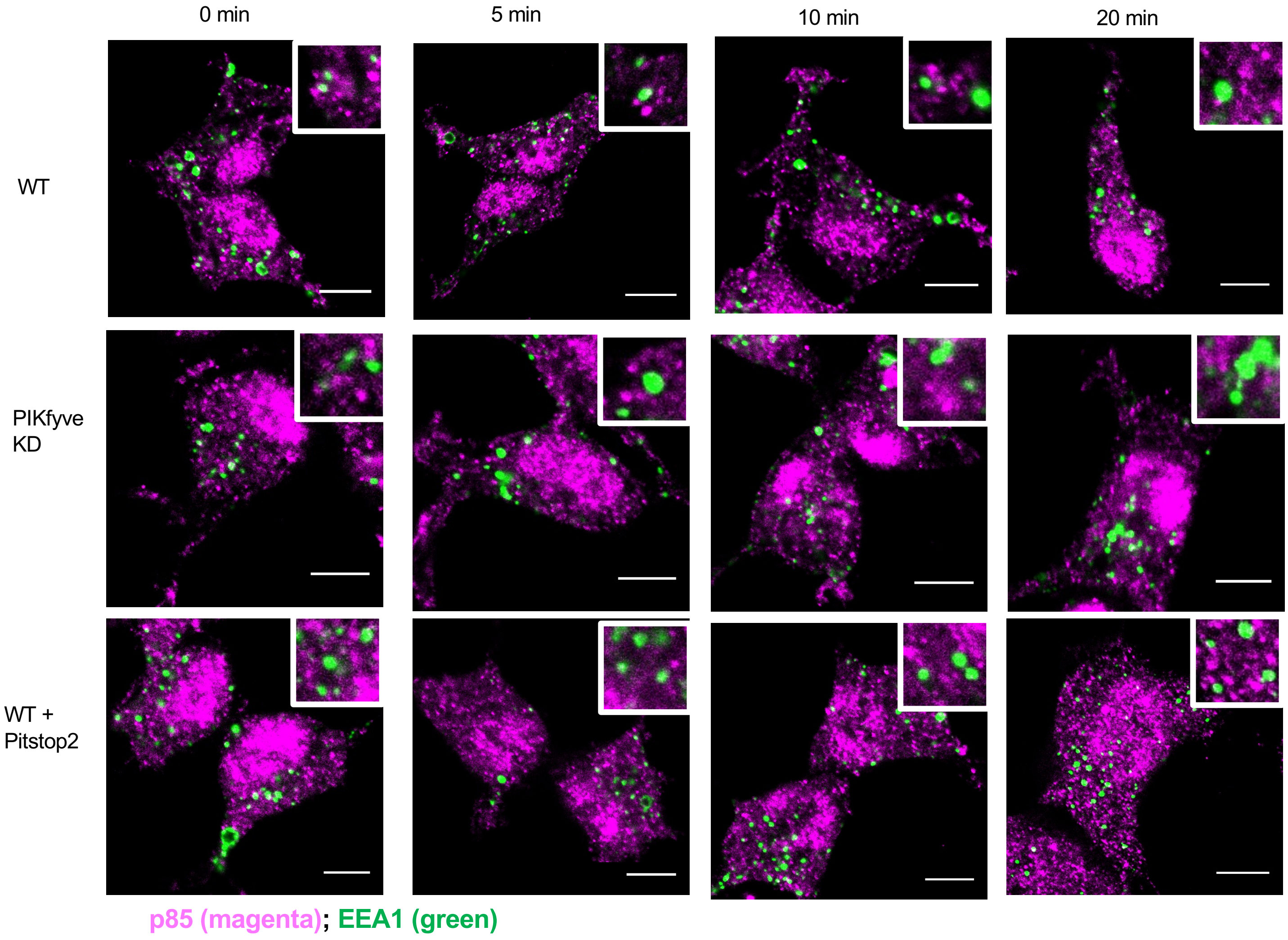


**Fig. S10. Time courses of subcellular localization of PI3K after PDGF stimulation of HEK293 cells**
**A.** Total internal reflection fluorescence microscopy (TIRFM) images of EGFP-PI3Kα-WT and EGFP-PI3Kα-K653A on the plasma membrane (PM) of PDGF (50 ng/ml)-stimulated HEK293 cells. EGFP-PI3Kα-WT was also pre-treated with 0.8 μM YM201636 for 1 h. **B.** Confocal microscopy images of endogenous PI3K (stained with the p110α antibody) in PDGF-stimulated HEK293 cells (untreated, pre-treated with PIKFyve siRNA (KD) and 30 μM Pitstop2 for 15 min). Yellow arrows indicate PM localization of PI3Kα. **C.** Confocal images of colocalization of endogenous PI3K and LAMP1 (stained with the p85 (magenta) and LAMP1 (green) antibodies) in PDGF-stimulated HEK293 cells (untreated, PIKFyve KD and, treated with Pitstop2). Insets show the magnified images of the lysosomes (indicated by dashed boxes). **D.** Expansion microscopy (ExM) images of colocalization of endogenous PI3K and LAMP1 (stained with the p85 (magenta) and LAMP1 (green) antibodies) in PDGF-stimulated HEK293 cells (untreated, PIKFyve KD and, treated with Pitstop2). Insets show the magnified images of the lysosomes (indicated by dashed boxes). Nuclei were stained with DAPI (blue). **E.** Confocal images of colocalization of endogenous PI3K and EEA1 (stained with the p85 (magenta) and EEA1 (green) antibodies) in PDGF-stimulated HEK293 cells (untreated, PIKfyve KD and, treated with Pitstop2). Insets show the magnified images of the early endosomes. Each image is representative of similar images collected from three independent measurements with >10 cells per measurement. White spots (and yellow arrows at 10 min) indicate co-localization. Scale bars for all but ExM images indicate 10 μm. For ExM images, Scale bars are 6.25 μm (25 μm at the post-expansion state).

**A** **p110α on lysosomes**

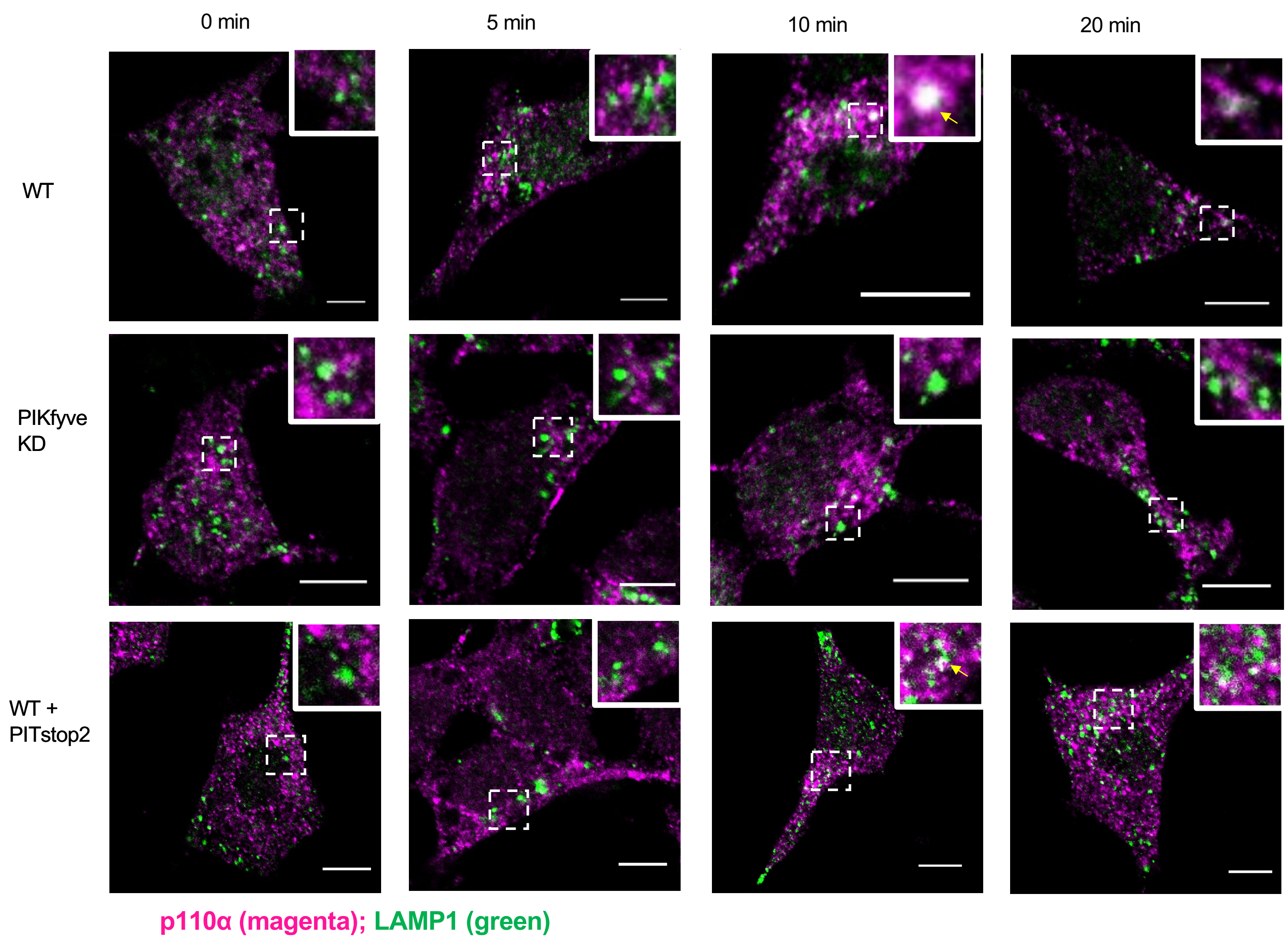


*Fig. S11*

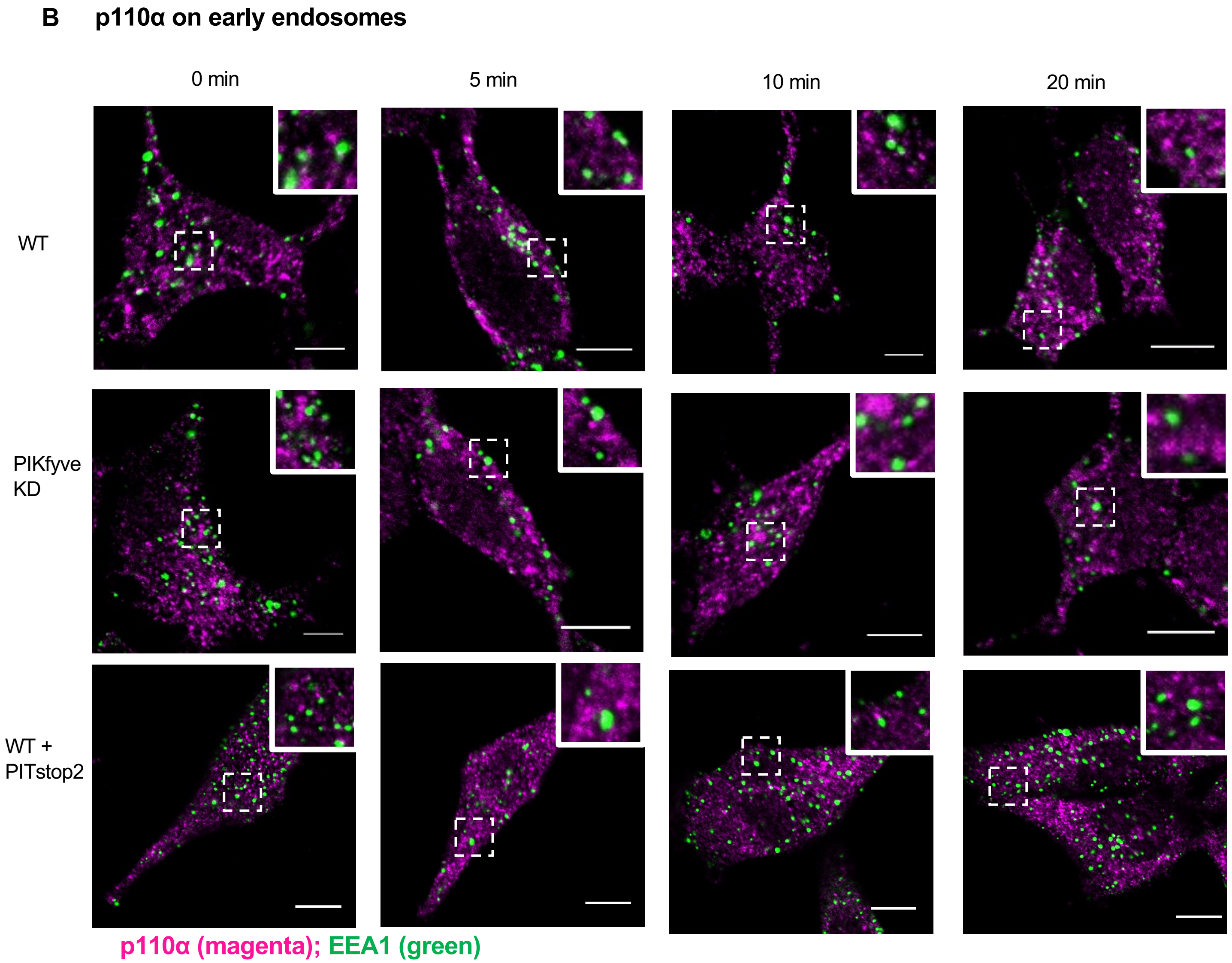


**Fig. S11. Time courses of subcellular localization of p110α after PDGF stimulation of HEK293 cells**
**A.** Confocal images of colocalization of endogenous p110α and LAMP1 (stained with the p110α (magenta) and LAMP1 (green) antibodies) in PDGF (50 ng/ml)-stimulated HEK293 cells (untreated, PIKFyve KD and, treated with Pitstop2 (30 μM for 15 min)). Insets show the magnified images of the lysosomes (indicated by dashed boxes). White spots indicate co-localization. **B.** Confocal images of colocalization of endogenous p110α and EEA1 (stained with the p110α (magenta) and EEA1 (green) antibodies) in PDGF-stimulated HEK293 cells (untreated, PIKfyve KD and, treated with Pitstop2). Insets show the magnified images of the early endosomes (indicated by dashed boxes). White spots indicate co-localization. Each image is representative of similar images collected from three independent measurements with >10 cells per measurement. White spots (and yellow arrows at 10 min) indicate co-localization. Scale bars indicate 10 μm

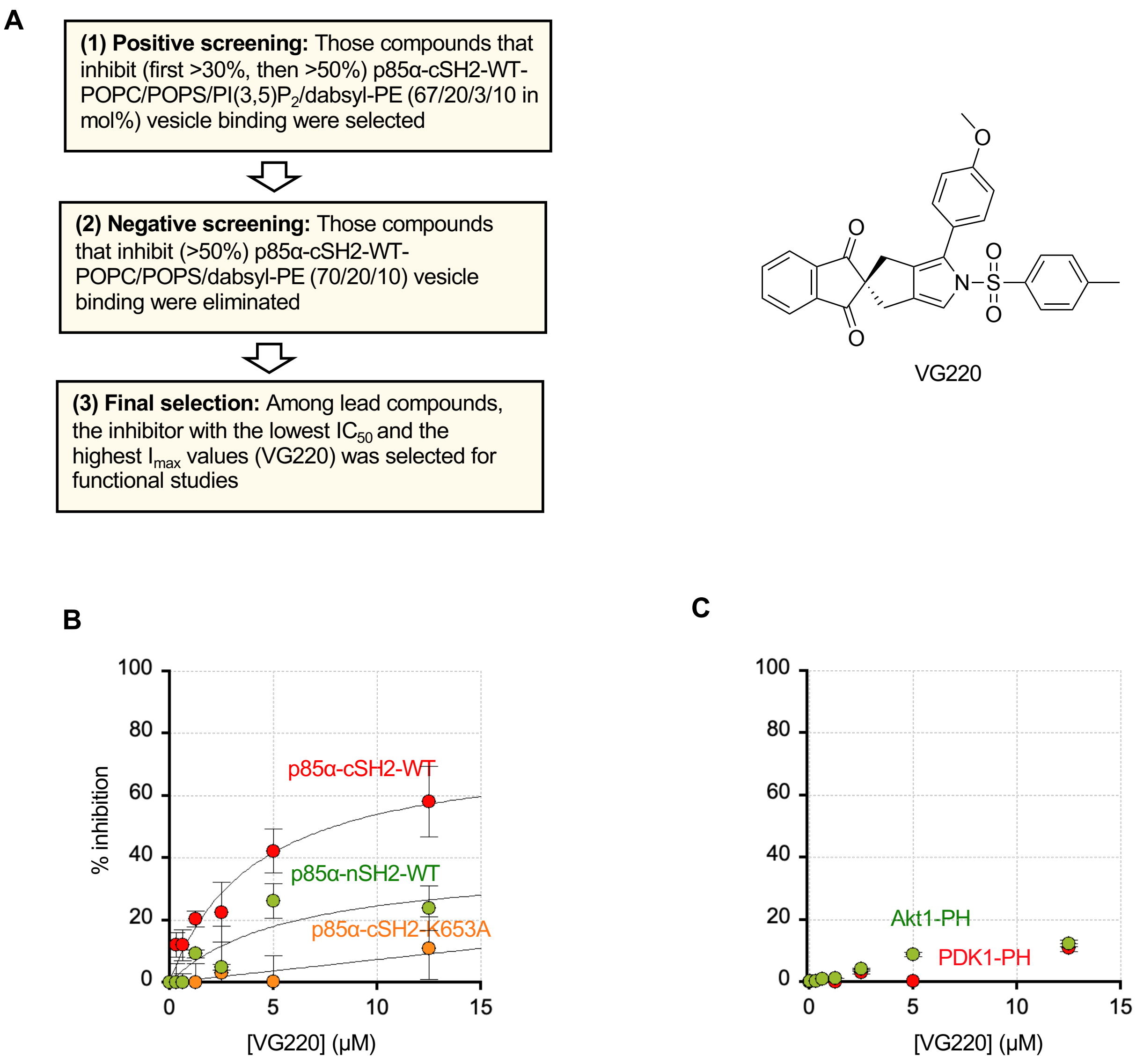


**Fig. S12. Development and characterization of a small molecule inhibitor of p85-cSH2-PI(3,5)$P_2$ interaction. A.** A general scheme of inhibitor screening and the structure of VG220. The best inhibitor, VG220, was selected after three (two positive and one negative) rounds of screening and detailed characterization of lead compounds. **B.** Dose-dependent inhibition of SH2 domain-PI(3,5)$P_2$ interaction by VG220**.** p85α-cSH2-WT, p85α-cSH2-K653A, and p85α-nSH2-WT were employed to demonstrate high specificity of VG220. **C**. Dose-dependent inhibition of PH domain-$PIP_3$ interaction by VG220. POPC/POPS/PI(3,5)$P_2$ (or $PIP_3$)/dabsyl-PE (67/20/3/10 in mol%) vesicles were used. Total protein and lipid concentrations were 100 nM and 50 μM respectively in 20 mM Tris buffer (pH 7.4) containing 0.16 M NaCl. $I_{max}$ and $IC_{50}$ values of inhibitors were determined using the equation, $I = I_{max} / (1 + IC_{50} / [I])$ (I: % inhibition, [I]: VG220 concentration). Data represent average and SD values calculated from three independent measurements.

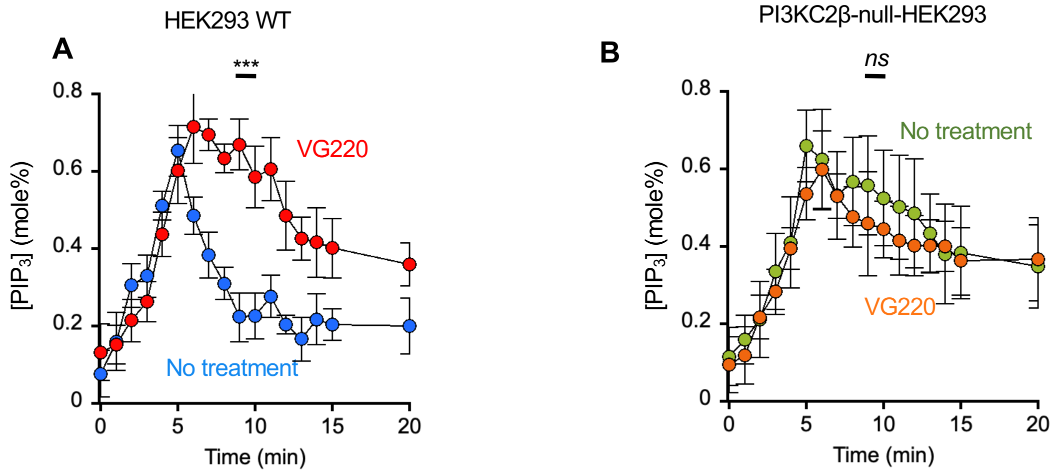


**Fig. S13. Effects of VG220 on the $PIP_3$ formation in WT(A) and PI3KC2β-null (B) HEK cells**
See Fig. 6B-C for experimental conditions. P values were calculated at 10 min post PDGF stimulation. $p$ = <0.0001 (before and after VG220 treatment in HEK293 cells) and 0.543 (before and after VG220 treatment in PI3KC2β-null-HEK293 cells).

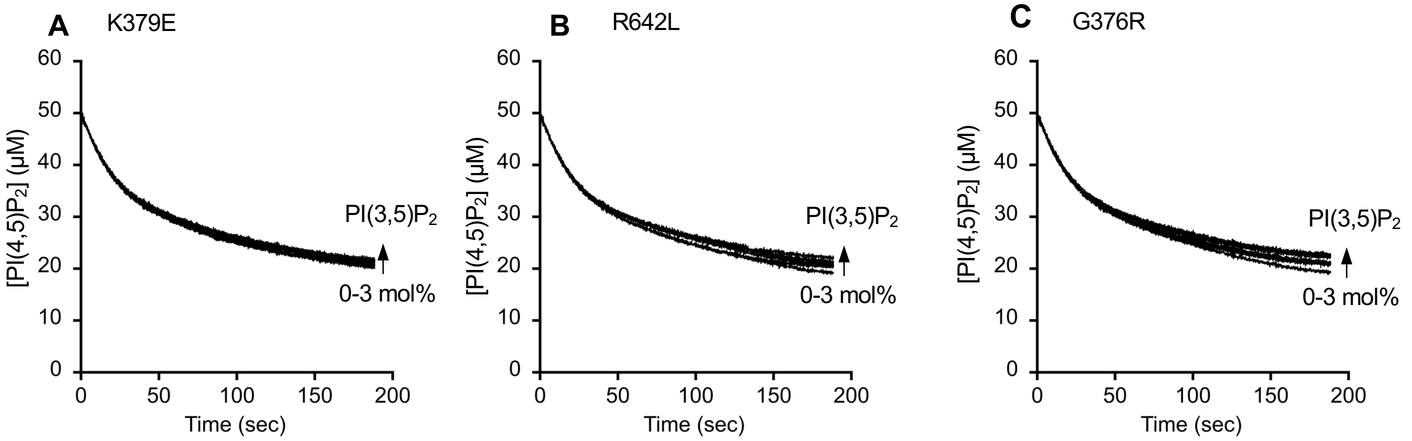


**Fig. S14. Inhibition of the enzymatic activity of PI3Kα-p85α mutants by PI(3,5)$P_2$**

Inhibitory effects of PI(3,5)$P_2$ on PI3Kα mutants, K379E (**A**), R642L (**B**), and G376R (**C**) acting on PI(4,5)$P_2$ in separate vesicles (i.e., in a *trans* mode) was measured by a cell-free kinetic assay using POPC/POPS/PI(4,5)$P_2$ (77:20:3) and POPC/POPS/$PIP_3$ (or PI(3)P) (77-*x*/20/*x*; *x* = 0-3 mole%)) LUVs. See **Fig. 4B** for experimental conditions.

### SUPPLEMENTARY TABLES

**Table S1. Lipid and peptide binding properties of p85-SH2 domains and mutants**

| Proteins | $K_d$ (nM) for POPC/POPS/X (77:20:3) LUVs [a] | | $K_d$ (μM) for pY peptide [b] |
|---|---|---|---|
| | X= PI(3,5)$P_2$ | X= PI(4,5)$P_2$ | |
| p85α-cSH2 | 195 ± 25 | 960 ± 90 | 0.4 ± 0.1 |
| p85β-cSH2 | 560 ± 80 | ND[c] | ND |
| p85α-cSH2-K653A | 2300 ± 400 | ND | 0.2 ± 0.1 |
| p85α-cSH2-R649A | 240 ± 30 | ND | 40 ± 5 |
| p85α-cSH2-C656R | 240 ± 35 | ND | ND |
| p85α-cSH2-Y685C | 170 ± 20 | ND | ND |
| *e*p85α-cSH2 | 200 ± 20 | ND | 43 ± 4 |

[a] Average ± S.D. values determined from SPR analysis (*n* = 3); see Extended Data Fig. 1b and 2d for binding isotherms and experimental details

[b] Average ± S.D. values determined from fluorescence anisotropy analysis (*n* = 3); see Extended Data Fig. 1f for binding isotherms and experimental details.

[c] Not determined
